# Behavioral deficits, learning impairment, and enhanced hippocampal excitability in co-isogenic *Prnp^ZH3/ZH3^* mice

**DOI:** 10.1101/2021.02.20.432083

**Authors:** A. Matamoros-Angles, A. Hervera, J. Soriano, E. Martí, P Carulla, F. Llorens, M Nuvolone, A. Aguzzi, I. Ferrer, A. Gruart, JM. Delgado-García, JA. Del Río

## Abstract

**Background:** The cellular prion protein (PrP^C^) has been associated with numerous cellular processes, such as cell differentiation and neurotransmission. Moreover, it was recently demonstrated that some functions were misattributed to PrP^C^ since results were obtained from mouse models with genetic artifacts. Here we elucidate the role of PrP^C^ in the hippocampal circuitry and its related functions, like learning and memory, using the new strictly co-isogenic *Prnp^0/0^* mouse *(Prnp^ZH3/ZH3^)*. Behavioral and operant conditioning tests were performed to evaluate memory and learning capabilities. *In vivo* electrophysiological recordings were carried out at CA3-CA1 synapses in living behaving mice, and spontaneous neuronal firing and network formation were monitored in primary neuronal cultures of *Prnp^ZH3/ZH3^ vs*. wild-type mice.

**Results:** Results showed decreased motility, impaired operant conditioning learning, and anxiety-related behavior in *Prnp^ZH3/ZH3^* animals. PrP^C^ absence enhanced susceptibility to high-intensity stimulations and kainate-induced seizures. However, long-term potentiation (LTP) was not enhanced in the *Prnp^ZH3/ZH3^* hippocampus. In addition, we observed a delay in neuronal maturation and network formation in *Prnp^ZH3/ZH3^* cultures.

**Conclusion:** In conclusion, PrP^C^ mediates synaptic function and protects the synapse from excitotoxic insults. Its deletion might evoke a susceptible epileptogenic brain that would fail to perform highly cognitive-demanding tasks such as associative learning and anxiety-like behaviors.

## Background

Cellular prion protein (PrP^C^) is a cell surface Glycosylphosphatidylinositol (GPI) anchored protein expressed in several tissues with high levels in the nervous system. It is expressed ubiquitously in the brain, especially in neurons and glial cells [1–5]. PrP^C^ is known for its crucial role in the pathogenesis of human and animal prionopathies [6, 7]. In these diseases, PrP^C^ is transformed into a misfolded β-sheet-rich isoform, the infectious prion protein (PrP^Sc^) [6]. Increasing knowledge about the participation of PrP^C^ in prion pathogenesis contrasts with puzzling data regarding its natural physiological role/s [8–10]. Indeed, this controversy was also strengthened by the absence, until a few years ago, of an appropriate *Prnp^0/0^* mouse model without PrP^C^ protein, with high breeding capability to dissect biological relevance in specific processes [9, 11–13].

PrP^C^ has been previously described as neuroprotective, mainly by using loss-of-function approaches [14–16], while in other studies, PrP^C^ overexpression was associated with increased susceptibility to neurotoxicity and cell death [15, 17–19]. This might mean that *Prnp* levels should be constrained to a certain level to develop their natural functions [11, 15]. This balance is altered in several injuries and neurodegenerative processes, presenting changes in mRNA and protein expression, for example, in Alzheimer’s disease, dementia with Lewy bodies, some tauopathies [20, 21], human prionopathies (e.g., sporadic Creutzfeldt Jakob Disease (sCJD) [22]), and multiple sclerosis [23].

As indicated, numerous studies have explored the physiological roles of PrP^C^ *in vivo* using *Prnp^0/0^* mice. However, it was later demonstrated that some physiological functions were unfortunately misattributed to PrP^C^ due to genetic artifacts generated during the production of the most commonly used knock-out model, still in use in some laboratories: the Zürich I (*Prnp^ZH1/ZH1^*) mouse [24]. This mouse displayed a mixed background (B6129: C57BL6/J + 129Sv) and was further backcrossed with C57BL/6J mice to generate the B6.129 mouse line [19, 25], and with FVB mice to generate the FVBN-*Prnp^ZH1/ZH1^* model [25] in order to reduce the 129/Sv-associated genes. However, the generated lines were systematically confounded by the *Prnp*-linked loci polymorphic region containing numerous 129/Sv-associated “*flanking genes*” (FG) close to *Prnp* deletion [26, 27]. In fact, after crossing *Prnp^ZH1/ZH1^* mice with C57BL/6J for more than ten generations to reduce FG, a remnant of ≈ 2-5 % of the 129/Sv genome markers still persisted in B6.129 mice [26, 28, 29]. Genome analysis of these models revealed that the number of FG in the chromosome 2 regions where *Prnp* is localized is 62, related to different functions (i.e., cancer, depression, anxiety, among others [26]). Thus, we consider that most of the physiological phenotypes attributed to the *Prnp* absence or overexpression, using these mouse models carrying FG, need to be revaluated and confirmed in other FG-free models. As an example, in previous studies, we and others demonstrated that these FGs masked the real neuroprotective function of PrP^C^ against kainate (KA) administration *in vivo* [19, 30–32]. Although a full description of the FG-associated effects in a null *Prnp* background is not available, one of these FGs is the signal regulatory protein alpha (Sirpα), an important regulator of several innate immune functions [33]. Although prion disease evolution is not modified in *Sirpα^0/0^* mice [34], it has been clearly demonstrated that Sirpα is mainly responsible for a phagocytic function previously attributed to PrP^C^ [26, 27]. The number of functions misattributed to PrP^C^ increased when a recent study described a substrain-related dependence of Cu(I)-ATPase activity among *Prnp^0/0^* mice related to the 129/Sv FGs and not PrP^C^ [35].

In neurons, PrP^C^ is transported along axons [36] and enriched at the synaptic terminal, where it has been described that it interacts with some subunits of the glutamate receptors (e.g., NMDA-R, GluK2/GluK3, GluN2D, or mGluR5), modulating their activity, and with anchoring proteins such as PSD-95 [37–39]. However, due to the different mouse strains used and some experimental differences, the involvement of PrP^C^ in neurotransmission is still elusive. For example, different studies reported reduced [40–43], normal [25], or enhanced [19, 44] long-term potentiation (LTP) in *Prnp^0/0^* mice compared to wild-type mice. Following these descriptions, the consequences of *Prnp* absence in memory, learning, and behavior lead to variable results in studies using mice carrying FGs [24, 45–48] or not [42, 49].

In the present study, we focused our attention on reexamining some PrP^C^ functions associated with neurotransmission, learning, and behavior, taking advantage of a recently generated *Prnp^0/0^* mouse model: the Zürich 3 (*Prnp^ZH3/ZH3^*) [50]. This co-isogenic mouse was generated in a pure C57BL/6J background using TALEN technology [50] and it is resistant to prion infection [51]. Here we performed a set of behavioral tests to analyze *Prnp^ZH3/ZH3^* mouse activity, learning, and memory capabilities. In addition, basic synaptic functions, KA-mediated excitability, and LTP induction were evaluated electrophysiologically in alert behaving mice. Finally, PrP^C^ roles during neuronal differentiation and activity were also evaluated in primary cortical cultures. Results indicate that adult *Prnp^ZH3/ZH3^* mice display reduced activity and anxiety-like behavior. They also fail to acquire different instrumental learning tasks. In addition, our experiments show that hippocampal CA3-CA1 *Prnp^ZH3/ZH3^* synapse fails to induce LTP, most likely due to an exacerbated endogenous excitability, further corroborated *in vivo* after KA administration. Lastly, our results are sustained by the observed alteration in the expression patterns of several genes associated with neuronal system function and synaptic protein-protein interactions in the *Prnp^ZH3/ZH3^* hippocampus by an RNAseq analysis and its RT-qPCR validation.

## Results

The absence of PrP^C^ has been related to deficiencies in behavior, learning, and memory in several mouse models with different results [24, 42, 52, 53]. In order to evaluate the implication of PrP^C^ in systemic behavioral tasks, we took advantage of the new knock-out model, the *Prnp^ZH3/ZH3^* mouse. First, we analyzed the nest building capacity between *Prnp^ZH3/ZH3^ vs. Prnp^+/+^* mice as an indicator of mouse welfare. In contrast to Schmitz *et al.,* where they used *Prnp^ZH1/ZH1^* mice (reviewed in [54]), results showed a slightly increased but not significantly nest-building capacity in *Prnp^ZH3/ZH3^* compared to controls (*Prnp^ZH3/ZH3^* = 3.81 ± 0.64, n = 7 *vs. Prnp^+/+^* = 3.00 ± 0.41, n = 7; mean ± S.E.M., *p* = 0.15; Mann-Whitney *U* non-parametric test) (Additional file 1: Fig. S1), suggesting that the two genotypes had similar welfare conditions [55].

### Reduced activity, increased thigmotaxis, and anxiety-related behavior in Prnp^ZH3/ZH3^ mice

First, we performed the open field test to measure the general locomotor activity levels, anxiety, and willingness in knock-out mice (Fig. 1). *Prnp^+/+^* and *Prnp^ZH3/ZH3^* mice (n = 49 for each genotype) were individually placed (in rounds of two animals in parallel) in the open field arena for 15 min, and their activity was evaluated on the X-Y-Z axes (Fig. 1b). *Prnp^ZH3/ZH3^* mice showed significantly reduced displacement in the field (*Prnp^+/+^* = 3725 ± 93 a.u. *vs. Prnp^ZH3/ZH3^* = 3370 ± 95 a.u.; \*\**p* = 0.009; Mann-Whitney *U* non-parametric test) (Fig 1c). Anxiety and stress increased the thigmotaxis behavior and the natural aversion to exploring the inner square of the field during the test [56]. Thus, to evaluate anxiety-like behavior in the *Prnp^ZH3/ZH3^* mice, we measured this thigmotaxis performance as the time spent in the center (inner region of the field) *vs.* the periphery (outer region of the field) for each mouse (Fig. 1b). *Prnp^+/+^* mice spent the same amount of time in the two regions, while *Prnp^ZH3/ZH3^* animals remained significantly more time in the periphery close to the walls, suggesting an apprehension of the center of the field that reflects an anxiety-like behavior (*Prnp^+/+^*: Center = 303.3 ± 14.5 *vs.* Periphery = 345.7 ± 18,0; and *Prnp^ZH3/ZH3^*: Center = 288.2 ± 15.1 *vs.* Periphery = 394.3 ± 20.3; mean ± S.E.M., *p* = 0.071 and \*\*\**p* < 0.001 respectively; Mann-Whitney *U* non-parametric test) (Fig. 1d). Stressed behavior was also assessed by counting the number of rearing and immobility episodes during the test. *Prnp^ZH3/ZH3^* mice displayed significantly fewer rearing episodes (*Prnp^+/+^* = 52 ± 2 *vs. Prnp^ZH3/ZH3^* = 26.6 ± 1.7; mean ± S.E.M., \*\*\**p* < 0,001; Mann-Whitney *U* non-parametric test) and more immobility episodes (*Prnp^+/+^* = 6.6 ± 1.2 *vs. Prnp^ZH3/ZH3^* = 12.4 ± 0.9; mean ± S.E.M., \*\*\**p* < 0.001; Mann-Whitney *U* non-parametric test) confirming an anxiety-like behavior (Fig. 1e).

**Fig. 1.**
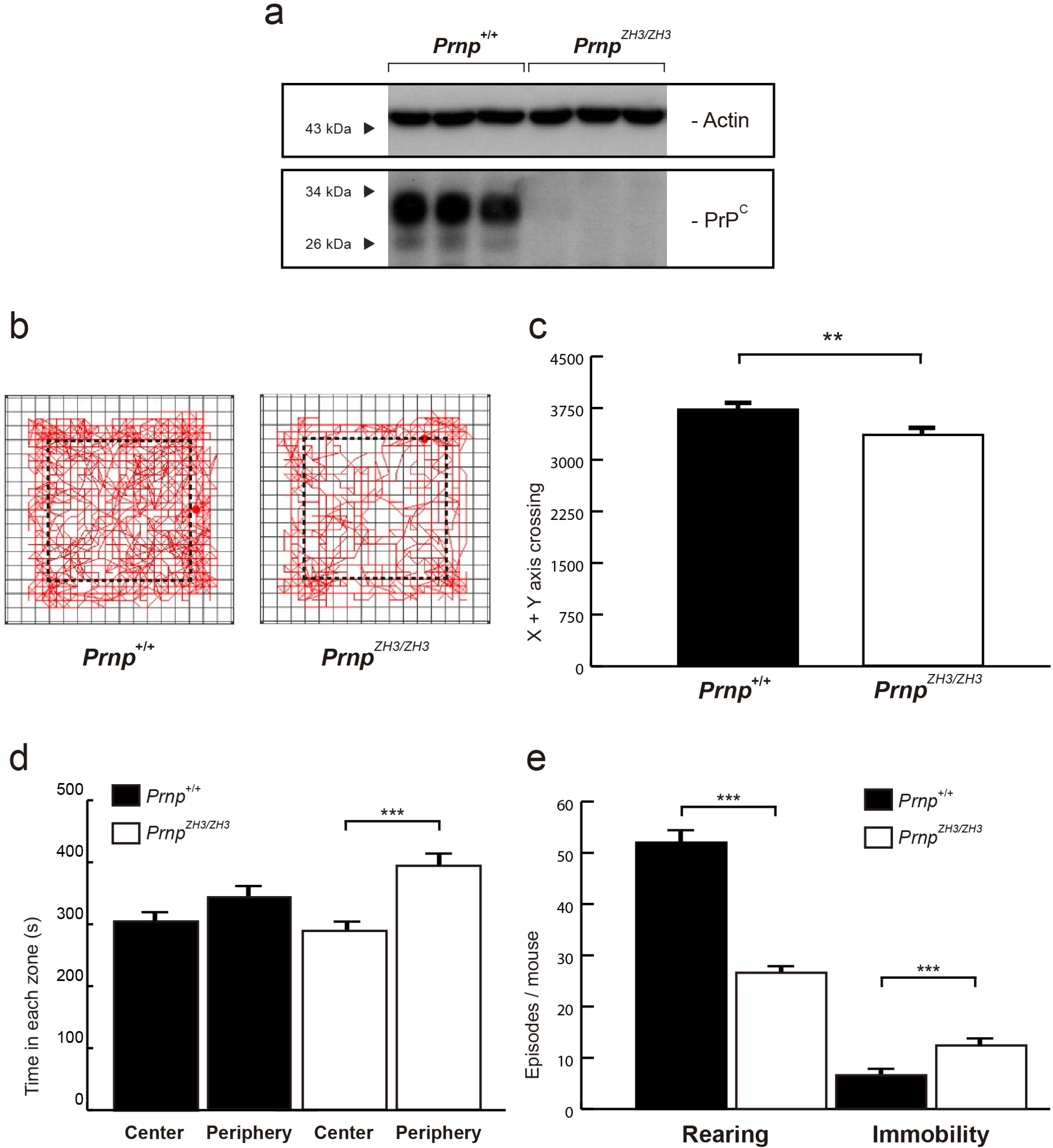
*Prnp^ZH3/ZH3^* mice show reduced activity and anxiety-related behavior. ***a***, Immunoblot analysis of PrP^C^ expression in *Prnp^+/+^* and *Prnp^ZH3/ZH3^* mice forebrain. Actin is used as a loading control. ***b***, Representative images of *Prnp^+/+^* and *Prnp^ZH3/ZH3^* mouse exploratory behavior in the open field test. The dotted line delineates the center and the periphery of the arena. ***c***, Mouse activity in the open field test represented as the number of lines crossed in the X+Y axis. ***d***, Time spent (s) by the mice in the center and periphery of the open field arena. ***e***, Number of rearing and immobility episodes displayed by *Prnp^+/+^* and *Prnp^ZH3/ZH3^* mice during the open field test. 98 animals (n = 49 for each genotype) were tested individually, in rounds of two animals in parallel. Data are presented as mean ± S.E.M. \*\**p* < 0.01 and \*\*\**p* < 0.001, Mann-Whitney *U* non-parametric test.

### Prnp^ZH3/ZH3^ mice and Prnp^ZH1/ZH1^ failed to acquire instrumental learning tasks

Our next goal was to examine the capabilities of *Prnp^ZH3/ZH3^* mice in performing highly demanding learning tasks. Instrumental learning capabilities were tested with operant conditioning in the Skinner box (n = 49 for each genotype) (Fig. 2). Collected data were compared to those obtained using *Prnp^ZH1/ZH1^* mice (Additional file 2: Fig. S2). Thirty-one percent of *Prnp^ZH3/ZH3^* mice did not reach the learning criterion (to obtain ≥ 20 pellets for two consecutive sessions) at the end of the training session. In contrast, all *Prnp^+/+^* mice (100 %) meet the selected criterion from the 6th session (Fig. 2a). Similarly, in a second set of experiments using *Prnp^ZH1/ZH1^* mice (*Prnp^+/+^* = 10 and *Prnp^ZH1/ZH1^* = 10), 50 % of *Prnp^ZH1/ZH1^* mice failed to reach the criterion, but 80 % of wild-type mice reached it at the end of the sessions (Additional file 2: Fig. S2a). These results strongly suggest that mice lacking *Prnp* (both ZH1 and ZH3 backgrounds) present evident instrumental learning deficiencies. Also, *Prnp^+/+^* mice pressed the lever significantly more times from session 3 onwards than *Prnp^ZH3/ZH3^* animals (Session 1: *Prnp^+/+^* = 7.8 ± 1.3 *vs. Prnp^ZH3/ZH3^* = 11.12 ± 1.9, *p > 0.99*; Session 2: *Prnp^+/+^* = 21.6 ± 2.51 *vs. Prnp^ZH3/ZH3^* = 15.8 ± 2.2, *p > 0.99;* Session 3: *Prnp^+/+^* = 40.2 ± 5.1 *vs. Prnp^ZH3/ZH3^* = 23.3 ± 3.2, \*\**p* = 0.0023; Session 4: *Prnp^+/+^* = 53.0 ± 3.9 *vs. Prnp^ZH3/ZH3^* = 29.4 ± 3.1, \*\*\**p* < 0.0001; Session 5: *Prnp^+/+^* = 55.1 ± 3.3 *vs. Prnp^ZH3/ZH3^* = 36.1 ± 3.9, \*\*\**p* = 0.0003; Session 6: *Prnp^+/+^* = 70.5 ± 4.1 *vs*. *Prnp^ZH3/ZH3^* = 38.3 ± 3.9 \*\*\**p* < 0.0001; Session 7: *Prnp^+/+^* = 59.9 ± 2.3 *vs Prnp^ZH3/ZH3^* = 39.6 ± 3.5, \*\*\**p* < 0.0001; two-way ANOVA + Bonferroni’s multiple comparisons test) (Fig. 2b). However, as observed in the open field test, *Prnp^ZH3/ZH3^* mice presented considerable inactive behaviors (Fig. 1). To distinguish the reduction of activity from real learning deficits, 44 mice (*Prnp^+/+^* = 24 and *Prnp^ZH3/ZH3^* = 20) were subjected to a more complex operant conditioning paradigm, the light ON/light OFF task (see Material and methods). As expected, the total number of lever pulses during the OFF period was higher in the *Prnp^+/+^* mice and drastically reduced along with sessions (Session 1 = 134.8 ± 13.3 *vs.* Session 10 = 39.7 ± 5.2) (Fig. 2c). In parallel, *Prnp^+/+^* mice increased the number of pulses in the ON period (Session 1 = 26.9 ± 1.8 *vs.* Session 10 = 45.7 ± 2.8) (Fig. 2c). In contrast, *Prnp^ZH3/ZH3^* mice showed a reduced decrease of pulses in the OFF period (Session 1 = 73.2 ± 11.7 *vs.* Session 10 = 26.5 ± 3.9) and an incipient increase in the ON period (Session 1 = 23.4 ± 1.6 *vs.* Session 10 = 31.1 ± 2.9) (Fig. 2d). Learning capacity, measured as the difference in the curve slope during ON or OFF periods, was drastically reduced in *Prnp^ZH3/ZH3^* mice (OFF: *Prnp^+/+^* = -12.5, R^2^ = 0.90; *Prnp^ZH3/ZH3^* = -5.9, R^2^ = 0.92; ON: *Prnp^+/+^* = 2.3, R^2^ = 0.96; *Prnp^ZH3/ZH3^* = 0.8, R^2^ = 0.48) (Figs. 2c-d). These differences show that *Prnp^ZH3/ZH3^* mice failed to learn to avoid OFF periods and push the lever during the ON periods, indicating that PrP^C^ seems to be necessary to properly acquire instrumental learning tasks. A similar study was developed using *Prnp^ZH1/ZH1^* mice, and the task accuracy ratio ((lever presses during light ON – lever presses during light OFF)/(total number of lever presses)) was evaluated. At the end of the experiment (sessions 7 and 8), the *Prnp^ZH1/ZH1^* mice showed lower values than wild-type mice (session 7: *Prnp^+/+^* = 0,31 *vs. Prnp^ZH1/ZH1^* = -0.02; session 8: *Prnp^+/+^* = 0,54 *vs. Prnp^ZH1/ZH1^* = 0.1), reinforcing the notion that the absence of PrP^C^ decreases the instrumental learning goals in mutant mice (Additional file 2: Fig. S2b).

**Fig. 2.**
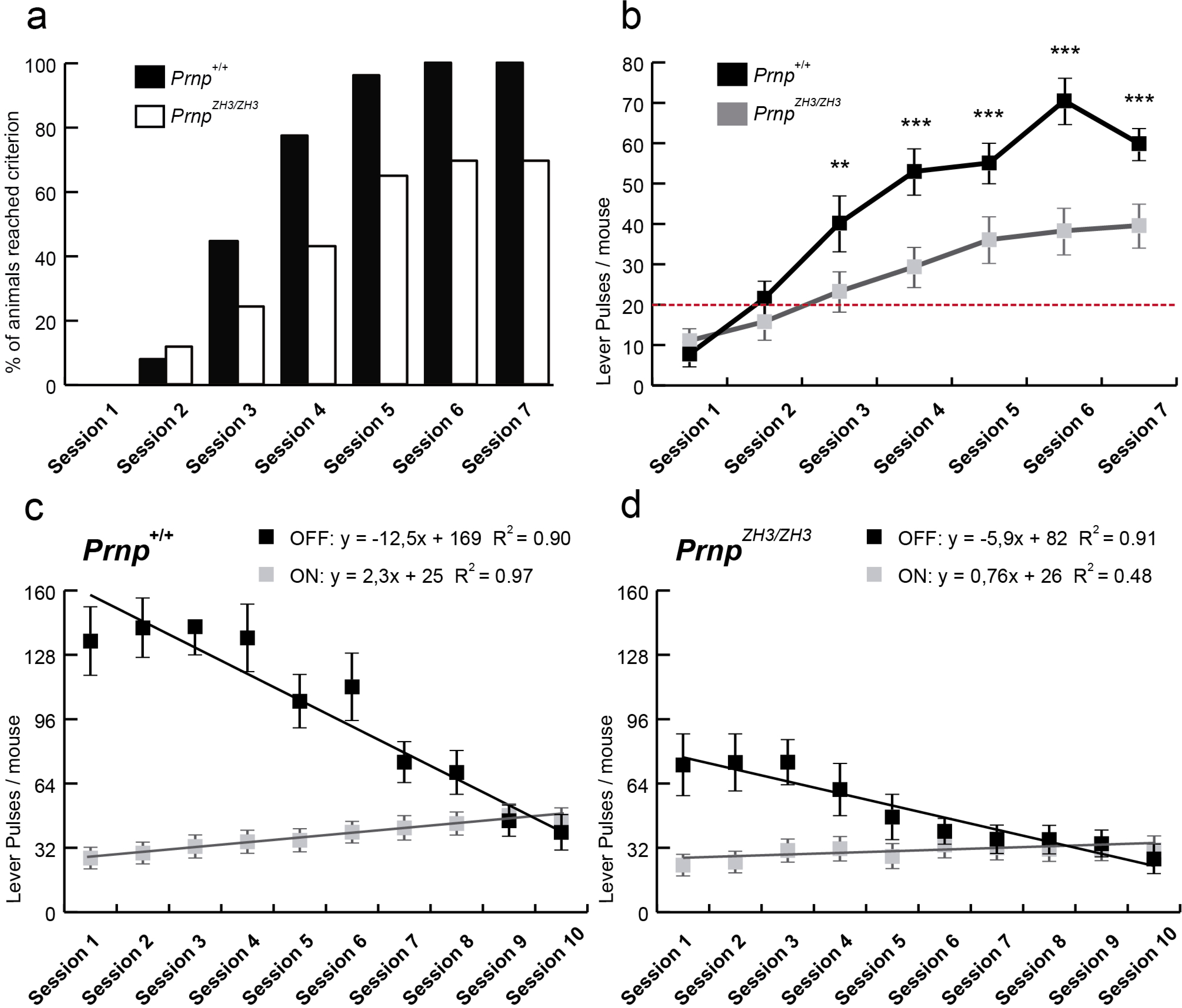
Impairments in the acquisition of an instrumental learning task in *Prnp^ZH3/ZH3^* mice. ***a***, Percentage of mice reaching the selected criterion (to collect ≥ 20 food pellets for two consecutive days) in the successive training sessions. ***b***, Lever presses of *Prnp^+/+^* and *Prnp^ZH3/ZH3^* mice during the fixed ratio (1:1) conditioning paradigm. The test was performed daily for seven consecutive days. ***c-d***, Lever presses of *Prnp^+/+^* (***c***) and *Prnp^ZH3/ZH3^* (***d***) mice during the ON/OFF training paradigm. Lines represent best linear fits for lever presses evoked during light ON (grey) and light OFF (black) periods. Equations corresponding to regression lines are illustrated in (***c***) and **(*d*)**, including R2 coefficients. Data are presented as a percentage in (***a***), and as mean ± S.E.M. in (***b-d***). \*\**p* < 0.01 and \*\*\**p* < 0.001, two-way ANOVA + Bonferroni’s multiple comparisons test.

To test for possible deficiencies in motor coordination and balance that could also affect in the results from the Skinner box and the open field tests, we compared the performances of both *Prnp^+/+^* and *Prnp^ZH3/ZH3^* mice in the accelerating rotarod test. After a training session, the mice’ latency to fall from the rods was tested for five sessions during two consecutive days. In the first day, the *Prnp^ZH3/ZH3^* mice displayed a significantly lower latency just in the first two session compared to the control mice (Additional file 3: Fig. S3a). However, from the third run, their performance was not significantly different (Session 1: *Prnp^+/+^* = 48.51 ± 3.34 *vs*. *Prnp^ZH3/ZH3^* = 24.48 ± 7.56, \**p* = 0.014; Session 2: *Prnp^+/+^* = 56.65 ± 6.56 *vs*. *Prnp^ZH3/ZH3^* = 27.30 ± 8.03, \*\**p* = 0.007; Session 3: *Prnp^+/+^* = 61.49 ± 4.69 *vs. Prnp^ZH3/ZH3^* = 41.11 ± 8.20, *p* = 0.121; Session 4: *Prnp^+/+^* = 55.71 ± 2.82 *vs. Prnp^ZH3/ZH3^* = 37.00 ± 6.45, *p* = 0.189; Session 5: *Prnp^+/+^* = 54.30 ± 1.73 *vs*. *Prnp^ZH3/ZH3^* = 38.39 ± 6.44, *p* = 0.380; mean ± S.E.M., two-way ANOVA + Bonferroni’s multiple comparisons test). In the second day, no significant difference was observed (Session 1: *Prnp^+/+^* = 45.61 ± 5.59 *vs*. *Prnp^ZH3/ZH3^* = 31.95 ± 8.16; Session 2: *Prnp^+/+^* = 53.54 ± 3.25 *vs*. *Prnp^ZH3/ZH3^* = 44.99 ± 7.00; Session 3: *Prnp^+/+^* = 53.67 ± 4.68 *vs*. *Prnp^ZH3/ZH3^* = 47.23 ± 8.03; Session 4: *Prnp^+/+^* = 59.91 ± 5.48 *vs. Prnp^ZH3/ZH3^* = 48.16 ± 9.06; Session 5: *Prnp^+/+^* = 52.64 ± 3.35 *vs*. *Prnp^ZH3/ZH3^* = 59.01 ± 9.76; in all the sessions *p* > 0.89, mean ± S.E.M.; two-way ANOVA + Bonferroni’s multiple comparisons test) (Additional file 3: Fig. S3b). These results indicate similar motor capacities in both groups, but *Prnp^ZH3/ZH3^* mice needed more trials to reach steady measurements for the same task compared to *Prnp^+/+^* mice, suggesting that the knock-out mice have deficits in acquiring instrumental learning as we also observe with the Skinner test (Fig. 2), but not any motor impairment.

Finally, episodic memory was evaluated with the object recognition test (Additional file 4: Fig. S4). In our experiments, most of the *Prnp^ZH3/ZH3^* mice interacted with the objects for just a few seconds (Additional file 4: Fig. S4c-d). Therefore, we ruled out this approach due to *Prnp^ZH3/ZH3^* mouse inactivity, related to the anxiety-like behavior we observed in the open field test. This inactivity led to reduced interactions with the objects that rendered the learning results unreliable. To further support increased anxiety levels in the object recognition test, the fecal bodies left in the arena after the habituation session were counted by the observer once the test subject was removed. *Prnp^ZH3/ZH3^* mice exhibited significant increase in fecal bodies present when compared to wild-type mice (*Prnp^ZH3/ZH3^* = 4.76 ± 0.56 *vs. Prnp^+/+^* = 1.29 ± 0.52. mean ± S.E.M., *** *p* < 0.001, Mann-Whitney *U* non-parametric test) (Additional file 4: Fig. S4b). These results also correlate with the thigmotaxis levels measured in the *Prnp^ZH3/ZH3^* mice and indicate that the knock-out mice showed increased emotionality and anxiety compared to their wild-type counterparts. This was in contrast to what was observed using *Prnp^ZH1/ZH1^*mice, where the test could be performed. In the habituation session, knock-out mice showed a significant decrease in rearing episodes (*Prnp^+/+^* = 43.21 ± 10.01 *vs*. *Prnp^ZH1/ZH1^* = 15.7 ± 2.8, mean ± S.E.M., \*\*\**p* < 0.001, Mann-Whitney *U* non-parametric test) (Additional file 4: Fig. S4e). No changes were observed in the training phase (Additional file 4: Fig. S4f), although the *Prnp^ZH1/ZH1^* showed a tendency (*p* = 0.053) to explore the second object less time ((second - first) / total) compared to the wild-type mice (*Prnp^+/+^* = 0.3 ± 0.15 *vs. Prnp^ZH1/ZH1^* = - 0.17 ± 0.15, mean ± S.E.M.) (Additional file 4: Fig. S4g).

### Increased paired-pulse facilitation at high intensities in Prnp^ZH3/ZH3^ Schaffer collateral pathway

PrP^C^ has been described as a regulator of glutamatergic neurotransmission in the hippocampus [19, 37]. As an example, PrP^C^ has been shown to inhibit NMDAr containing the NR2D subunit [37], or the GluR6/7 receptor [38] see also [57] for review. Therefore, we analyzed the activation of the well-characterized hippocampal Schaffer collateral pathway (CA3-CA1 synapses). Stimulating and recording electrodes were permanently implanted in the CA3 and CA1 regions, allowing us to record and quantify the evoked fEPSPs in living behaving mice (Fig. 3a). First, we evaluated the putative synaptic facilitation or depression evoked at CA3-CA1 synapses by paired-pulse stimulation of the ipsilateral Schaffer collaterals (Fig. 3b). Paired-pulse stimuli were presented to *Prnp^+/+^* (n = 27) and *Prnp^ZH3/ZH3^* (n = 24) mice at different inter-stimulus intervals (from 10 to 500 ms). As already reported for CA3-CA1 synapses [58], this approach generates a higher fEPSP from the second stimulus (fEPSP2) than from the first (fEPSP1) at short intervals due to presynaptic facilitation. In our experiments, as can be observed in the representative examples of fEPSP1 and fEPSP2 (Fig. 3c), no differences were observed between *Prnp^+/+^* and *Prnp^ZH3/ZH3^* facilitation (Fig. 3b), suggesting that PrP^C^ does not participate in presynaptic mechanisms related to synaptic facilitation at the least at the selected intensities (2 × Threshold; ≈ 0.2 mA).

**Fig. 3.**
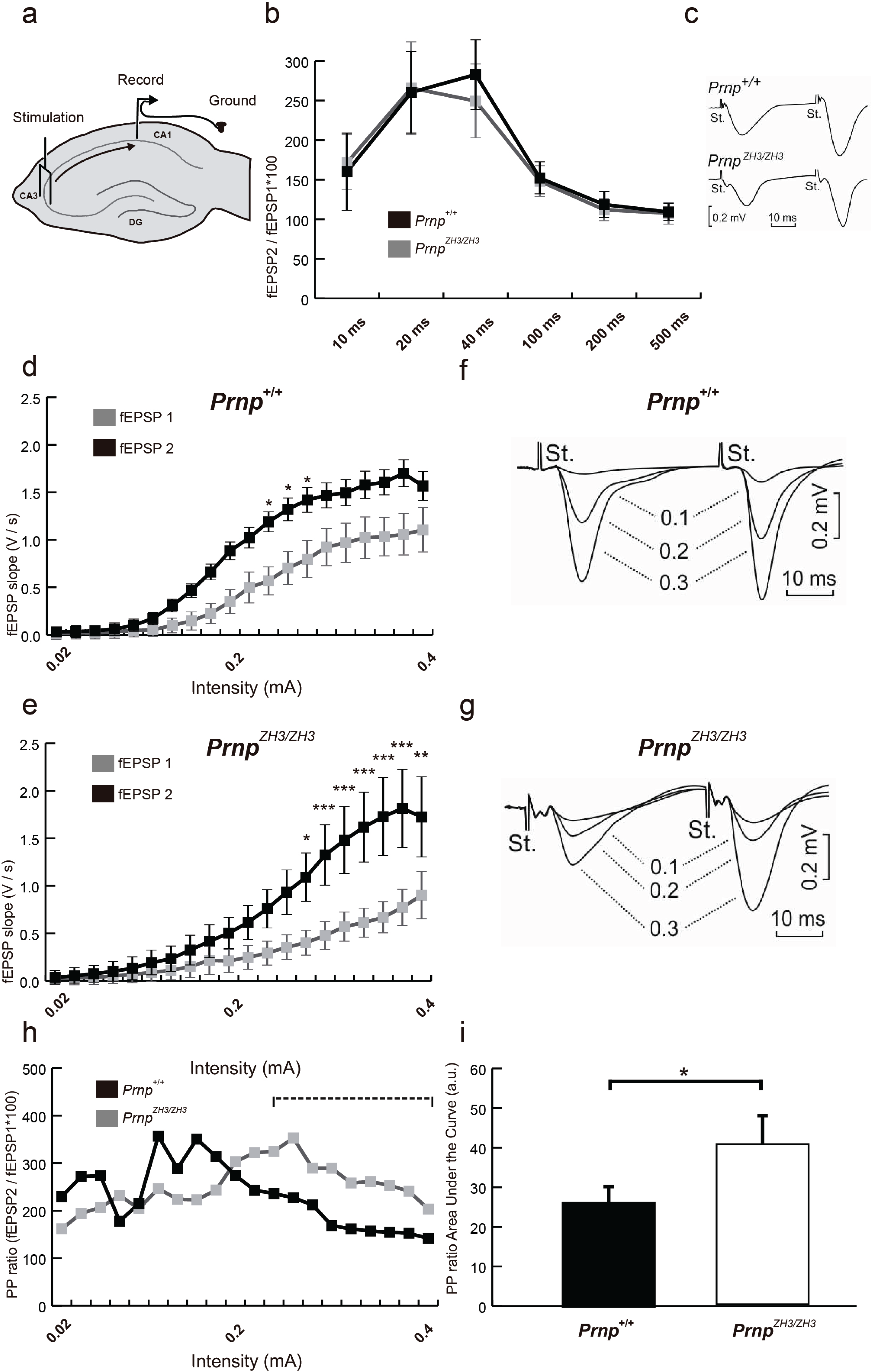
CA3-CA1 synapses in *Prnp^ZH3/ZH3^* mice show enhanced excitability. ***a***, Schematic representation of electrodes implanted in mouse dorsal hippocampus. Two stimulation electrodes are implanted in the Schaffer collateral pathway in the CA3 region and two recording electrodes in the CA1 *stratum radiatum*. ***b***, Effects of the paired-pulse stimulation of the Schaffer collateral pathway at increasing inter-stimulus intervals (10, 20, 40, 100, 200, 500 ms). Data are presented as the percentage of increase of fEPSP2 in relation to fEPSP1 (fEPSP2/fEPSP1 * 100). ***c***, The inset illustrates representative examples of fEPSPs (averaged 5 times) evoked by paired pulses (40 ms of inter-pulse interval) of similar intensities (2 × Threshold; ≈ 0.2 mA) in *Prnp^+/+^* and *Prnp^ZH3/ZH3^* mice. ***d****-**e**,* Input/output curves of fEPSPs (V/s) in CA1 after the presentation of paired-pulses of increasing intensities in the CA3 area (0.02 mA to 0.4 mA) of *Prnp^+/+^* (***d***) and *Prnp^ZH3/ZH3^* **(*e*)** mice. ***f****-**g**,* The insets show representative recordings of fEPSPs evoked in *Prnp^+/+^* (***f***) and *Prnp^ZH3/ZH3^* (***g***) mice by paired pulses (40 ms of inter-pulse interval) of similar intensities (0.1 mA, 0.2 mA, and 0.3 mA). ***h***, Paired-pulse ratio (fEPSP2 / fEPSP1 * 100) of data illustrated in *(**d**)*, (***e***) and *(**i**)*, Area under the curve (a.u.) of PP ratio from 0.24 mA to 0.4 intensities. Data are presented as mean ± S.E.M; *p* < 0.05, \*\**p* < 0.01, and \*\*\**p* < 0.001, two-way ANOVA + Bonferroni’s multiple comparisons test.

Next, we analyzed the consequences of PrP^C^ deficiency in hippocampal synaptic excitability at a large range of stimulus intensities (Fig. 3d-g). The slope of fEPSP facilitation evoked by paired-pulse (40 ms inter-stimulus interval) stimulation was measured at increasing intensities (from 0.02 to 0.4 mA). In *Prnp^+/+^* mice (n = 14), fEPSP1 and fEPSP2 increased steadily more or less in parallel after 0.18 mA stimulation, reaching asymptotic values at 0.32 mA (Fig. 3d-e). fESPS2 was significantly greater in three stimulation intensities before arriving at the asymptotic values showing synaptic facilitation (0.20 mA: fESPS1 = 0.35 ± 0.1 V/s and fEPSP2 = 0.88 ± 0.2 V/s, *p* = 0.124; 0.22 mA: fESPS1 = 0.50 ± 0.1 V/s and fEPSP2 = 1.02 ± 0.2 V/s, *p* = 0.144; 0.24 mA: fESPS1 = 0.57 ± 0.1 V/s and fEPSP2 = 1.19 ± 0.2 V/s, \**p* = 0.030; 0.26 mA: fESPS1 = 0.70 ± 0.1 V/s and fEPSP2 = 1.32 ± 0.2 V/s, \**p* = 0.031; 0.28 mA: fESPS1 = 0.80 ± 0.1 V/s and fEPSP2 = 1.42, ± 0.2 V/s, *\*p* = 0.028, 0.30 mA: fESPS1 = 0.92 ± 0.1 V/s and fEPSP2 = 1.47, ± 0.2 V/s, *p* = 0.111; 0.32 mA: fESPS1 = 0.97 ± 0.2 V/s and fEPSP2 = 1.50, ± 0.3 V/s, *p* = 0.192; 0.34 mA: fESPS1 = 1.02 ± 0.2 V/s and fEPSP2 = 1.58, ± 0.3 V/s, *p* = 0.172; mean ± S.E.M.; two-way ANOVA + Bonferroni’s multiple comparisons test). From 0.26 mA stimulation, fEPSP1 and fEPSP2 were statistically equal in *Prnp^+/+^* mice; thus, there was no synaptic facilitation at high intensities (Fig. 3d,f). This phenomenon has been described as a putative protective mechanism in high intensity insults to maintain hippocampal homeostasis [58]. In contrast, in *Prnp^ZH3/ZH3^* mice (n = 15), fEPSP1 and fEPSP2 did not increase in parallel, showing an increased facilitation to paired-pulse presentations, and therefore suggesting the absence of this protective mechanism (Fig. 3e and g). fEPSP2 was significantly greater than fEPSP1 at higher intensities (0.24 mA: fESPS1 = 0.293 ± 0.1 V/s and fEPSP2 = 0.76 ± 0.1 V/s, *p* = 0.36; 0.26 mA: fESPS1 = 0.35 ± 0.1 V/s and fEPSP2 = 0.93 ± 0.2 V/s, *p* = 0.069; 0.28 mA: fESPS1 = 0.4 ± 0.1 V/s and fEPSP2 = 1.09 ± 0.2 V/s, \**p* = 0.010; 0.30 mA: fESPS1 = 0.48 ± 0.1 V/s and fEPSP2 = 1.33 ± 0.3 V/s, *\*\*\*p* = 0.0008; 0.32 mA: fESPS1 = 0.57 ± 0.1 V/s and fEPSP2 = 1.5 ± 0.3 V/s, *\*\*\*p* = 0.0002; 0.34 mA: fESPS1 = 0.61 ± 0.1 V/s and fEPSP2 = 1.62 ± 0.3 V/s, *\*\*\*p* < 0.0001; 0.36 mA: fESPS1 = 0.67 ± 0.1 V/s and fEPSP2 = 1.73 ± 0.3 V/s, \*\*\**p* < 0.0001; 0.38 mA: fESPS1 = 0.77 ± 0.1 V/s and fEPSP2 = 1.81 ± 0.4 V/s, \*\*\**p* < 0.0001 0.40 mA: fESPS1 = 0.9 ± 0.2 V/s and fEPSP2 = 1.73 ± 0.4 V/s, *\*\*p* = 0.004; mean ± S.E.M.; two-way ANOVA + Bonferroni’s multiple comparisons test). *Prnp^+/+^* fEPSP1 increased steadily to greater asymptotic values than *Prnp^ZH3/ZH3^* fEPSP1 (from 0.3 mA stimulation ≈ 50 % increased), but fEPSP2 were almost equal (fEPSP1: *Prnp^+/+^* ≈ 1.0 V/s; *Prnp^ZH3/ZH3^* ≈ 0.7 V/s and fEPSP2: *Prnp^+/+^* ≈ 1.5 V/s; *Prnp^ZH3/ZH3^* ≈ 1.6 V/s). Consequently, the increase in fEPSP1 related to fEPSP2 was ≈ 50 % in *Prnp^+/+^* individuals, but > 140 % in their *Prnp^ZH3/ZH3^* counterparts.

Exacerbation of synaptic facilitation was clearly observed with the paired-pulse (PP) ratio (fEPSP2/fEPSP1 x 100). At high intensities, the PP ratio was larger in *Prnp^ZH3/ZH3^* connections (Fig. 3h). The area under the curve (AUC) from 0.24 mA intensity was significantly lower in the *Prnp^+/+^* (*Prnp^+/+^* = 26.07 ± 4.0 a.u *vs. Prnp^ZH3/ZH3^* = 40.75 ± 7.1 a.u., mean ± S.E.M., \**p* = 0.04; Mann-Whitney *U* non-parametric test) (Fig. 3i). These results suggest that PrP^C^ regulates neuronal excitability or, perhaps, synaptic homeostasis at high-intensity stimulations, hinting at a neuroprotective role.

### High-frequency stimulation evoked epileptic seizures in Prnp^ZH3/ZH3^ Schaffer collaterals but failed to increase LTP

As indicated, several studies reported differing data on LTP in *Prnp^0/0^* mice (see Background). So, to analyze LTP in *Prnp^ZH3/ZH3^* mice, we performed an LTP induction protocol based on high-frequency stimulation (HFS) in 40 mice (*Prnp^+/+^* = 20 and *Prnp^ZH3/ZH3^* = 20) (Fig. 4). First, the baseline fEPSPs were recorded for 15 min, evoked by double pulses at an inter-stimulus interval of 40 ms. Afterward, the HFS protocol was presented. This consisted of five trains (200 Hz, 100 ms) of pulses (1/s) presented six times (1/min). Recordings were maintained for 60 min immediately after the HFS and repeated 30 min daily for four days from the HFS presentation session. *Prnp^+/+^* displayed significant LTP for both pulses (Fig. 4a). fESPS1 and fESPS2 recordings were significantly larger than the baseline after the HFS, and this potentiation lasted for the five days of recording sessions (Fig. 4e). As expected, HFS reduced paired-pulsed facilitation on the first day [58]. However, from the second day, facilitation recovered steadily but with a range of increase from 350 % to 150 % with respect to fEPSP1 baseline (Baseline: fEPSP1 = 100 %; fEPSP2 = 268.2 ± 43.1 %; Day 1: fEPSP1 = 478.3 ± 78.4 %, \*\*\**p* < 0,001; fEPSP2 = 441.1 ± 140.2 %, \**p* = 0,040; Day 2: fEPSP1 = 399.9 ± 65.0 %, \*\*\**p* < 0,001; fEPSP2 = 554.8 ± 164.2 %, \*\*\**p* < 0,001; Day 3: fEPSP1 = 310.4 ± 47.0 %, \*\*\**p* < 0,001; fEPSP2 = 466.3 ± 106.7 %, \**p* = 0,012; Day 4: fEPSP1 = 279.2 ± 36.0 %, \*\*\**p* < 0,001; fEPSP2 = 438.1 ± 90.7 %, \**p* = 0,046; Day 5: fEPSP1 = 226.25 ± 29.4 %, \*\**p* = 0,0014; fEPSP2 = 433.14 ± 87.7 %, *p* = 0,057; mean ± S.E.M.; two-way ANOVA + Bonferroni’s multiple comparisons test) (Fig. 4a). In contrast, in *Prnp^ZH3/ZH3^* mice, LTP induction was virtually absent and paired-pulsed facilitation was maintained (≈ 60 %) from the first day (e.g.,: Baseline: EPSP1 = 100 %; fEPSP2 = 158.4 ± 23.0 %; Day 1: fEPSP1 = 153.43 ± 15.7 %; fEPSP2 = 224.3 ± 39.8 %; Day 3: fEPSP1 = 114.2 ± 11.9 %; fEPSP2 = 160.4 ± 23.8 %; in all the sessions *p* > 0.6; mean ± S.E.M.; two-way ANOVA + Bonferroni’s multiple comparisons test) (Fig. 4b,f).

**Fig. 4.**
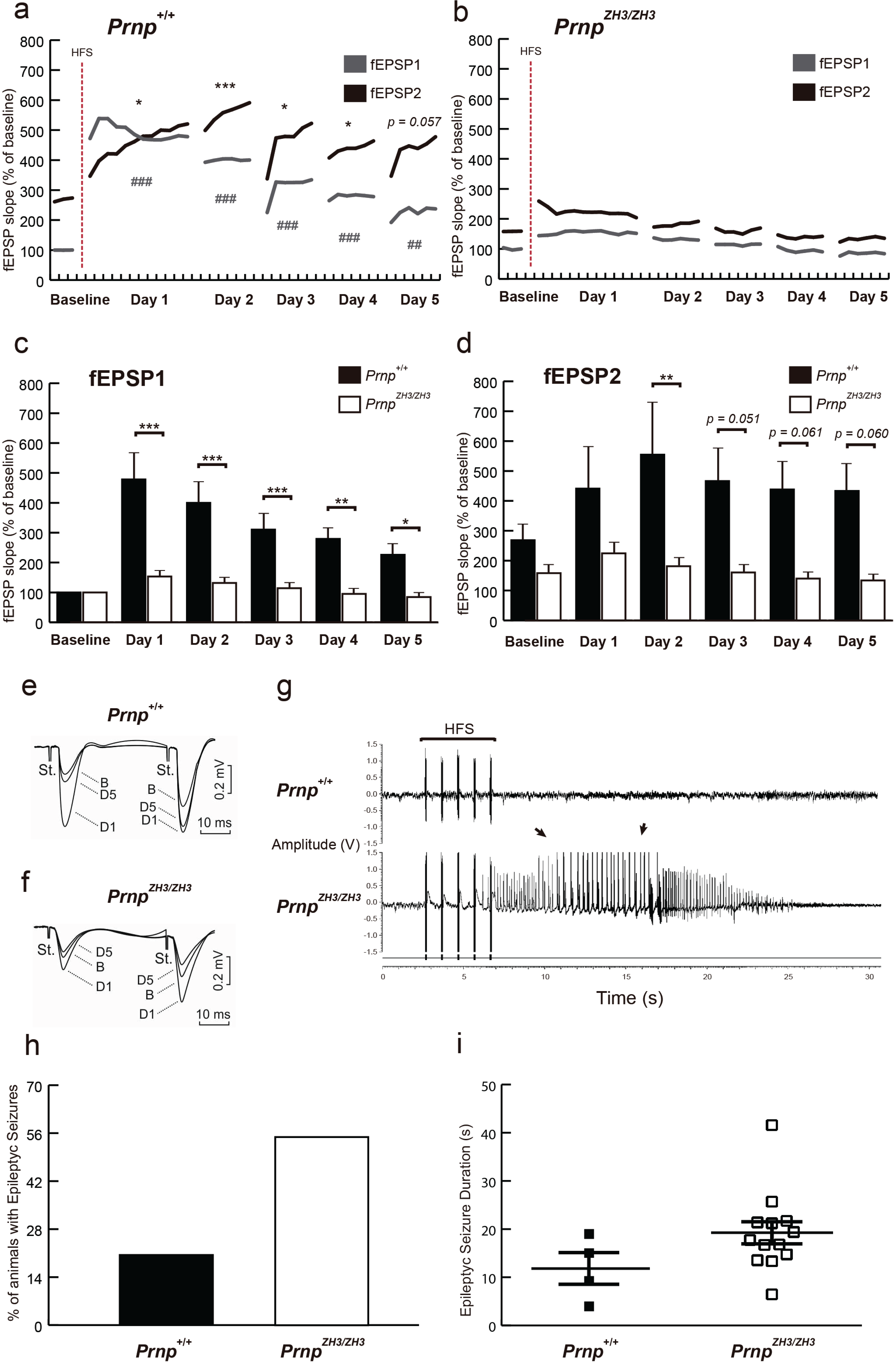
LTP is not induced at CA3-CA1 synapses of *Prnp^ZH3/ZH3^* mice, and the HFS presentation generates epileptic seizures. ***a***-***b***, Evolution of fEPSP1 evoked in the CA1 region by paired-pulsed stimulation of Schaffer collaterals for the *Prnp^+/+^* (***a***) and *Prnp^ZH3/ZH3^* (***b***) mice after the HFS session. Data are presented as the percentage of increase from baseline. Significant differences with baseline values are presented for fEPSP1 (#) and fEPSP2 (*) recordings in *Prnp^+/+^* mice. ***c***-***d***, fEPSP mean slopes from *Prnp^+/+^* (***c***), and *Prnp^ZH3/ZH3^* (***d***) mice before and after the HFS session. Data are presented as the percentage of increase from baseline values. ***e***-***f***, The insets show representative recordings (averaged 5 times) of fEPSPs evoked in *Prnp^+/+^* (***e***) and *Prnp^ZH3/ZH3^* (***f***) mice by paired pulses (40 ms of inter-pulse interval) of similar intensities (2 × Threshold; ≈ 0.2 mA). ***g***, Representative examples of long (30 s) recordings carried out after an HFS stimulation protocol in *Prnp^+/+^* and *Prnp^ZH3/ZH3^* Schaffer collaterals. Note the presence of a hippocampal seizure in the *Prnp^ZH3/ZH3^* mouse (arrows). ***h***, Percentage of mice that presented epileptic seizures following HFS presentations. ***i***, Seizure duration (s) following HFS. Data are presented as Mean ± S.E.M. \**p* < 0.05, \*\**p* < 0.01, \*\*\**p* < 0.001, ^##^*p* < 0.01 and ^###^*p* < 0.001, two-way ANOVA + Bonferroni’s multiple comparisons test and Mann-Whitney *U* non-parametric test.

In addition, *Prnp^+/+^* presented significantly larger fEPSP1 than *Prnp^ZH3/ZH3^* mice (Day 1: *Prnp^+/+^* = 478.3 ± 78.3 % *vs. Prnp^ZH3/ZH3^* = 153.4 ± 15.7 %, \*\*\**p* < 0,001; Day 2: *Prnp^+/+^* = 399.90 ± 65.0 % *vs. Prnp^ZH3/ZH3^* = 131.77 ± 12.3 %, \*\*\**p* < 0,001; Day 3: *Prnp^+/+^* = 310.42 ± 47.0 % *vs. Prnp^ZH3/ZH3^* = 114.22 ± 11.9 %, \*\*\**p* = 0.0009; Day 4: *Prnp^+/+^* = 279.20 ± 36.05 % *vs. Prnp^ZH3/ZH3^* = 95.06 ± 9.26%, \*\**p* = 0.0021 Day 5: *Prnp^+/+^* = 226.3 ± 29.4 % *vs. Prnp^ZH3/ZH3^* = 84.7 ± 8.0 %, \**p* = 0.034; mean ± S.E.M.; two-way ANOVA + Bonferroni’s multiple comparisons test) (Fig. 4c). Following the same tendency, in *Prnp^ZH3/ZH3^* mice, fEPSP2 was also smaller than in *Prnp^+/+^* (Baseline: *Prnp^+/+^* = 268.24 ± 43.1 % *vs. Prnp^ZH3/ZH3^* = 158.4 ± 23.0 %, *p* > 0.99; Day 1: *Prnp^+/+^* = 441.1 ± 140.2 % *vs. Prnp^ZH3/ZH3^* = 224.3 ± 39.8 %, *p* = 0.370; Day 2: *Prnp^+/+^* = 554.80 ± 164.2 % *vs. Prnp^ZH3/ZH3^* = 181.4 ± 25.9 %, \*\**p* = 0.0083; Day 3: *Prnp^+/+^* = 466.3 ± 106.7 % *vs. Prnp^ZH3/ZH3^* = 160.39 ± 23.8 %, *p* = 0,051; Day 4: *Prnp^+/+^* = 438.12 ± 90.7 % *vs. Prnp^ZH3/ZH3^* = 139.73 ± 19.9 %, *p* = 0,062; Day 5: *Prnp^+/+^* = 433.1 ± 87.7 % *vs. Prnp^ZH3/ZH3^* = 133.53 ± 18.0 %, *p* = 0.06; mean ± S.E.M.; two-way ANOVA + Bonferroni’s multiple comparisons test) (Fig. 4d). These results indicate that LTP increased fEPSPs in *Prnp^+/+^* connections but not in *Prnp^ZH3/ZH3^* ones. These results were surprising and were not in accordance with previous publications in which *Prnp^ZH1/ZH1^* mice showed even exacerbated LTP [19]. In an attempt to explain these results, we checked the *in situ* registers in detail during the HFS protocol (Fig. 4g). We observed that 55 % of the *Prnp^ZH3/ZH3^* mice suffered from epileptic seizures due to HFS in contrast to 20 % of the *Prnp^+/+^* mice (Fig. 4h). The *Prnp^ZH3/ZH3^* epileptic crises tended to be longer (although not statistically significantly) than those suffered by *Prnp^+/+^* mice (*Prnp^+/+^* = 12.08 ± 3.3 s; *Prnp^ZH3/ZH3^* = 19.55 ± 2.5 s, mean ± S.E.M., *p* = 0.12; Mann-Whitney *U* non-parametric test) (Fig. 4i). We postulate that this exacerbated excitability in *Prnp^ZH3/ZH3^* synapse impaired a proper LTP generation. The HFS may bring about an aberrant synaptic activation (even generating epileptic seizures) that enables activation of the molecular mechanisms needed for LTP induction. Therefore, as published with chemoconvulsants models [28, 31], PrP^C^ might exert protection against electrically-induced seizures.

To gain insight into the gene expression patterns altered in the *Prnp^ZH3/ZH3^* mice, RNA-seq was performed from the hippocampus region of 8 *Prnp^+/+^* and 8 *Prnp^ZH3/ZH3^* animals. Around 700 genes showed alterations in their expression profile (323 upregulated and 390 downregulated in *Prnp^ZH3/ZH3^* compared to *Prnp^+/+^*) (Additional files 5 and 6: Table S1 and Table S2). According to pathway analysis in Reactome v77, the main alterations related to brain functions were the down-regulation of genes associated with the *“MECP2 regulates neuronal receptors and channels”* (10 genes; *p*adj = 1.06 E-07), the “*neuronal system”* (28 genes; *p*adj. = 0.002), and “*protein-protein interaction at synapses”* (10 genes; *p*adj. = 8.65 E-04) (Additional file 7: Fig. S5), all in line with the previous behavioral and electrophysiological findings. Among the dysregulated genes, we validated some by RT-qPCR that could explain the phenotype shown by the *Prnp^ZH3/ZH3^* mice: including the downregulation of glutamate ionotropic receptor NMDA type subunit 2B (*Grin2b*), the Gamma-Aminobutyric Acid Type A Receptor Subunit Rho2 (*Gabrr2*), the Potassium Voltage-Gated Channels: Subfamily J Member 2 (*Kcnj2*) and 6 (*Kcnj6*), Subfamily A Member 1 (*Kcna1*), and Subfamily Q member 3 (*Kcnq3*) (Additional file 7: Fig. S5).

### Enhanced susceptibility to KA-induced seizures in Prnp^ZH3/ZH3^ mice correlates with neuronal death in the hippocampus

Next, we aimed to explore whether the absence of PrP^C^ in the *Prnp^ZH3/ZH3^* mice increased their susceptibility to epileptic seizures following KA (i.p.) injections, as reported in *Prnp^ZH1/ZH1^* mice (B6129 and B6.129 backgrounds) [28] (Fig. 5). All experiments were carried out on a blind basis, and two different researchers carried out data evaluation (see Methods). Three consecutive injections of KA (10 mg/Kg b.w.) were administrated at intervals of 30 min. The epileptic behavior was monitored for 3 h and was categorized into six stages according to its severity (Fig. 5a). Results indicate that 67 % of *Prnp^+/+^* mice did not suffer any severe epileptic episodes (stage I-IV). Only 22 % and 11 % of wild-type mice reached stage V and VI, respectively. In contrast, 55% of *Prnp^ZH3/ZH3^* mice suffered severe epileptic episodes, 20 % reaching stage V and 35 % at stage VI (Fig. 5a and Additional file 8: Movie. S1). Moreover, *Prnp^ZH3/ZH3^* mice presented more seizures and blinking episodes per animal than *Prnp^+/+^* individuals (Seizure: *Prnp^ZH3/ZH3^* = 2.45 ± 0.74 *vs. Prnp^+/+^* = 1.06 ± 0.83; *p* = 0.019; Blinking: *Prnp^ZH3/ZH3^* = 0.95 ± 0.29 *vs. Prnp^+/+^* = 0.22 ± 0.13; mean ± S.E.M. *p* = 0.069; Mann-Whitney *U* non-parametric test) (Fig. 5b).

**Fig. 5.**
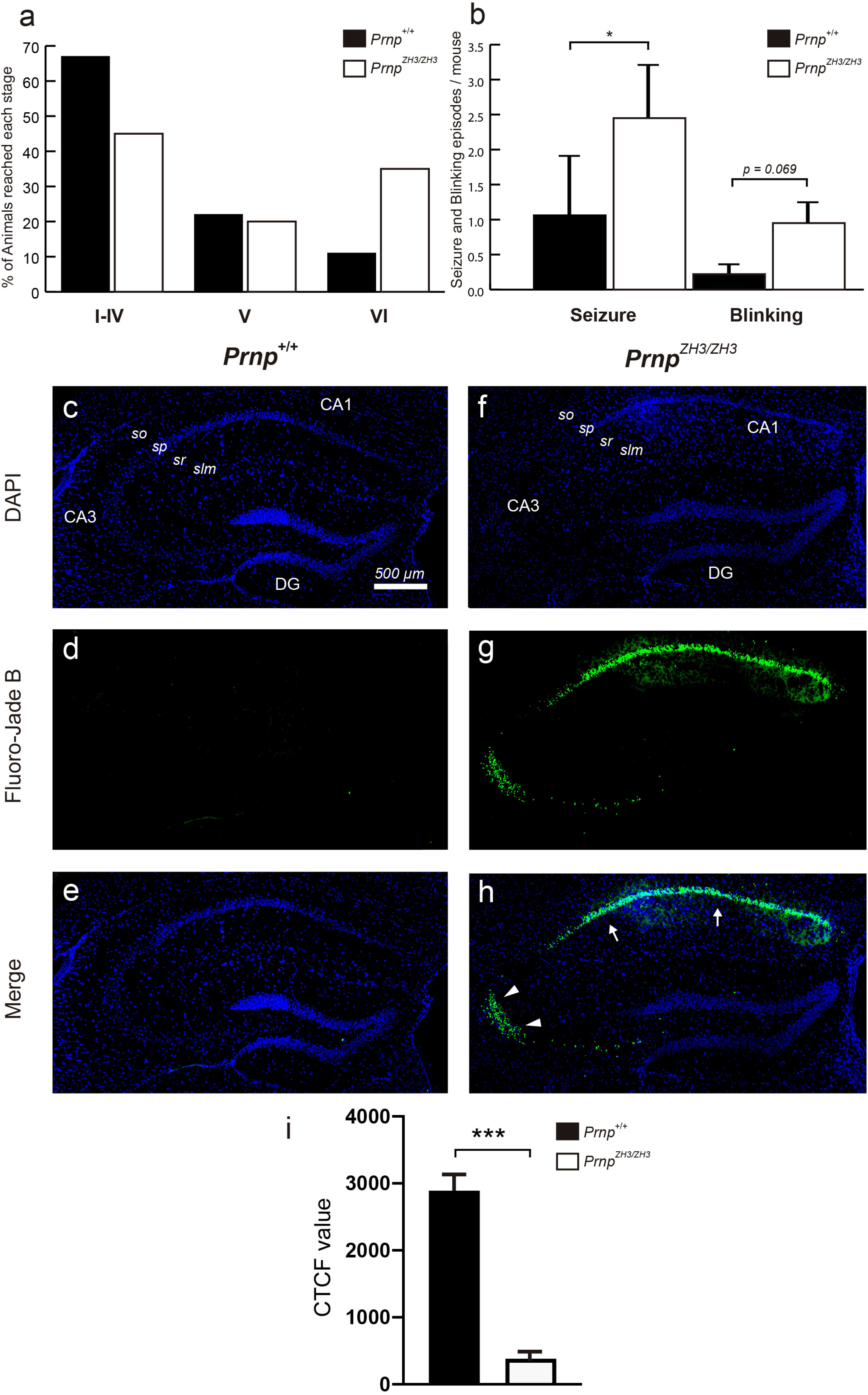
*Prnp^ZH3/ZH3^* mice are more susceptible to KA-induced epilepsy correlating with increased neuronal death in CA1 and CA3 pyramidal layers. ***a***, Percentage of mice reaching stage I-IV, V, or VI epileptic phenotype after KA administration (10 mg/Kg). ***b***, Number of seizures and blinking episodes presented by *Prnp^+/+^* and *Prnp^ZH3/ZH3^* mice for 3 h after KA administration. ***c-h***, Photomicrographs showing the pattern of neurodegeneration with Fluoro-Jade B staining seven days after KA treatment in *Prnp^+/+^* (***c****-**e***) and *Prnp^ZH3/ZH3^* (***f-h***) mouse hippocampus. Nuclei are stained with DAPI (***c****, **f***). Dying cells (***d****, **g***, stained with Fluoro-Jade B) are located in the pyramidal cell layer of CA1 (arrows) and CA3 (arrowheads) areas. ***i***, Graph illustrating the analysis of the CTCF values in the CA1-3 pyramidal layer of *Prnp^ZH3/ZH3^* and wild-type mice (see Methods for details). Data are presented as a percentage in ***a*** and as mean ± S.E.M, in ***b*** and ***i***; \**p* < 0.05 and \*\*\**p* < 0.001, Mann–Whitney *U* non-parametric test. Abbreviations: so, *stratum oriens*; sp, *stratum pyramidale*; sr, *stratum radiatum*; slm, *stratum lacunosum-moleculare*; DG, dentate gyrus. The scale bar in (***c***) is also representative for *(**d-h**)*.

Additionally, we evaluated neuronal damage after KA-induced epilepsy with Fluoro-Jade B (Fig. 5c-h). *Prnp^ZH3/ZH3^* mice showed relevant numbers of labeled cells in the pyramidal layer of the CA1 and CA3 (Fig. 5c-e), while no signal was observed in *Prnp^+/+^* sections (Fig. 5f-h). Indeed, CTCF analysis of Fluoro-Jade B labeling in the pyramidal layer of the CA1 and CA3 (see Methods for details) demonstrated statistical differences between *Prnp^ZH3/ZH3^ vs* wild-type mice: CTCF value for *Prnp^ZH3/ZH3^* = 2893 ± 349.3 *vs*. wild-type = 380 ± 84.05; mean ± S.E.M., *** *p* < 0,001; Mann–Whitney *U* non-parametric test (Fig. 5i). These results corroborated the absence of *Prnp* to generate an exacerbated synaptic excitability in the hippocampal region that increases susceptibility to electrical and KA-induced seizures, causing neuronal death in the CA1 and CA3 regions of the pyramidal layer of the hippocampus proper.

### Neuronal Prnp^ZH3/ZH3^-derived cultures show reduced bursting and impairment network formation in vitro

PrP^C^ has also been described as a regulator of neurogenesis and neuronal differentiation *in vitro* and *in vivo* (see [6–8] for reviews). Furthermore, defects in neuronal network connectivity and maturation are related to epilepsy [59]. Consequently, we tested whether *Prnp^ZH3/ZH3^* increased excitability might be due to changes in the neuronal differentiation inducing aberrant connectivity and an immature neuronal network. To analyze this, calcium imaging was performed in primary cortical cultures (n = 10 in both genotypes) from *Prnp^+/+^* and *Prnp^ZH3/ZH3^* mouse embryos (E16.5-E17.5) expressing the GECI indicator GCaMP6f under the neuronal syntaxin promoter [60], allowing us to record calcium traces of the same neuronal population after 8, 11, 13, and 15 days *in vitro* (DIV) (Fig. 6).

**Fig. 6.**
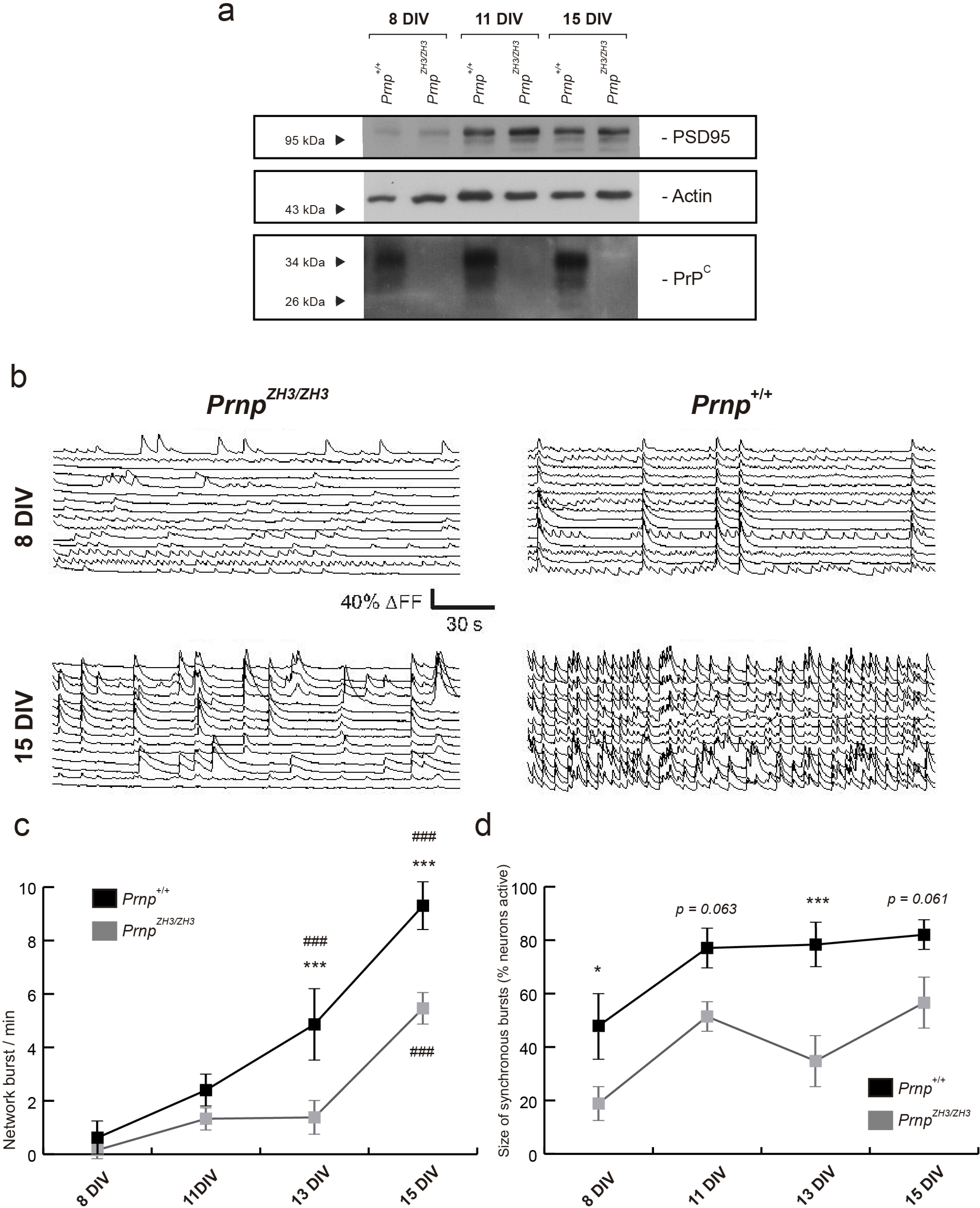
Reduced bursting and network formation in neuronal *Prnp^ZH3/ZH3^*-derived cultures. ***a***, Immunoblot analysis of PrP^C^ expression and PSD95 in *Prnp^+/+^* and *Prnp^ZH3/ZH3^* derived primary neuronal cultures after 8, 11, and 15 DIV. Note the absence of PrP^C^ in the *Prnp^ZH3/ZH3^* cultures and the same PSD95 expression in each DIV. Actin is used as a loading control. ***b***, Representative examples of neuronal traces at 8 and 15 DIV in the *Prnp^+/+^* and *Prnp^ZH3/ZH3^* primary cultures. Note the asynchrony in the *Prnp^ZH3/ZH3^* culture. ***c***, Evolution of network bursting in *Prnp^+/+^* and *Prnp^ZH3/ZH3^* neuronal cultures from 8 to 15 DIV. Data are presented as the mean of bursts/min ± S.E.M. ***d***, Evolution of size of synchronous bursts from 8 to 15 DIV. Data are presented as the mean percentage of active neurons ± S.E.M. Asterisks (*) indicate significant differences between *Prnp^+/+^* and *Prnp^ZH3/ZH3^* bursting. Number sign (#) indicates significant differences with the respective baseline bursting at 8 DIV. \**p* < 0.05, \*\*\**p* < 0.001 and ^###^*p* < 0.001, two-way ANOVA + Bonferroni’s multiple comparisons test.

*Prnp^+/+^* and *Prnp^ZH3/ZH3^* cultures displayed the same number of collective bursts/min at 8 and 11 DIV (Fig. 6c). After that, a delay in *Prnp^ZH3/ZH3^* neuron activity was observed compared to controls. *Prnp^+/+^* cultures increased the number of bursts/min significantly at 13 DIV; however, *Prnp^ZH3/ZH3^* cultures needed two additional days, at 15 DIV (Fig. 6b-c). Moreover, *Prnp^+/+^* neurons exhibited significantly more bursts/min at 13 DIV and 15 DIV than *Prnp^ZH3/ZH3^* ones and overall the latter showed a reduced firing interval along development (8 Div: *Prnp^+/+^* = 0.62 ± 0.4 *vs. Prnp^ZH3/ZH3^* = 0.16 ± 0.1 bursts/min; 13 Div: *Prnp^+/+^* = 4.86 ± 1.1 *vs. Prnp^ZH3/ZH3^* = 1.38 ± 0.4; \*\*\**p* < 0.001; 15 Div: *Prnp^+/+^* = 9.30 ± 0.7 *vs. Prnp^ZH3/ZH3^* = 5.46 ± 0.4; mean ± S.E.M. \*\*\**p* < 0.001; ANOVA + Bonferroni’s multiple comparisons test) (Fig. 6c). The *Prnp^ZH3/ZH3^* cultures also showed a reduction in the size of synchronous bursts. In the *Prnp^+/+^* cultures, around 80 % of the neurons showed synchronic activity at 11, 13, and 15 DIV, while this value was around 50 % in *Prnp^ZH3/ZH3^* cultures (Fig. 6d). These results demonstrate that collective bursting is reduced and delayed in *Prnp^ZH3/ZH3^* cultures, suggesting that *Prnp* expression is necessary for network formation and maturation.

## Discussion

PrP^C^ has been associated with several physiological functions using *in vivo* approaches; however, the consequences of PrP^C^ deletion in behavior and cognition have not been extensively evaluated [9]. There are some studies about *Prnp^0/0^* mouse behavior, motor capabilities, and learning performance, but the results are not conclusive, especially after the description of the so-called FG in the background of the *Prnp^ZH1/ZH1^* model that masks specific PrP^C^ roles [26]. For example, concerning KA susceptibility, a clear decrease is observed in mice expressing a lower percentage of 129/Sv-associated polymorphisms (B6.129) compared to the B6129 original strain of the *Prnp^ZH1/ZH1^* mice with a higher percentage of 129/Sv genome [28]. Here we assess the consequences of the absence of PrP^C^ in behavior and neurotransmission using the new strictly co-isogenic mouse model *Prnp^ZH3/ZH3^* [50]. However, another relevant aspect of these studies is the age of the analyzed mice, since physiological differences in the absence of *Prnp* have been described in association with age for *Prnp^ZH1/ZH1^* [45] or FVB/N-*Prnp^ZH1/ZH1^* [61]. Thus, in our study, we only used and compared results obtained from mice of 3-5 months of age. Concerning nest-building behavior, our results suggest similar behavior in *Prnp^ZH3/ZH3^* and wild-type mice. This contrasts with previously reported results [42], but it has been largely demonstrated that this capacity is dependent on mouse background [62, 63].

Our results also reveal that *Prnp^ZH3/ZH3^* mice display reduced activity in the open field test. The reduced rearing exploration and peripheral preference in the arena, and the high defecation rate observed in *Prnp^ZH3/ZH3^* mice, suggest an anxiety-like behavior. Indeed, *Prnp^ZH3/ZH3^* showed higher thigmotaxis than wild-type mice (see [64] for technical details). Our data correlate with those reported by Schmitz *et al.* [45] and Lobao-Soares *et al.*, both using 3-month-old *Prnp^ZH1/ZH1^* mice, illustrating reduced mobility between the inner and outer regions of the open field and increased defecation in *Prnp^ZH1/ZH1^* mice [65]. However, they are in contrast to Nico *et al.*, [66], where they found no differences, and with Gadotti *et al.* [52], that showed ten-week-old *Prnp^ZH1/ZH1^* mice displayed increased crossing in open-field test although thigmotaxis changes were not analyzed in the study. Although technical details could also play a role (i.e., handling of the mice, the initial position of the mice in the field, geometry of the field, etc.), we think that these discrepancies reinforce the relevance of the homogenous genetic background in our study *vs.* the others. In fact, the differences we observed also extend to the rotarod test. Our results showed no motor alternations in the *Prnp^ZH3/ZH3^* mice, but they need more sessions to learn the task compared to controls. Previous publications also showed any motor alterations in two different knock-out strains [45, 65, 67]. However, Nazor *et al*., observed changes just in mice older than 95 days [61].

In our experiments, *Prnp^ZH3/ZH3^* mice also failed to achieve instrumental learning in the Skinner box. These results are similar to those observed in *Prnp^ZH1/ZH1^* mice (B6.129 background, 2-5 % 129/Sv markers, [28]). Striking differences in motility between wild-type and knock-out mice were identified using this approach. However, learning capacity based on the ON/OFF paradigm confirmed the operant conditioning deficiencies, a type of associative learning.

Following our results, alterations in locomotor activity and increased latency to initiate exploration were previously reported in other *Prnp^0/0^* mouse models [42, 47]. Anxiety-related behavior [54], depressive tendencies [52], and alterations in spatial memory and learning have also been described [42, 45]. In contrast, Bueler *et al.* reported no alterations in *Prnp^ZH1/ZH1^* behavior [24]. This disparity in results might be explained by the age of the animals used in the studies. Bueler and collaborators performed the test with 2- and 3-month-old mice, which could potentially uncover the behavior impairment as it was reported to be an age-dependent decline in other publications using the *Prnp^ZH1/ZH1^* model [45, 68]. Another study also showed no deficits in *Prnp^0/0^* mouse behavior, where *Prnp* was conditionally deleted at 12 or 16 months, and the results included no alterations in the Morris water maze or object recognition test [53]. We reported behavioral deficits in *Prnp^0/0^* mouse models with some discrepancies with the previous works, most likely related to age-related sampling and the mice’s background.

Glutamate neurotransmission is, in large part, responsible for cortical signaling, and its impairment has been related to behavioral deficits [69]. PrP^C^ has been described as a regulator of glutamate synapses [13]. Even glutamate inhibition with an NMDA antagonist (MK-801) ameliorates depressive-like behavior in *Prnp^0/0^* mice [52]. Thus, our next step was to study glutamate connectivity to better understand behavioral alterations. The hippocampal Schaffer collaterals and their implication in operant conditioning, spatial learning, and anxiety-related behavior were evaluated as a well-defined model of glutamate circuitry [70].

The paired-pulse facilitation test did not show differences between *Prnp^ZH3/ZH3^* and *Prnp^+/+^* animals. Therefore, PrP^C^ deletion did not alter synaptic facilitation, at least in the living behaving mouse model, at the least at relatively low stimulation intensities (Fig. 3b). These results may be explained by the fact that synaptic facilitation is mainly a presynaptic phenomenon [71], and PrP^C^ has been related to postsynaptic neurotransmission mechanisms [37, 38]. These differences in pre- and post-synaptic mechanisms would explain the different results collected from *Prnp^ZH3/ZH3^* mice and their littermate controls in paired-pulse ratio and LTP tests.

Nevertheless, *Prnp^ZH3/ZH3^* mice displayed increased synaptic facilitation at high intensities, a fact not observed in controls. This could represent a sort of compensatory phenomenon for their evident LTP deficits. In addition, *Prnp^ZH3/ZH3^* mice presented an increased susceptibility to KA-induced seizures. This epileptogenic phenotype may explain our results on anxiety behavior in *Prnp^ZH3/ZH3^* mice. Comorbid anxiety disorders affect patients with epilepsy [72], and cognitive decline has been described in epileptic animal models [73]. Moreover, increased excitability was previously reported, especially in susceptibility to KA, NMDA, and pentylenetetrazol (PTZ) insults [30, 31].

However, contradictory results were published by other groups, who described an elevated epileptic threshold in *Prnp^0/0^* hippocampal slices treated with bicuculline, zero-magnesium conditions, and PTZ [74], and also normal neurotransmission-associated parameters compared with wild-type mice [25]. Both studies recorded hippocampal slices of the FVB/N-*Prnp^0/0^* model, a mouse with a triple mixture background (FVB/129Sv/C57BL6) which carried the FG [26]. Different susceptibility to KA-induced seizures among mice with different backgrounds has been described; even the FVB background has been shown as highly susceptible to epilepsy [75, 76]. Our results using *Prnp^ZH3/ZH3^* animals in KA susceptibility were similar to those previously obtained with the other available *Prnp^0/0^* co-isogenic mouse model, the Edinburgh 129/Ola [28]. Therefore, we postulate that these contradictory results published about the excitability of *Prnp^0/0^* synapse are likely associated with the mouse backgrounds, the FG effect, and the experimental approach. However, the ZH3 mice results demonstrate that PrP^C^ indeed protects against KA-induced epilepsy.

Our results show that the presentation of HFS protocols causes epileptic seizures in most *Prnp^ZH3/ZH3^* mice but fails to generate significant LTP at the hippocampus. Some controversial results have been published about the implication of PrP^C^ in LTP generation. Different experimental approaches (hippocampal slices or *in vivo* experiments) and a mixture of mouse models with distinct backgrounds were used, generating non-comparable data (i.e., [19, 25, 40, 42]). Here, using ZH3 mice, we hypothesize that the absence of PrP^C^ ends in LTP induction failure due to exacerbated synaptic excitability, although we cannot rule out putative GABAergic disinhibition. It is convincingly demonstrated that severe epileptic seizures cause neuronal death, which hampers LTP generation. Moreover, non-severe epileptic seizures generate similar molecular and synaptic changes to LTP [77, 78]. This suggests that non-severe HFS-induced seizures somehow saturate the postsynaptic terminal, over-activating LTP-induction mechanisms that reduce LTP production capacity by HFS. Additionally, PrP^C^ has also been described as interacting with key elements required for LTP-related mechanisms AMPA or NMDA receptors [37, 39, 79, 80], see also [57] for review.

The presented data show that *Prnp^0/0^* hippocampal synapse is highly excitable and epileptogenic. Alterations in brain connectivity due to developmental alterations, traumas, or infections contribute to this neuromodulation imbalance [59]. In order to assess whether the epileptic phenotype displayed by *Prnp^ZH3/ZH3^* animals came from neuronal connectivity alterations, we studied bursting and network formation in *Prnp^ZH3/ZH3^-*derived primary neuronal cultures. Relevant PrP^C^ expression *in vitro* is observed from 4-5 DIV [81]. Our results indicate that *Prnp^ZH3/ZH3^* cultures did not mature or connect properly; they displayed asynchronous and very low bursting compared to wild-type cultures. These results suggest that the absence of PrP^C^ causes a delay in neuronal maturation, but more relevantly in neural network formation and function. In fact, the role of PrP^C^-mediated signals in neuritogenesis has been demonstrated [82–84]. However, to our knowledge, this is the first description of network alterations due to the absence of PrP^C^. However, and as indicated by Benvegnu and coworkers [85], there are gene expression changes during the development of FVB/N *Prnp^0/0^* and wild-type hippocampus. Hence, changes in ion conductance or channel receptor expression might be involved in this delay. In this line, electrophysiological experiments with biochemical characterization might confirm the basis of the delay in maturation *in vitro*.

Finally, the gene ontology analysis of the RNAseq of *Prnp^ZH3/ZH3^* and *Prnp^+/+^* mouse hippocampus showed down-regulation of genes associated with the neuronal system and protein-protein interaction at synapses, fitting the phenotype we observed in the knock-out mice. The significant downregulation of *Gabrr2* and *Grin2b* in *Prnp^ZH3/ZH3^* mice would produce dysregulation in the excitatory/inhibitory balance, increasing the excitability of the system, as we describe with the KA and the HFS analyses. The alteration of the inhibitory neurotransmission was already shown in *Prnp^0/0^* models since they were susceptible to suffering from epileptic crises (see references above). PrP^C^ has been widely described as a regulator of glutamatergic neurotransmission and its receptors, as we show here with the *Grin2b*. Moreover, mutations at *Grin2b* have recently been related to a rare brain disease, the *GRIN2B*-related neurodevelopmental disorder that causes intellectual disability, autism-spectrum-like behavior, epilepsy, and, sometimes, locomotor deficiencies as well [86, 87]. Therefore, the altered expression of *Grin2b* might contribute to the behavior and learning deficits observed in the *Prnp^ZH3/ZH3^* mice.

## Conclusions

In conclusion, our study points that the absence of PrP^C^ impairs neuronal network formation and connectivity, producing enhanced susceptibility to excitotoxicity insults such as HFS and KA exposure. This epileptogenic circuitry seems to impair highly cognitive-demanding functions such as associative learning, and it produces anxiety-like behavior.

## Methods

### Animals

Adult C57BL/6J mice (*Prnp^+/+^*) were purchased from Charles River Laboratories (Paris, France). *Prnp^ZH3/ZH3^* mice line was provided by A. Aguzzi (Switzerland) (see [50] for details). *Prnp^ZH1/ZH1^* mice [24] were purchased from the European Mouse Mutant Archive (EMMA, Monterotondo, Italy). A total of 185 adult (3-5 months old) male mice (ZH3: *Prnp^+/+^*= 81 and *Prnp^ZH3/ZH3^*= 84; ZH1: *Prnp^+/+^*= 10 and *Prnp^ZH1/ZH1^* = 10) were used in the present study. In ZH1 mouse experiments, null *Prnp^ZH1/ZH1^* and control mice (*Prnp^+/+^*) were obtained by crossing heterozygous *Prnp^+/ZH1^* mice to obtain a mixed background (B6.129). It is well described that behavior and neural physiology are different between male and female rodent models due to several hormone and non-hormone-derived reasons [88]. Thus, we used only males in order to establish an equivalent group comparable with previous publications. All experiments were performed following the protocols and guidelines of the Ethical Committee for Animal Experimentation (CEEA) of the University of Barcelona. CEEA of the University of Barcelona approved the protocol for using animals in this study (CEEA approval #276/16 and 141/15). Behavioral and electrophysiological studies were performed following the guidelines of the European Union Council (2010/276:33-79/EU) and current Spanish regulations (BOE 34:11370-421, 2013) for the use of laboratory animals in chronic experiments. Experiments were also approved by the Ethics Committee for Animal Care and Handling of the Pablo de Olavide University (UPO-JA 06/03/2018/025).

### Immunoblotting

Proteins from brain tissue lysates or primary cortical neurons were extracted using RIPA buffer with protease and phosphatase inhibitor cocktails (Roche). Total lysates were obtained by 30 sec. of centrifugation at 4°C. The protein concentration of the lysates was quantified using Pierce BCA Protein Assay Kit (Thermo Scientific). 10-50μg of proteins were loaded to SDS-PAGE gels and transferred and transferred to nitrocellulose membranes for 1 h. Membranes were blocked with Tris-buffered solution with 0.1 % tween, 5 % skimmed milk, and 2 % of FBS for 1 h at room temperature (RT) and incubated with PSD95 (1:1.000, MAB1598; Millipore), PrP (1:500; 6H4; Thermo) or Actin (1:20.000;MAB1501; Millipore) antibodies at 4°C O/N. Following HRP-linked secondary antibody (Dako) incubation for 1h at RT, membranes were developed with ECL substrate (Thermo).

### Behavioral studies

A total of 147 animals were used in these sets of experiments (ZH3: *Prnp^+/+^*= 63 and *Prnp^ZH3/ZH3^* = 64; ZH1: *Prnp^+/+^*= 10 and *Prnp^ZH1/ ZH1^* = 10). Mice were housed alone in boxes on a 12/12 h light/dark cycle with constant ambient temperature (21 ± 1 °C). Water and food were provided *ad libitum* except for the instrumental learning tests (see below).

### Nest building

For this test, a total of 14 mice (3 months old) were used (*Prnp^ZH3/ZH3^* = 7 and *Prnp^+/+^* = 7). On the first day of testing, one piece of tissue paper (36 x 12 cm) was placed in the cage to facilitate nest building (Additional file 1: Fig. S1). The presence and the quality of each nest were photo-documented and evaluated the following day according to a modified five-point scale using the method described by Deacon [89]. Two different blinded researchers evaluated the nest generated by each mouse. Data are presented as the mean ± S.E.M. in (Additional file 1: Fig. S1). The statistical analysis was performed with the Mann-Whitney non-parametric test (GraphPad Prism 8 software).

### Open field test

In this test, *Prnp^ZH1/ZH1^* mice were not used since detailed studies were already developed using this model [45, 54]. In our experiments, mice (*Prnp^+/+^* = 49 and *Prnp^ZH3/ZH3^* = 49) were placed in a square open field altimeter box (35 × 35 × 25 cm, Cibertec, Madrid, Spain). The field had a grid (16 x 16 cm) of infrared lasers on the XY axis and one on the Z-axis. Locomotor activity was measured for 15 min in mice with the MUX-XYZ16L software. Mice were placed in the box’s periphery for 15 min for two consecutive days, and their behavior was recorded. The first day was considered a training session to reduce mouse anxiety associated with manual handling, and the data analyzed and displayed in the manuscript corresponded to the second session. The system inferred mouse activity by counting laser intersections. For anxiety-related behavior measurement, the center (inner) square of the field (10 x 10 cm) was considered as the central zone and the rest of the square as the peripheral (outer) zone [56] (see Fig. 1a). For quantification and to distinguish motility from exploratory behavior, it was considered that a mouse spent time in one of the regions (center *vs.* periphery) if it remained in the region at least 3 s. Rearing episodes were considered when the animal stood up for at least 3 s, and immobility episodes if immobile for an additional 3 s. Obtained data were analyzed, and the sum of the crossed X and the Y axes are presented together to show total mouse mobility in the experiments. The time spent in the maze periphery zones measures thigmotaxis or wall-hugging behavior and indicates anxiety-related behavior [56]. Data are presented as the mean ± S.E.M. The statistical analysis was performed with a *T*-test or Mann-Whitney *U* non-parametric test (GraphPad Prism 8 software). The asterisks indicate significant differences: \*\**p* < 0.01 and \*\*\**p* < 0.001. The arena and the walls were cleaned with soap and ethanol between trials to remove olfactory cues between experiments.

### Operant conditioning tests

The instrumental learning tests were performed as described in previous studies of our group [90]. Six Skinner boxes were used simultaneously (12.5 x 13.5 x 18.5 cm; MED Associates, St. Albans, VT, USA). Each Skinner box was housed in a sound-attenuating cubicle (90 x 55 x 60 cm) constantly exposed to white noise (± 45 dB) and dim light (Cibertec, S.A, Madrid, Spain). The boxes had a trough to receive food pellets (Noyes formula P; 45 mg; Sandown Scientific, Hampton, UK) by pressing a lever. Before the test, mouse food availability was monitored for seven days to reduce initial mouse weight to 85 %. First, mice (ZH3: *Prnp^+/+^* = 49 and *Prnp^ZH3/ZH3^* = 49; and ZH1: *Prnp^+/+^* = 10 and *Prnp^ZH1/ZH1^* = 10) were trained to press the lever to receive food pellets in a fixed-ratio (1:1) schedule. Seven daily sessions (20 min/each) were held. The boxes were cleaned with soap and ethanol (30 %) between trials. Obtaining ≥ 20 pellets for two consecutive sessions was defined as the criterion to assume the learning criteria achievement. Following this first operant conditioning test, we increased the paradigm complexity to test the mice in a more demanding cognitive task for an additional 10 days. Only animals that met the learning criterion were tested (ZH3: *Prnp^+/+^* = 24 and *Prnp^ZH3/ZH3^* = 20; and ZH1: *Prnp^+/+^* = 8 and *Prnp^ZH1/ZH1^* = 5). The paradigm consisted of light (ON period) and dark periods (OFF period) randomly distributed during the session. The light was provided by a small light bulb located over the lever. During the ON period (20 s), lever presses were reinforced with food pellets at a ratio of 1:1. During the OFF period, lever presses were not rewarded and were penalized by adding ten additional seconds (20 ± 10 s) to the next ON period. The number of lever presses during the different conditioning paradigms was monitored and recorded with the MED-PC program (MED Associates, St. Albans, VT, USA). Statistical analysis was carried out using two-way ANOVA with repeated measures and Bonferroni’s multiple comparisons test (GraphPad Prism 8 software). Asterisks indicate significant differences: \*\**p* < 0.01; and \*\*\**p* < 0.001. Data are presented as the mean ± S.E.M. or as a percentage (as indicated in each Figure).

### Rotarod test

For this test, a total of 15 mice of 4 months were used (*Prnp^ZH3/ZH3^* = 8 and *Prnp^+/+^* = 7). Motor performance was tested using an accelerating Rotarod. Mice were pre-trained to the task to reach a minimum of 30 sec. performance at 5 rpms on the 1st day of testing. In each training run, animals were placed on the rods at an initial speed of 5 rpm for 30 sec. After that, the testing consisted of 5 consecutive trials with 15 min. inter-trial intervals. Each trial consisted of 30 sec. at 5 rpm followed by 5 rpm increases every 15 sec. with a cut-off of 5 min. Results are expressed as the mean latency of animals to fall from the rod ± S.E.M. The statistical analysis was performed with the two-way ANOVA + Bonferroni’s multiple comparisons test (GraphPad Prism 8 software).

### Object recognition test

The object recognition test was performed in a homemade arena (30 x 25 x 20 cm), as described [91]. A total of 23 *Prnp^+/+^* and 24 *Prnp^ZH3/ZH3^* mice were analyzed. Additionally, 6 *Prnp^ZH1/ZH1^* mice were also used with 7 *Prnp^+/+^* counterparts. The test consisted of four phases of 10 min/each. First, animals were habituated to the field without any object (habituation session). One hour later, two identical plastic objects were placed in the center of the arena for the training session. A short-term memory test was performed 2-3 h later by changing one of the objects (see Additional File 4: Fig. S4). ZH1 mouse mobility was expressed as the number of rearing episodes during the habituation session. The arena and the objects were cleaned with soap and 30 % ethanol between trials to remove olfactory cues. Mouse behavior was recorded with a video camera placed over the arena, and these recordings were used to measure the exploratory behavior blindly. Sniffing and gently touching the objects were counted as exploratory behavior. To further support increased anxiety levels in the *Prnp^ZH3/ZH3^* mice, fecal bodies left in the maze during the habituation session were counted by the observer once the test subject was removed since it has been demonstrated that highly emotional animals exhibit increased defecation [56]. Statistical analysis was performed with the Mann-Whitney *U* non-parametric test (GraphPad Prism 8 software). Data are presented as the mean ± S.E.M. or as a percentage (indicated in each Figure).

### Mouse surgery

A total of 98 adult male (3-5 months) mice were implanted with stimulating and recording electrodes (*Prnp^+/+^* = 49 and *Prnp^ZH3/ZH3^* = 49). Four of them died during surgery, and 33 mice were excluded because of the inability to obtain reliable and clean recordings. Thus, the experiments were performed with 61 mice (*Prnp^+/+^* = 31 and *Prnp^ZH3/ZH3^* = 30). Surgery was performed as described in [19, 92]. Mice were deeply anesthetized with ketamine (35 mg/kg) and xylazine (2 mg/kg), and electrodes were aimed at the right dorsal hippocampus. Two recording electrodes were implanted in the *stratum radiatum* of the CA1 area (2.2 mm caudal to Bregma, 1.2 mm lateral, and 1.3 mm ventral), and two stimulating electrodes were implanted in the Schaffer collateral pathway of the CA3 region (1.5 mm posterior to Bregma, 2 mm lateral, and 1.3 mm ventral). Electrodes were made of 50 µm Teflon-coated tungsten wires (Advent Research, Eynsham, UK). Electrode localizations were checked according to the field excitatory postsynaptic potential (fEPSP) profile evoked by a single stimulation. A silver wire was fixed to the skull as ground. All the wires were soldered to a six-pin socket (RS Amidata, Madrid, Spain) fixed to the skull with dental cement. Recordings were started at a minimum of one week after the surgery.

### Electrophysiology recordings

Animals were consecutively recorded in groups of six individuals since they reach the total number of animals used in each experiment. Each animal was placed in a small plastic cubicle (5 x 5 x 10 cm) inside a large Faraday box (30 x 30 x 20 cm). fEPSPs were recorded with a high impedance probe (2 x 10^12^ Ω, 10 pF) using differential amplification at a bandwidth of 0.1 Hz-10 kHz (P511, Grass-Telefactor, West Warwick, RI, USA). For each experiment, artefactual recordings were discarded. The stimulation intensity threshold of each animal was set with paired-pulse stimulations at 40 ms of inter-stimulus interval. The stimulus intensity was set to 40-60 % of the amount necessary to evoke a suturing response. These intensity values were used for all the experiments.

### Paired-pulse stimulation

For synaptic facilitation experiments, 51 mice (*Prnp^+/+^* = 27 and *Prnp^ZH3/ZH3^* = 24) were stimulated at Schaffer collaterals with a pair of pulses at different inter-stimulus intervals (10, 20, 40, 100, 200, and 500 ms) at 2 × Threshold intensities (≈ 0.2 mA). Threshold values were previously defined for each mouse. As classically defined, threshold values were determined as the intensity evoking fEPSP responses in 50 % of the cases. For all the inter-pulse intervals, the stimulations were repeated ten times. Data are represented as the mean percentage increases of fEPSP2 from fEPSP1 recordings (fEPSP2 / fEPSP1 x 100) ± S.E.M.

### Input/output curves

Schaffer collaterals of 29 mice (*Prnp^+/+^* = 14 and *Prnp^ZH3/ZH3^* = 15) were stimulated with paired pulses at 20 increasing intensities (from 0.02 mA to 0.4 mA, increased in steps of 0.02 mA) at 40 ms of inter-stimulus interval. For all the selected intensities, the stimulations were repeated ten times. Data are represented as the mean of fEPSP slopes (V/s) ± S.E.M. The same data are presented as the mean of paired-pulsed ratio (PP ratio) ± S.E.M. PP ratio is the percentage of the increase of the fEPSP2 from fEPSP1 recordings (fEPSP2 / fEPSP1 x 100). The area under the curve (AUC) was calculated from the PP ratio of all the animals using GraphPad Prism 8 software. Statistical analysis was performed using the Mann-Whitney-Wilcoxon non-parametric test or two-way ANOVA + Bonferroni’s multiple comparisons test (GraphPad Prism 8 software). The asterisks indicate significant differences: \**p* < 0.05; \*\**p* < 0.01; and \*\*\**p* < 0.001 in the Figure.

### Long-term potentiation experiments

For long-term potentiation experiments, 40 mice (*Prnp^+/+^* = 20 and *Prnp^ZH3/ZH3^* = 20) were stimulated at Schaffer collaterals. In a first experimental step, fEPSP baseline values were evoked and recorded for 15 min, with paired-pulse stimulus presented every 20 s (40 ms inter-stimulus). Stimulus intensities were selected to evoke fEPSPs of about 0.2-0.3 mV of amplitude (see insets in Fig. 4a,b). Next, LTP was evoked with a high-frequency stimulation (HFS) protocol. HFS consisted of five trains of pulses at a rate of 1/s (200 Hz, 100 ms) with the same intensity as the baseline recording. The HFS was repeated six times at intervals of 1 min. After the HFS protocol, fEPSPs were recorded, as for baseline, for 1 h. The following four days, the recordings were repeated for 30 min. fEPSPs and 1 V rectangular pulses corresponding to stimulus presentations were saved on a PC using an analog/digital converter (CED 1401 Plus, Cambridge, England). Data were analyzed offline using Spike2 and Signal 5.04 software with homemade representation programs [58]. Collected recordings were represented and analysed offline with the help of commertial (Spike 2 and Signal 5.04) programs following procedures described elsewhere. The slope of collected fEPSPs was computed as its first derivative (volts/s). Five successive fEPSs were averaged and the mean value of the slope was determined. Data are presented as the mean of the percentage compared to the baseline ± S.E.M. The statistical analysis was performed using two-way ANOVA + Bonferroni’s multiple comparisons test (GraphPad Prism 8 software). The asterisks and symbols indicate significant differences: \**p* < 0.05; \*\**p* < 0.01; and \*\*\**p* < 0.001; ^##^*p* < 0.01; and ^###^*p* < 0.001.

### KA-induced epilepsy and seizure analysis

Adult (3-4 months old) male mice were used for these sets of experiments (*Prnp^+/+^* = 18 and *Prnp^ZH3/ZH3^* = 20) essentially as described in [28]. A KA (Sigma-Aldrich, Darmstadt, Germany) solution was freshly prepared for each experiment in 0.1 M phosphate buffer. Mice were injected with KA (10 mg/kg b.w.) three times: at 0 min, 30 min, and 60 min. After the first injection, mice were placed in clean boxes (1-3 mice/box). The presence of epileptic seizures was monitored *in situ* and recorded with a video camera for 3 h after drug administration. Seizure severity was scored in grades following the following criteria: grade I-II: hypoactivity and immobility; grade III-IV: hyperactivity and scratching; grade V: loss of balance control and intermittent convulsions; grade VI: continuous seizures and bouncing activity (also reported as blinking episodes or ”*pop-corn*“ behavior). Data are presented as the mean ± S.E.M. or as a percentage (as indicated in each Figure). Statistical analysis was performed with the Mann-Whitney *U* non-parametric test (GraphPad Prism 8 software). The asterisk indicates significant differences: \**p* < 0.05 in the Figure.

### RNAseq

Libraries were prepared using the TruSeq Stranded mRNA Sample Prep Kit v2 according to the manufacturer’s protocol. Briefly, 500 ng of total RNA was used for poly(A)-mRNA selection using Oligo (dT) magnetic beads and subsequently fragmented to approximately 300bp. cDNA was synthesized using reverse transcriptase (SuperScript II, Invitrogen) and random primers. The second strand of the cDNA incorporated dUTP in place of dTTP. Double-stranded DNA was further used for library preparation. dsDNA was subjected to A-tailing and ligation of the barcoded Truseq adapters. All purification steps were performed using AMPure XP Beads. Library amplification was performed with PCR using the primer cocktail supplied in the kit. Final libraries were analyzed using Agilent DNA 1000 chip to estimate the quantity, check the size distribution, and then quantified by qPCR using the KAPA Library Quantification Kit (KapaBiosystems, Merck, Darmstadt, Germany) before amplification Illumina’s cBot. Libraries were sequenced 1 x 50bp on Illumina’s HiSeq 2500.

The quality of the fast files was checked using the FastQC software (http://www.bioinformatics.babraham.ac.uk/projects/fastqc/). An estimation of ribosomal RNA in the raw data was obtained with riboPicker [93]. Reads were aligned with the STAR mapper [94] to release M14 of the Mus musculus Gencode version of the genome (GRMm38/mm10 assembly) (https://www.gencodegenes.org/mouse/release_M14.html). A raw count of reads per gene was also obtained with STAR (-quantMode TranscriptomeSAM GeneCounts option). The R/Bioconductor package DESeq2 [95, 96] was used to assess differential expression between experimental groups (Wald statistical test + false discovery rate correction). Prior to processing the differential expression analysis, genes for which the sum of raw counts across all samples was less than two were discarded. Deregulated genes with a *p*adj <0.05 were used to disclose relevant pathway alterations in the REACTOME v77 pathway database gene expression. The analysis has been done just with the protein-coding genes. The genes difference was considered biologically relevant if they are upregulated or downregulated with a fold change of >1.2 or < 0.85, respectively. A pathway was considered relevant if it was related to neuronal and/or cerebral functions, showed significance (*p*adj < 0,05) and contained more than 10 deregulated genes.

### RT-qPCR

For RT-qPCR validations, cDNA was obtained with a High-Capacity cDNA Reverse Transcription kit (Applied Biosystems) following the supplier’s instructions. RT-qPCR reactions contained 4.5 μL cDNA, mixed with 0.5 μL 20X TaqMan Gene Expression Assays and 5 μL of 2X TaqMan Universal PCR Master Mix (Applied Biosystems) for a final volume of 10 μL. TaqMan probes used were: *Grin2b* Mm00433820_m1, *Gabrr2* Mm00433507_m1, *Kacnj6* Mm01215650_m1, *Kcna1* Mm00439977_s1, *Kcnj2* Mm00434616_m1, *Kcnq3* Mm00548884_m1 (Applied Biosystems). *Actb* Mm02619580_g1 and *Aars Mm00507627_m1* were used as endogenous controls. The assay was performed using technical duplicates per sample in 384-well optical plates with ABI Prism 7900 Sequence Detection system (Applied Biosystems, Life Technologies) following the supplier’s parameters: 50°C for 2 min, 95°C for 10 min, and 40 cycles of 95°C for 15 s and 60°C for 1 min. The Sequence Detection Software (SDS version 2.2.2, Applied Biosystems) was used for data processing. It was further analyzed with the ΔΔCt method, which consists of obtaining ΔCt by normalizing each target gene to its endogenous control, followed by subtracting the mean-ΔCt of the control group samples to each ΔCt to obtain ΔΔCt values, and finally using these ΔΔCt values as the negative exponent with base 2, thereby obtaining fold change per sample.

### Fluoro-Jade B staining

Mice were perfused seven days after the KA administration with 4 % paraformaldehyde (PFA) dissolved in 0.1 M phosphate buffer (pH = 7.3-7.4). Brains were dissected and postfixed overnight with the same fixative solution. The following day, they were cryoprotected in 0.1 M phosphate buffer containing 30 % sucrose (w/v). After freezing in dry ice, 50 µm thick coronal sections were obtained with a freezing microtome (Leica, Wetzlar, Germany). Sections containing the dorsal hippocampus were selected and rinsed in 0.1M phosphate buffer and mounted on gelatin-coated slides. The slides containing sections were dried at 37°C overnight. The following day, they were heated at 50°C for 45 min before staining to improve adhesion. The staining started with pretreatment for 3 min in absolute alcohol, followed by 1 min in 70 % ethanol and 1 min in deionized water. After that, they were oxidized in a solution of 0.06 % KMnO_4_ for 15 min. Following three rinses (2 min/each) in deionized water, they were incubated in a solution of 0.001 % Fluoro-Jade B (Chemicon, Temecula, CA, USA) containing 0.05 % of DAPI in 0.1% acetic acid for 30 min. Finally, sections were rinsed in deionized water (3 min), dehydrated with ethanol, cleared with xylene, and coverslipped with Eukitt^TM^ (Merck, Darmstadt, Germany). Considering that wild-type mice did not displayed Fluoro-Jade B labeled cells in the CA1-CA3 regions after KA treatments and only a very few background could be seen at high magnification and long exposures times (> 500 ms), the Fluoro-Jade B fluorescence in the pyramidal layer of the dorsal hippocampal region (4 sections of each mouse, n = 4 mice per genotype) was photo-documented using an Olympus BX61 epifluorescence microscope equipped with a cooled DP12L camera (Hamburg, Germany). Photomicrographs were obtained using a 40X objective with identical time exposure between preparations from each wild-type and respective knock-out mouse. No modifications were applied to the obtained pictures. Changes in Fluoro-Jade B labeling were determined by analyzing the corrected total cell fluorescence (CTCF) values (see Matamoros-Angles *et al*., [97], for details) in the pyramidal layer of hippocampal CA1-3 regions of four mice of each genotype, taking a region of interest of (200 x 100 μm) centered in the pyramidal layer. Data were expressed as mean ± S.E.M. The statistical analysis of the obtained data was performed using Mann-Whitney *U* non-parametric test in GraphPad Prism 8 software. A value of \*\*\**p* < 0.001 was considered statistically significant in the CTCF analysis.

### Primary cortical cultures of Prnp^ZH3/ ZH3^ and wild-type mice

Primary cortical cultures were fashioned from E16.5-E17.5 *Prnp^+/+^* and *Prnp^ZH3/ZH3^* mouse embryos, as explained elsewhere [98]. Brains were removed from the skull and rinsed in cold Hank’s balanced salt solution (HBSS) containing glucose (6.5 mg/ml). The meninges were removed, and the cortical lobes were isolated. Tissue pieces were treated with trypsin for 15 min at 37°C. After the addition of horse serum followed by centrifugation, cells were isolated mechanically with a polished glass pipette after treatment with 0.025 % DNAse for 10 min at 37°C. One million cells were plated on a 35 mm diameter glass-bottom gridded culture dish (Ibidi, Martinsried, Germany) previously coated with poly-D-lysine (Sigma-Aldrich). Neurobasal^TM^ medium supplemented with 6.5 mg/ml glucose, 2 mM glutamine, penicillin/streptomycin, 5 % of horse serum, and B27 was used as a culture medium (all from Invitrogen-Thermo Fisher Scientific, MA, USA). As *Prnp^0/0^*-derived cells are sensitive to serum removal [99], after 24 h, the serum was reduced to 2.5 %. The medium was changed every two days. Horse serum was entirely removed on the eighth day of culture.

### Calcium imaging in neuronal culture

Primary cortical neurons were infected 24 h after seeding with AAV9-Synapsin-GCaMP6f [60] (Watertown, MA, USA). In our cultures, the genetically encoded calcium indicators started to express 3-4 days after infection. Calcium changes in GCaMP6f-expressing neurons were recorded at 8, 11, 13, and 15 days *in vitro* (DIV) using an Olympus IX71 inverted microscope (Olympus, Hamburg, Germany), equipped with an ORCA-Flash 4.0 camera (Hamamatsu Photonics, Japan). During recording, the cells were maintained in a microscope stage incubator at 5 % CO_2_ and 37°C (Okolab S.R.L., Italy). The same region of the culture was recorded throughout the days following the culture dish grid references. Images (1024 x 1024 pixels) were captured using a 20x objective and 470 nm wavelength (CoolLED’s pE-300^white^, Delta Optics, Madrid, Spain) every 100 ms for 8-10 min using the CellSens^TM^ software (Olympus) or the Micro-Manager Open Source Microscopy Software (https://micro-manager.org). Exposure levels and frequency were maintained between cultures and evaluation days. GCaMP6f activity was measured in four different identified squares of each culture dish during these four days.

### Neuronal activity traces, spike events, and network bursts

The recordings were analyzed offline using two MATLAB^TM^ toolboxes: NETCAL (www.itsnetcal.com) [100, 101] and NeuroCa [102]. In NeuroCa, an automatic analysis was performed afterward to corroborate obtained NETCAL results. Using NETCAL, a highly contrasted image of the recording’s average fluorescence was created, and regions of interest (ROIs) were automatically detected as those objects with a circular shape whose brightness was over a preset threshold. NETCAL and NeuroCa software-rendered a similar number of ROIs and calcium traces. About 400 ROIs, uniformly covering the field of view, were typically identified per recording. The average fluorescence *Fi (t)* in each ROI *i* along the recording was then extracted, corrected from global drifts and artifacts, and finally normalized as *(Fi (t) - F_(0,i)_) / F_(0,i)_ = fi (t)*, where *F_0,i_* is the background fluorescence of the ROI. The time series of *fi (t)* was analyzed with NETCAL to infer neuronal activation timing using the Schmitt trigger method. Our analysis used +2 S.E.M. of the baseline noise as the high threshold, +1.5 S.E.M. as the low threshold, and 200 ms as the minimum event length. Calcium traces were calculated, and raster plots of network activity were then constructed by representing the trains of detected neuronal spikes over time. Next, network bursts were analyzed to quantify the ability of the neuronal networks to exhibit collective dynamics, i.e., the collective activation of a group of neurons in a short time window. Bursts were investigated using two approaches. In the first approach, raster plots of spike events were scanned to detect collective occurrences in which at least 5 % of the neurons in the network fired synchronously within a 500 ms window. This threshold of 5 % was set to disregard random activations that coincided in time. In the second approach, the fluorescence time series of all neurons in the network were averaged. The resulting trace was analyzed with the Schmitt trigger method to detect sufficiently strong fluorescence peaks associated with bursting episodes. Both approaches produced consistent results. Although the detected bursts contained a different number of participating neurons, this information was disregarded in the present analysis and treated later. The total number of detected network bursts divided by the recording duration reflected the culture’s activity and was indicated as bursts/min.

The fraction of active neurons in the network was calculated as follows. All detected ROIs were assigned as neurons. After inferring the spike trains, those neurons exhibiting at least two spikes along the recording were considered active, and their number *N_A_* was set as a proxy of the healthy population in the neuronal network. The average fraction of active neurons in each condition was then determined as *f = N_A_/N_T_*, where *N_T_* is the total number of detected ROIs.

At least ten videos of each genotype/day from different culture plates were consecutively analyzed. Data are presented as the mean of network burst/min ± S.E.M. Statistical analysis was performed using two-way ANOVA + Bonferroni’s multiple comparisons test (GraphPad Prism 8 software). Asterisks indicate significant differences between *Prnp^+/+^* and *Prnp^ZH3/ZH3^* cultures at a given DIV: \**p* < 0.05 and \*\*\**p* < 0.001. The # symbols indicate significant differences between a given DIV with the initial value at 8 DIV: ^###^*p* < 0.001.

## List of abbreviations

AMPA: Alpha-amino-3-hydroxy-5-methyl-4-isoxazolepropionic acid
ANOVA: Analysis of variance
AUC: Area under the ROC curve
CA3-CA1: Cornnu ammonis 3 and 1 of the hippocampus.
CEEA: Ethical Committee for Animal Experimentation
CTCF: Corrected total cell fluorescence
fEPSP: Field excitatory postsynaptic potential
FG: Flanking genes
GECI: Genetically encoded calcium indicator
GLUK2: Glutamate ionotropic receptor kainate type subunit 2
GPI: Glycosylphosphatidylinositol
HFS: High frequency stimulation
KA: Kainate
LTP: Long term potentiation
mGluR5: Metabotropic glutamate receptor 5
MK-801: NMDA antagonist
NMDA-R: N-methyl-D-aspartate receptor
PFA: Paraformaldehyde
*Prnp*: Cellular prion protein gene
PrP^C^: Cellular prion protein
PSD-95: Post synaptic density protein of 95 kD
PTZ: Pentylenetetrazol
ROI: Region of interest
sCJD: sporadic Creutzfeldt-Jakob disease
SEM: Standard deviation of the mean
TALEN: Transcription activator-like effector nuclease

## Acknowledgments

The authors thank Tom Yohannan for editorial advice and Miriam Segura-Feliu, María Sánchez-Enciso, and José M. González-Martín for their technical help. This research was supported by *PRPSEM* Project with ref. RTI2018-099773-B-I00 from MCINN/AEI/10.13039/501100011033/ FEDER “Una manera de hacer Europa”, the CERCA Programme, and the Commission for Universities and Research of the Department of Innovation, Universities, and Enterprise of the Generalitat de Catalunya (SGR2017-648), CIBERNED (CMED2018-2) to JADR and IF. The project leading to these results received funding from the “la Caixa” Foundation (ID 100010434) under the agreement LCF/PR/HR19/52160007 and the María de Maeztu Unit of Excellence (Institute of Neurosciences, University of Barcelona) MDM-2017-0729 to JADR. JS was supported by FIS2016-78507-C2-2-P from (MCIU/FEDER/AEI), SGR2017-1061 from the Generalitat de Catalunya, and the European Union’s Horizon 2020 research and innovation program under the grant agreement No. 713140 (MESOBRAIN). Support was also received from MINECO (BFU2017-82375-R), and Junta de Andalucía (BIO-122, UPO-1250734, and P18-FR-823) grants to AG and JMDG. F.LL. was supported by Instituto Carlos III (grant PI19-00144). A.M-A. was supported by the Tatiana Pérez de Guzmán el Bueno Foundation.

## Author contributions

A M-A, A.H., J.S., P.C., E.M., and F.L. performed most of the experiments. A M-A. and A.H. participated in all the experiments, performing and designing, and contributed equally to this project. P.C performed the behavior experiments. J.S. analyzed and interpreted the calcium imaging data. E.M. and F.L. planned and organized the RNAseq and performed the gene ontology analysis. A.G. and J.M D-G developed the electrophysiological experiments. J.A.D.R., A.G., J.M. D.-G., A.A., M.N., and I.F designed and supervised the study. J.M. D.-G. and J.A.D.R analyzed the results. J.A.D.R. and J.M. D.-G. wrote the draft and circulated it among the authors.

