## Supplementary information for "Behavioral deficits, learning impairment, and enhanced hippocampal excitability in co-isogenic *Prnp^ZH3/ZH3^* mice"

### Additional Files

**Additional File 1: Figure S1**

*Prnp*$^{ZH3/ZH3}$ mice showed similar nest-building behavior to wild-type mice. ***a***, Representative images of nests constructed by *Prnp*$^{+/+}$ (left) and *Prnp*$^{ZH3/ZH3}$ mice (right). ***b***, Quantification illustrating the mean of the nest score (see Material and Methods for details). Data are presented as mean ± S.E.M.

**Additional File 2: Figure S2**

*Prnp*$^{ZH1/ZH1}$ mice failed to acquire an instrumental learning test in the Skinner box test using a fixed ratio (1:1) schedule. ***a***, Percentage of *Prnp*$^{+/+}$ and *Prnp*$^{ZH1/ZH1}$ mice reached the selected criterion during the training sessions. ***b***, Task accuracy ((lever presses during light ON – lever presses during the light OFF) / Total number of lever presses) during the light ON / light OFF conditioning paradigm. Data are presented as a percentage in ***a*** and as mean ± S.E.M. in ***b***.

**Additional File 3: Figure S3**

*Prnp*$^{ZH3/ZH3}$ mice have deficits in acquiring instrumental learning but not motor impairment in the Rotarod test. After a training session in the Rotarod, mice were tested two consecutive days with 5 sessions in each. Latency of mice to fall (sec.) from the Rotarod in the first day (***a***) or in the second (***b***). Data are presented as mean ± S.E.M.. $*p < 0.05$ and $**p < 0.01$, two-way ANOVA + Bonferroni's multiple comparisons test.

**Additional File 4: Figure S4**

Stressed-like behavior in *Prnp*$^{ZH3/ZH3}$ mice impairing object recognition test performance. *Prnp*$^{ZH1/ZH1}$ mice also showed stressed-like behavior and failed to acquire short-term memory. ***a***, Scheme of object recognition protocol that consisted of 3 sessions (10 min each). First, mice were habituated in the empty arena. One hour later, they were placed again in the arena with two identical objects for the training session. Finally, 2-3 h later, they were once more placed in the arena, replacing one object with a novel one for the short-term memory test. Fecal bodies (*Prnp*$^{ZH3/ZH3}$) and rearing episodes (*Prnp*$^{ZH1/ZH1}$) were counted in the habituation session as an indicator of animal stress. ***b***, Number of fecal bodies generated per animal (*Prnp*$^{+/+}$ and *Prnp*$^{ZH3/ZH3}$) during the habituation session. Data are presented as mean ± S.E.M. ***c-d*** Time that *Prnp*$^{+/+}$ (***c***), and *Prnp*$^{ZH3/ZH3}$ (***d***) mice interacted with the objects in the short-term test. Data are presented as the percentage of mice interacting with the object in each time interval (0 to 35 s). ***e***, Number of rearing episodes in the habituation session (*Prnp*$^{+/+}$ and *Prnp*$^{ZH1/ZH1}$). ***f-g***, Relative time that the two exposed objects are explored to the total time invested. Object 1 ($O_1$) and object 1' ($O_1$') in the training session (***f***) and number, and object 1 ($O_1$) and object 2 ($O_2$) in the short-term memory test (***g***). Data are presented as percentages in ***C-D*** and as mean ± S.E.M. in ***b***, ***e***, ***f***, ***g***. *$p < 0.05$ and ***$p < 0.001$, Mann-Whitney *U* non-parametric test.

**Additional File 5: Table S1**

List of the protein-coding significantly downregulated genes in *Prnp*$^{ZH3/ZH3}$ hippocampus compared to *Prnp*$^{+/+}$.

**Additional File 6: Table S2**

List of the protein-coding significantly upregulated genes in *Prnp*$^{ZH3/ZH3}$ hippocampus compared to *Prnp*$^{+/+}$.

**Additional File 7: Figure S5**

Gene ontology of the genes significantly downregulated and upregulated in *Prnp*$^{ZH3/ZH3}$ hippocampus compared to *Prnp*$^{+/+}$. ***a***, Volcano plot of the protein-coding genes analyzed in the RNAseq (~16.000). In the X-axis is plotted the Fold change (log2) and in the Y-axis the *p*adj (-log10). The red line separates the significantly expressed genes (*p*adj < 0,05). The blue lines indicate the 0,85 and 1,2 fold changes. ***b***, Gene ontology analysis of the downregulated and upregulated genes with the Reactome software (see Materials and Methods for details). Validation of the main genes altered by RT-qPCR: *Grin2b* (***c***), *Gabrr2* (***d***), *Kcnj6* (***e***), *Kcna1* (***f***), *Kcnj2* (***g***), and *Kcnq3* (***h***). Data are presented as mean ± S.E.M. *$p < 0.05$ and ***$p < 0.001$.

**Additional File 8: Movie S1**

Representative movie of *Prnp*$^{+/+}$ (bottom and up-left) and *Prnp*$^{ZH3/ZH3}$ (middle and up-right) mouse behavior after KA (middle and bottom) and PBS (top) administration. Note the blinking and seizure episodes suffered by the *Prnp*$^{ZH3/ZH3}$ mice treated with KA (labeled with asterisks at the beginning of the video).

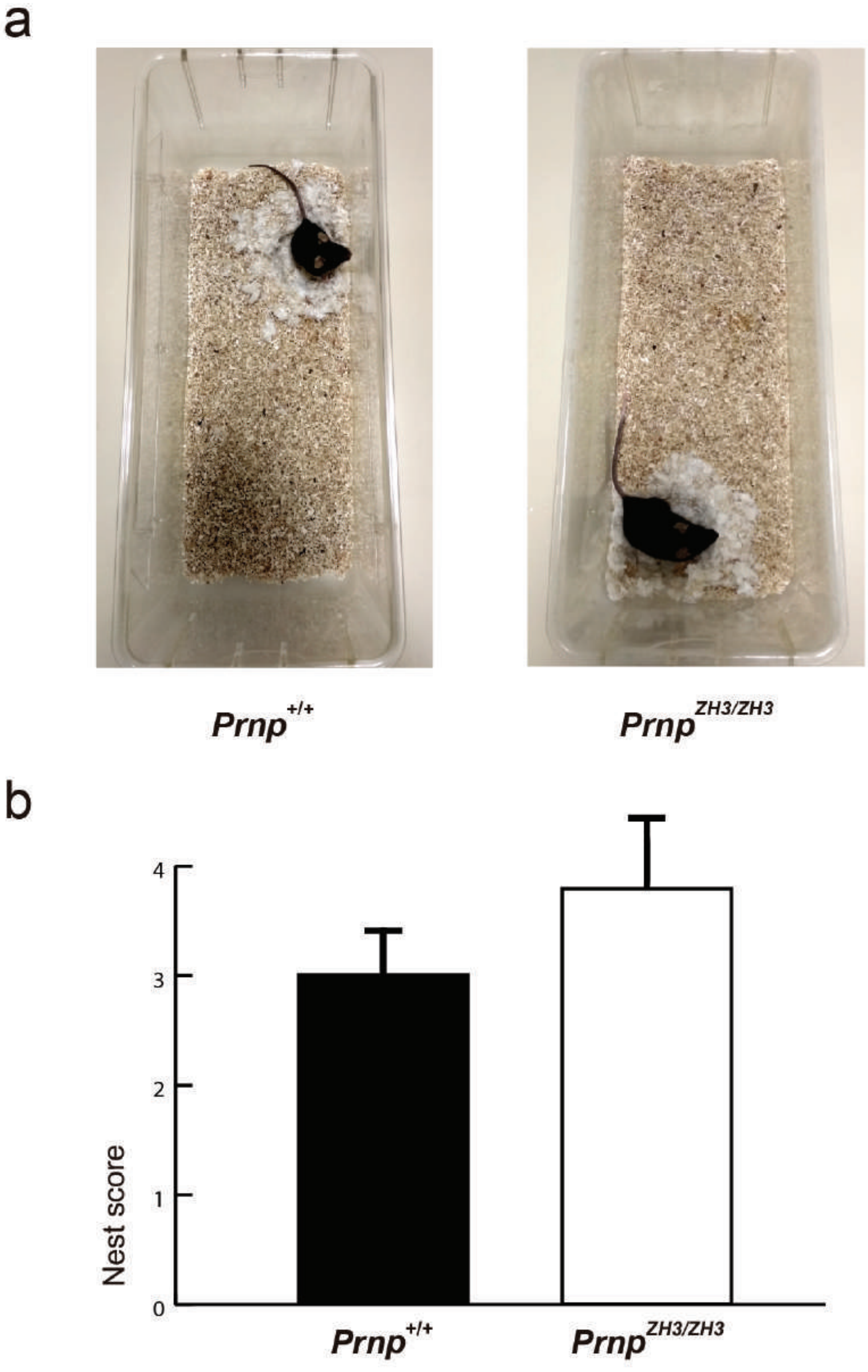


Additional File 1: Figure S1

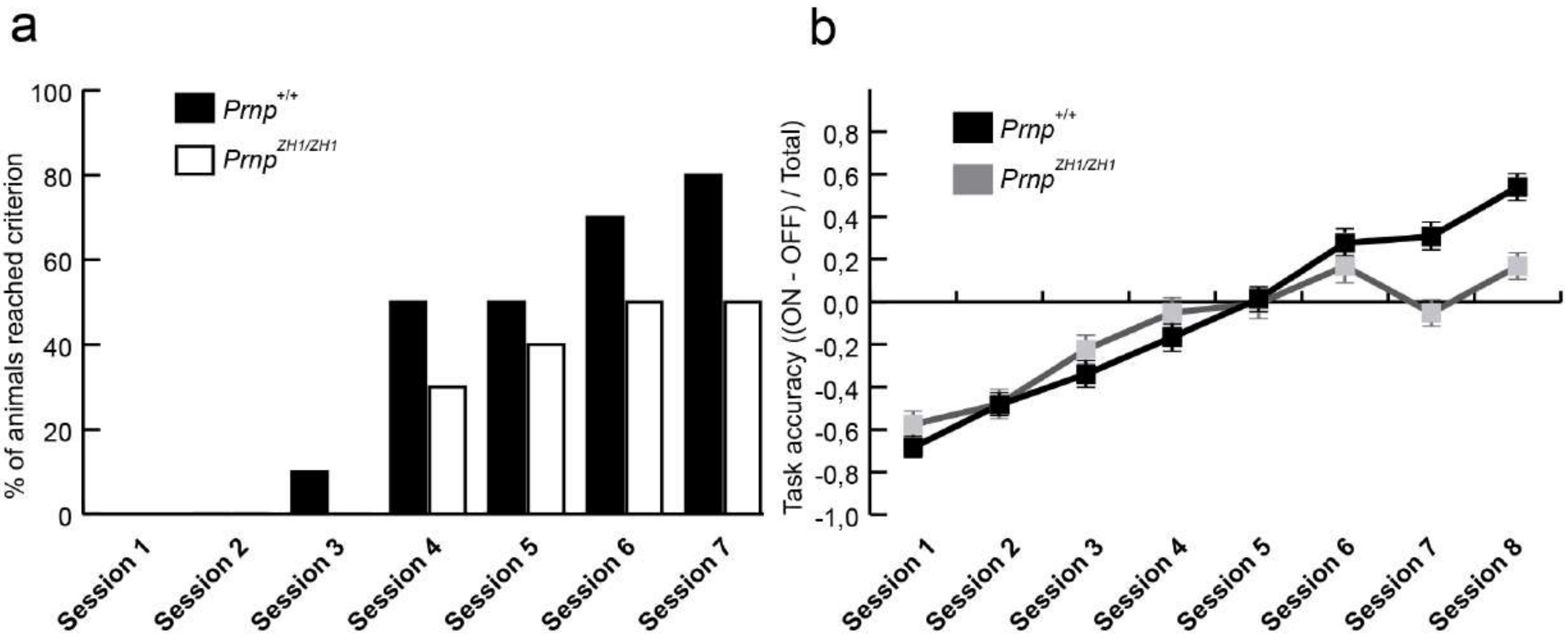


Additional File 2: Figure S2

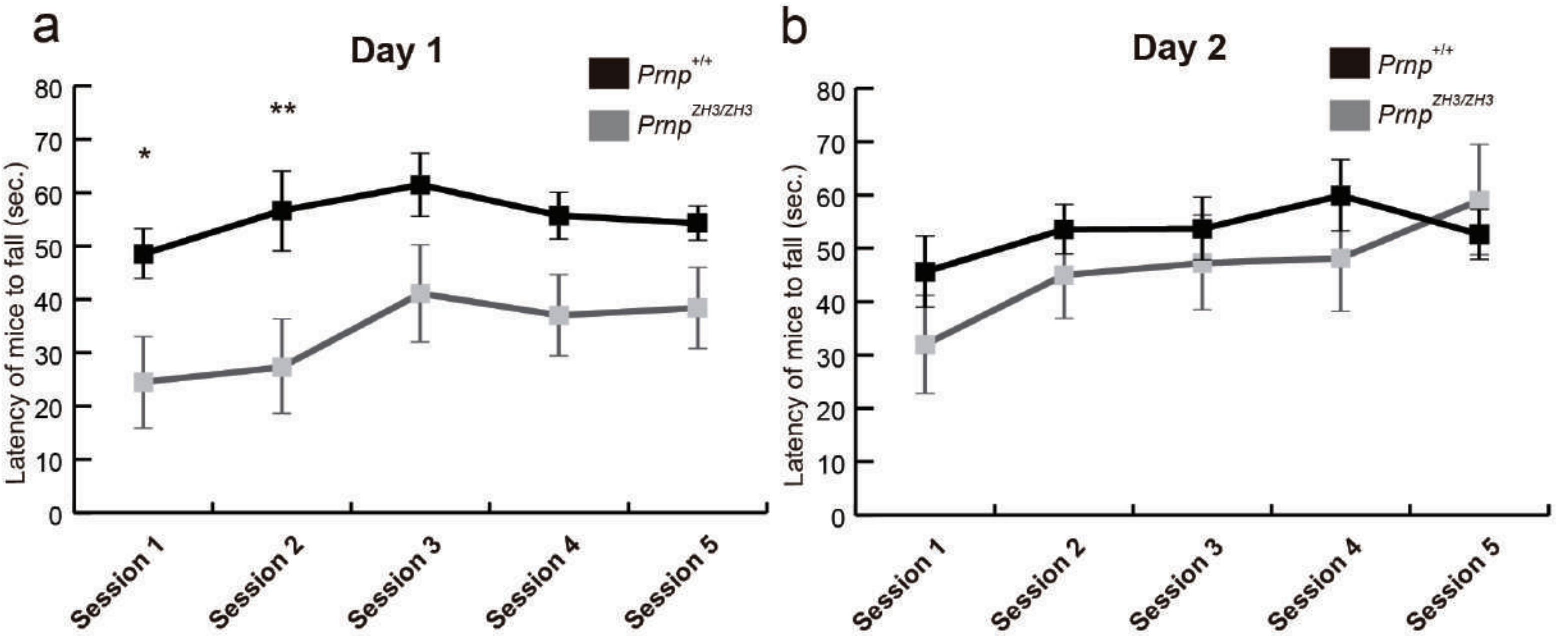


Additional File 3: Figure S3

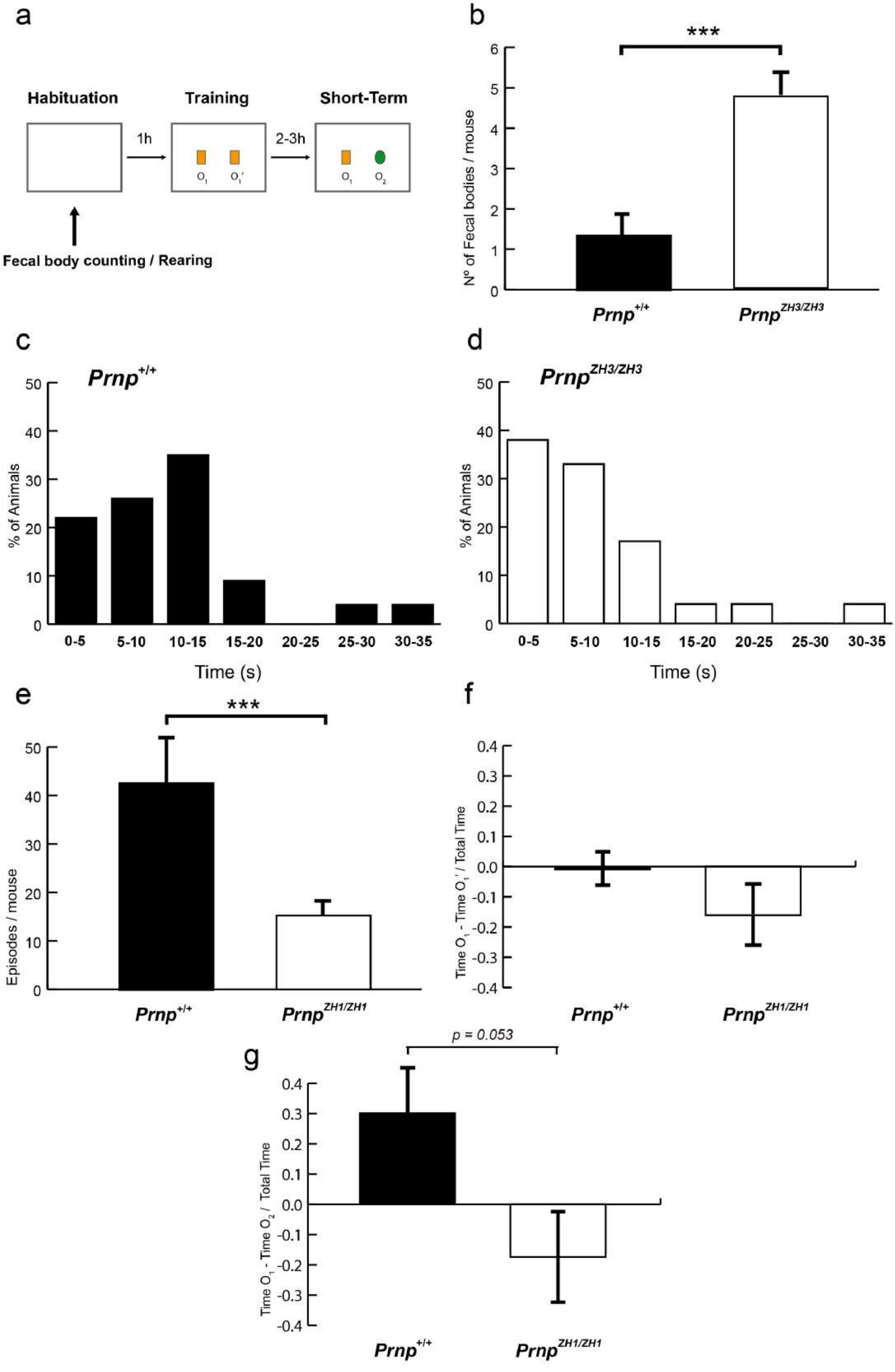


Additional File 4: Figure S4

#### Additional File 5: Table S1

### Downregulated genes in *Prnp* $^{ZH3/ZH3}$ compared to *Prnp* $^{+/+}$

| Gene | Name | padj_Ko vs Wt |
|---|---|---|
| **Abcb1b** | ATP-binding cassette, sub-family B (MDR/TAP), member 1B(Abcb1b) | 3,13E-03 |
| **Actr2** | ARP2 actin-related protein 2(Actr2) | 2,71E-04 |
| **Adcy9** | adenylate cyclase 9(Adcy9) | 4,55E-04 |
| **Adgrl3** | adhesion G protein-coupled receptor L3(Adgrl3) | 5,00E-04 |
| **Afap1l1** | actin filament associated protein 1-like 1(Afap1l1) | 1,51E-03 |
| **Aff1** | AF4/FMR2 family, member 1(Aff1) | 4,40E-02 |
| **Ago3** | argonaute RISC catalytic subunit 3(Ago3) | 3,64E-02 |
| **AI593442** | expressed sequence AI593442(AI593442) | 9,02E-03 |
| **Aim2** | absent in melanoma 2(Aim2) | 1,49E-03 |
| **Aldh1l2** | aldehyde dehydrogenase 1 family, member L2(Aldh1l2) | 1,61E-04 |
| **Aldh7a1** | aldehyde dehydrogenase family 7, member A1(Aldh7a1) | 1,02E-04 |
| **Ang** | angiogenin, ribonuclease, RNase A family, 5(Ang) | 1,07E-02 |
| **Ank3** | ankyrin 3, epithelial(Ank3) | 7,57E-03 |
| **Anln** | anillin, actin binding protein(Anln) | 4,66E-02 |
| **Ano1** | anoctamin 1, calcium activated chloride channel(Ano1) | 2,45E-02 |
| **Aoc1** | amine oxidase, copper-containing 1(Aoc1) | 1,61E-02 |
| **Ap4b1** | adaptor-related protein complex AP-4, beta 1(Ap4b1) | 2,54E-02 |
| **Arfgef3** | ARFGEF family member 3(Arfgef3) | 1,28E-02 |
| **Arhgap27** | Rho GTPase activating protein 27(Arhgap27) | 2,37E-02 |
| **Arhgef19** | Rho guanine nucleotide exchange factor (GEF) 19(Arhgef19) | 2,25E-02 |
| **Arhgef37** | Rho guanine nucleotide exchange factor (GEF) 37(Arhgef37) | 2,36E-02 |
| **Arsi** | arylsulfatase i(Arsi) | 3,15E-02 |
| **Arsj** | arylsulfatase J(Arsj) | 1,99E-06 |
| **Atf4** | activating transcription factor 4(Atf4) | 1,85E-06 |
| **Atf5** | activating transcription factor 5(Atf5) | 5,06E-04 |
| **Atm** | ataxia telangiectasia mutated(Atm) | 5,52E-03 |
| **Atp2b1** | ATPase, Ca++ transporting, plasma membrane 1(Atp2b1) | 9,92E-04 |
| **Atp5g1** | ATP synthase, H+ transporting, mitochondrial F0 complex, subunit C1 (subunit 9)(Atp5g1) | 1,08E-07 |
| **Atrx** | alpha thalassemia/mental retardation syndrome X-linked(Atrx) | 1,41E-03 |
| **Bach2** | BTB and CNC homology, basic leucine zipper transcription factor 2(Bach2) | 5,75E-03 |
| **Bbs12** | Bardet-Biedl syndrome 12 (human)(Bbs12) | 2,10E-05 |
| **Bclaf1** | BCL2-associated transcription factor 1(Bclaf1) | 3,73E-05 |
| **Bdnf** | brain derived neurotrophic factor(Bdnf) | 1,01E-07 |
| **Best3** | bestrophin 3(Best3) | 3,27E-03 |
| **Bex1** | brain expressed X-linked 1(Bex1) | 4,41E-04 |
| **Bhlhe41** | basic helix-loop-helix family, member e41(Bhlhe41) | 6,25E-04 |
| **Birc6** | baculoviral IAP repeat-containing 6(Birc6) | 2,16E-02 |
| **Birc7** | baculoviral IAP repeat-containing 7 (livin)(Birc7) | 3,64E-02 |
| **Bmpr2** | bone morphogenetic protein receptor, type II (serine/threonine kinase)(Bmpr2) | 9,38E-03 |
| **Bok** | BCL2-related ovarian killer(Bok) | 5,93E-03 |
| **Brwd3** | bromodomain and WD repeat domain containing 3(Brwd3) | 2,55E-02 |
| **Btaf1** | B-TFIID TATA-box binding protein associated factor 1(Btaf1) | 7,67E-06 |
| **C1ql2** | complement component 1, q subcomponent-like 2(C1ql2) | 4,41E-02 |
| **C2cd4a** | C2 calcium-dependent domain containing 4A(C2cd4a) | 2,62E-02 |
| **Cacna1e** | calcium channel, voltage-dependent, R type, alpha 1E subunit(Cacna1e) | 2,13E-02 |
| **Cacnb4** | calcium channel, voltage-dependent, beta 4 subunit(Cacnb4) | 1,07E-02 |
| **Cacng5** | calcium channel, voltage-dependent, gamma subunit 5(Cacng5) | 2,19E-02 |
| **Cacul1** | CDK2 associated, cullin domain 1(Cacul1) | 2,55E-03 |
| **Camp** | cathelicidin antimicrobial peptide(Camp) | 4,17E-02 |
| **Caps2** | calcyphosphine 2(Caps2) | 8,80E-03 |
| **Capza1** | capping protein (actin filament) muscle Z-line, alpha 1(Capza1) | 1,41E-03 |
| **Car9** | carbonic anhydrase 9(Car9) | 2,05E-03 |
| **Card6** | caspase recruitment domain family, member 6(Card6) | 1,29E-02 |
| **Cars** | cysteinyl-tRNA synthetase(Cars) | 3,87E-08 |
| **Cast** | calpastatin(Cast) | 4,76E-02 |
| **Cdh26** | cadherin-like 26(Cdh26) | 1,54E-02 |
| **Cdh8** | cadherin 8(Cdh8) | 3,50E-03 |
| **Cdhr4** | cadherin-related family member 4(Cdhr4) | 1,37E-02 |
| **Cfap157** | cilia and flagella associated protein 157(Cfap157) | 4,37E-02 |
| **Cgn** | cingulin(Cgn) | 4,14E-02 |
| **Chac1** | ChaC, cation transport regulator 1(Chac1) | 1,37E-06 |
| **Chaf1a** | chromatin assembly factor 1, subunit A (p150)(Chaf1a) | 4,67E-02 |
| **Chl1** | cell adhesion molecule L1-like(Chl1) | 1,80E-04 |
| **Chrna1** | cholinergic receptor, nicotinic, alpha polypeptide 1 (muscle)(Chrna1) | 4,38E-02 |
| **Chst9** | carbohydrate (N-acetylgalactosamine 4-0) sulfotransferase 9(Chst9) | 2,01E-04 |
| **Ciart** | circadian associated repressor of transcription(Ciart) | 1,13E-02 |
| **Clstn2** | calsyntenin 2(Clstn2) | 1,72E-04 |
| **Cntn3** | contactin 3(Cntn3) | 4,66E-03 |
| **Cort** | cortistatin(Cort) | 1,80E-04 |
| **Cpeb4** | cytoplasmic polyadenylation element binding protein 4(Cpeb4) | 1,96E-03 |
| **Crhbp** | corticotropin releasing hormone binding protein(Crhbp) | 9,83E-03 |
| **Csmd3** | CUB and Sushi multiple domains 3(Csmd3) | 3,82E-02 |
| **Cxcl5** | chemokine (C-X-C motif) ligand 5(Cxcl5) | 7,17E-03 |
| **D130040H23Rik** | RIKEN cDNA D130040H23 gene(D130040H23Rik) | 4,62E-02 |

#### Additional File 5: Table S1

| | | |
|---|---|---|
| **Dap3** | death associated protein 3(Dap3) | 3,10E-04 |
| **Dbf4** | DBF4 zinc finger(Dbf4) | 1,82E-02 |
| **Dbh** | dopamine beta hydroxylase(Dbh) | 4,18E-02 |
| **Defb42** | defensin beta 42(Defb42) | 1,70E-02 |
| **Dennd2c** | DENN/MADD domain containing 2C(Dennd2c) | 3,55E-02 |
| **Dgkh** | diacylglycerol kinase, eta(Dgkh) | 2,54E-04 |
| **Dgki** | diacylglycerol kinase, iota(Dgki) | 1,09E-03 |
| **Dhx33** | DEAH (Asp-Glu-Ala-His) box polypeptide 33(Dhx33) | 1,16E-04 |
| **Dnah9** | dynein, axonemal, heavy chain 9(Dnah9) | 8,65E-04 |
| **Dnajb14** | DnaJ heat shock protein family (Hsp40) member B14(Dnajb14) | 8,00E-03 |
| **Dock10** | dedicator of cytokinesis 10(Dock10) | 3,85E-02 |
| **Dock4** | dedicator of cytokinesis 4(Dock4) | 1,90E-03 |
| **Drd5** | dopamine receptor D5(Drd5) | 2,19E-02 |
| **Drosha** | drosha, ribonuclease type III(Drosha) | 5,56E-03 |
| **Dsel** | dermatan sulfate epimerase-like(Dsel) | 5,34E-04 |
| **Dthd1** | death domain containing 1(Dthd1) | 9,87E-03 |
| **Duox2** | dual oxidase 2(Duox2) | 1,50E-04 |
| **Eda2r** | ectodysplasin A2 receptor(Eda2r) | 2,42E-02 |
| **Eno1** | enolase 1, alpha non-neuron(Eno1) | 6,22E-03 |
| **Eomes** | eomesodermin(Eomes) | 2,45E-02 |
| **Epha6** | Eph receptor A6(Epha6) | 1,85E-04 |
| **Epm2aip1** | EPM2A (laforin) interacting protein 1(Epm2aip1) | 5,71E-03 |
| **Eps8l1** | EPS8-like 1(Eps8l1) | 9,80E-03 |
| **Epx** | eosinophil peroxidase(Epx) | 8,84E-03 |
| **Etaa1** | Ewing tumor-associated antigen 1(Etaa1) | 4,94E-03 |
| **Exosc9** | exosome component 9(Exosc9) | 2,80E-04 |
| **Farp1** | FERM, RhoGEF (Arhgef) and pleckstrin domain protein 1 (chondrocyte-derived)(Farp1) | 4,75E-04 |
| **Fgfbp3** | fibroblast growth factor binding protein 3(Fgfbp3) | 1,31E-04 |
| **Filip1** | filamin A interacting protein 1(Filip1) | 3,22E-03 |
| **Fkbp10** | FK506 binding protein 10(Fkbp10) | 9,60E-04 |
| **Fnip1** | folliculin interacting protein 1(Fnip1) | 2,16E-04 |
| **Foxk1** | forkhead box K1(Foxk1) | 9,17E-05 |
| **Frmpd2** | FERM and PDZ domain containing 2(Frmpd2) | 9,24E-03 |
| **Frrs1l** | ferric-chelate reductase 1 like(Frrs1l) | 4,36E-04 |
| **Fry** | FRY microtubule binding protein(Fry) | 1,39E-04 |
| **Fstl4** | follistatin-like 4(Fstl4) | 3,43E-04 |
| **Fzd3** | frizzled class receptor 3(Fzd3) | 3,44E-02 |
| **Gabrr2** | gamma-aminobutyric acid (GABA) C receptor, subunit rho 2(Gabrr2) | 4,91E-03 |
| **Garem1** | GRB2 associated regulator of MAPK1 subtype 1(Garem1) | 2,54E-06 |
| **Gbe1** | glucan (1,4-alpha-), branching enzyme 1(Gbe1) | 4,88E-02 |
| **Gdf9** | growth differentiation factor 9(Gdf9) | 6,70E-03 |
| **Gemin6** | gem (nuclear organelle) associated protein 6(Gemin6) | 8,98E-03 |
| **Gfra2** | glial cell line derived neurotrophic factor family receptor alpha 2(Gfra2) | 2,69E-02 |
| **Ggps1** | geranylgeranyl diphosphate synthase 1(Ggps1) | 1,88E-03 |
| **Gipr** | gastric inhibitory polypeptide receptor(Gipr) | 1,15E-06 |
| **Glis3** | GLIS family zinc finger 3(Glis3) | 3,45E-02 |
| **Glp2r** | glucagon-like peptide 2 receptor(Glp2r) | 2,58E-02 |
| **Gnpda1** | glucosamine-6-phosphate deaminase 1(Gnpda1) | 2,98E-05 |
| **Gpc4** | glypican 4(Gpc4) | 5,02E-03 |
| **Gpr150** | G protein-coupled receptor 150(Gpr150) | 1,62E-02 |
| **Gpr151** | G protein-coupled receptor 151(Gpr151) | 2,76E-02 |
| **Gpr21** | G protein-coupled receptor 21(Gpr21) | 1,90E-02 |
| **Gpr22** | G protein-coupled receptor 22(Gpr22) | 2,77E-02 |
| **Grin2a** | glutamate receptor, ionotropic, NMDA2A (epsilon 1)(Grin2a) | 7,81E-03 |
| **Grin2b** | glutamate receptor, ionotropic, NMDA2B (epsilon 2)(Grin2b) | 5,96E-05 |
| **Gtpbp10** | GTP-binding protein 10 (putative)(Gtpbp10) | 1,11E-04 |
| **Gucy1a2** | guanylate cyclase 1, soluble, alpha 2(Gucy1a2) | 2,12E-03 |
| **Gucy2g** | guanylate cyclase 2g(Gucy2g) | 3,87E-03 |
| **Hcn1** | hyperpolarization-activated, cyclic nucleotide-gated K+ 1(Hcn1) | 1,70E-02 |
| **Hdac1** | histone deacetylase 1(Hdac1) | 1,70E-03 |
| **Hectd2** | HECT domain containing 2(Hectd2) | 2,05E-04 |
| **Hecw2** | HECT, C2 and WW domain containing E3 ubiquitin protein ligase 2(Hecw2) | 4,59E-03 |
| **Henmt1** | HEN1 methyltransferase homolog 1 (Arabidopsis)(Henmt1) | 8,18E-03 |
| **Herc2** | HECT and RLD domain containing E3 ubiquitin protein ligase 2(Herc2) | 7,55E-03 |
| **Hfe** | hemochromatosis(Hfe) | 3,41E-02 |
| **Hfm1** | HFM1, ATP-dependent DNA helicase homolog(Hfm1) | 7,18E-05 |
| **Hjurp** | Holliday junction recognition protein(Hjurp) | 5,99E-05 |
| **Hk2** | hexokinase 2(Hk2) | 7,85E-03 |
| **Hmgn2** | high mobility group nucleosomal binding domain 2(Hmgn2) | 2,75E-03 |
| **Hpdl** | 4-hydroxyphenylpyruvate dioxygenase-like(Hpdl) | 3,81E-02 |
| **Hspb6** | heat shock protein, alpha-crystallin-related, B6(Hspb6) | 5,71E-03 |
| **Htr5b** | 5-hydroxytryptamine (serotonin) receptor 5B(Htr5b) | 1,43E-02 |
| **Hunk** | hormonally upregulated Neu-associated kinase(Hunk) | 1,17E-02 |
| **Iars2** | isoleucine-tRNA synthetase 2, mitochondrial(Iars2) | 7,82E-06 |
| **Ide** | insulin degrading enzyme(Ide) | 3,91E-09 |
| **Idua** | iduronidase, alpha-L-(Idua) | 2,12E-03 |
| **Igfbpl1** | insulin-like growth factor binding protein-like 1(Igfbpl1) | 4,39E-02 |
| **Igtp** | interferon gamma induced GTPase(Igtp) | 3,55E-02 |
| **Il1rap** | interleukin 1 receptor accessory protein(Il1rap) | 1,15E-02 |

#### Additional File 5: Table S1

| | | |
|---|---|---|
| **Ildr2** | immunoglobulin-like domain containing receptor 2(Ildr2) | 8,67E-03 |
| **Inhba** | inhibin beta-A(Inhba) | 4,55E-02 |
| **Ints7** | integrator complex subunit 7(Ints7) | 1,81E-09 |
| **Irgm2** | immunity-related GTPase family M member 2(Irgm2) | 3,15E-02 |
| **Itpka** | inositol 1,4,5-trisphosphate 3-kinase A(Itpka) | 3,91E-02 |
| **Izumo4** | IZUMO family member 4(Izumo4) | 4,17E-02 |
| **Kansl1l** | KAT8 regulatory NSL complex subunit 1-like(Kansl1l) | 3,30E-03 |
| **Kcna1** | potassium voltage-gated channel, shaker-related subfamily, member 1(Kcna1) | 7,51E-07 |
| **Kcna4** | potassium voltage-gated channel, shaker-related subfamily, member 4(Kcna4) | 4,41E-02 |
| **Kcnb2** | potassium voltage gated channel, Shab-related subfamily, member 2(Kcnb2) | 7,68E-04 |
| **Kcnh5** | potassium voltage-gated channel, subfamily H (eag-related), member 5(Kcnh5) | 1,02E-02 |
| **Kcnh7** | potassium voltage-gated channel, subfamily H (eag-related), member 7(Kcnh7) | 1,44E-02 |
| **Kcnj2** | potassium inwardly-rectifying channel, subfamily J, member 2(Kcnj2) | 3,05E-03 |
| **Kcnj6** | potassium inwardly-rectifying channel, subfamily J, member 6(Kcnj6) | 7,00E-05 |
| **Kcnq3** | potassium voltage-gated channel, subfamily Q, member 3(Kcnq3) | 5,24E-03 |
| **Kcns1** | K+ voltage-gated channel, subfamily S, 1(Kcns1) | 2,25E-02 |
| **Kctd16** | potassium channel tetramerisation domain containing 16(Kctd16) | 3,49E-03 |
| **Kctd4** | potassium channel tetramerisation domain containing 4(Kctd4) | 2,37E-02 |
| **Khdrbs1** | KH domain containing, RNA binding, signal transduction associated 1(Khdrbs1) | 6,55E-06 |
| **Klhl40** | kelch-like 40(Klhl40) | 4,26E-02 |
| **Krt12** | keratin 12(Krt12) | 4,76E-05 |
| **Krt222** | keratin 222(Krt222) | 6,70E-04 |
| **Krt9** | keratin 9(Krt9) | 1,76E-08 |
| **Lactb2** | lactamase, beta 2(Lactb2) | 6,41E-03 |
| **Lamc2** | laminin, gamma 2(Lamc2) | 3,94E-02 |
| **Lats2** | large tumor suppressor 2(Lats2) | 3,36E-02 |
| **Lca5** | Leber congenital amaurosis 5 (human)(Lca5) | 1,97E-03 |
| **Lcp1** | lymphocyte cytosolic protein 1(Lcp1) | 8,36E-04 |
| **Lefty1** | left right determination factor 1(Lefty1) | 5,71E-10 |
| **Lefty2** | left-right determination factor 2(Lefty2) | 1,02E-02 |
| **Lmbrd2** | LMBR1 domain containing 2(Lmbrd2) | 3,85E-03 |
| **Lnpep** | leucyl/cystinyl aminopeptidase(Lnpep) | 1,63E-02 |
| **Lpar6** | lysophosphatidic acid receptor 6(Lpar6) | 2,76E-03 |
| **Lrp2** | low density lipoprotein receptor-related protein 2(Lrp2) | 1,28E-02 |
| **Lrrc46** | leucine rich repeat containing 46(Lrrc46) | 4,29E-02 |
| **Lrrtm4** | leucine rich repeat transmembrane neuronal 4(Lrrtm4) | 3,09E-04 |
| **Ltf** | lactotransferrin(Ltf) | 3,72E-02 |
| **Lyrm7** | LYR motif containing 7(Lyrm7) | 5,28E-12 |
| **Lysmd1** | LysM, putative peptidoglycan-binding, domain containing 1(Lysmd1) | 1,84E-02 |
| **Lyst** | lysosomal trafficking regulator(Lyst) | 3,17E-02 |
| **Man1a** | mannosidase 1, alpha(Man1a) | 9,92E-05 |
| **Manba** | mannosidase, beta A, lysosomal(Manba) | 4,74E-08 |
| **Map1b** | microtubule-associated protein 1B(Map1b) | 1,03E-02 |
| **Mars2** | methionine-tRNA synthetase 2 (mitochondrial)(Mars2) | 1,24E-03 |
| **Mast4** | microtubule associated serine/threonine kinase family member 4(Mast4) | 2,89E-02 |
| **Mcm2** | minichromosome maintenance complex component 2(Mcm2) | 1,98E-02 |
| **Mcm3** | minichromosome maintenance complex component 3(Mcm3) | 3,84E-02 |
| **Mcm5** | minichromosome maintenance complex component 5(Mcm5) | 9,38E-03 |
| **Med13** | mediator complex subunit 13(Med13) | 3,45E-02 |
| **Megf6** | multiple EGF-like-domains 6(Megf6) | 2,58E-02 |
| **Met** | met proto-oncogene(Met) | 3,48E-02 |
| **Mex3b** | mex3 RNA binding family member B(Mex3b) | 3,55E-02 |
| **Mfn2** | mitofusin 2(Mfn2) | 1,57E-06 |
| **Micu2** | mitochondrial calcium uptake 2(Micu2) | 7,18E-04 |
| **Mob1b** | MOB kinase activator 1B(Mob1b) | 2,45E-02 |
| **Mplkip** | M-phase specific PLK1 intereacting protein(Mplkip) | 4,20E-02 |
| **Mthfd2** | methylenetetrahydrofolate dehydrogenase (NAD+ dependent), methenyltetrahydrofolate cyclohydrolase(Mthfd2) | 6,61E-06 |
| **Mtus2** | microtubule associated tumor suppressor candidate 2(Mtus2) | 1,49E-02 |
| **Musk** | muscle, skeletal, receptor tyrosine kinase(Musk) | 3,02E-06 |
| **Myb** | myeloblastosis oncogene(Myb) | 3,99E-02 |
| **Mylk3** | myosin light chain kinase 3(Mylk3) | 6,22E-03 |
| **Myt1l** | myelin transcription factor 1-like(Myt1l) | 7,64E-05 |
| **Mzt1** | mitotic spindle organizing protein 1(Mzt1) | 1,30E-03 |
| **Nars** | asparaginyl-tRNA synthetase(Nars) | 1,78E-08 |
| **Ncapd2** | non-SMC condensin I complex, subunit D2(Ncapd2) | 1,73E-02 |
| **Ncf2** | neutrophil cytosolic factor 2(Ncf2) | 7,67E-04 |
| **Ndst4** | N-deacetylase/N-sulfotransferase (heparin glucosaminyl) 4(Ndst4) | 4,37E-03 |
| **Nek10** | NIMA (never in mitosis gene a)- related kinase 10(Nek10) | 1,44E-02 |
| **Nf1** | neurofibromatosis 1(Nf1) | 1,71E-04 |
| **Nfe2l3** | nuclear factor, erythroid derived 2, like 3(Nfe2l3) | 2,25E-02 |
| **Ngp** | neutrophilic granule protein(Ngp) | 4,41E-02 |
| **Nhsl2** | NHS-like 2(Nhsl2) | 3,28E-02 |
| **Nkain3** | Na+/K+ transporting ATPase interacting 3(Nkain3) | 3,77E-02 |
| **Nodal** | nodal(Nodal) | 3,03E-02 |
| **Nos1ap** | nitric oxide synthase 1 (neuronal) adaptor protein(Nos1ap) | 1,20E-05 |
| **Notch1** | notch 1(Notch1) | 1,90E-03 |
| **Npy2r** | neuropeptide Y receptor Y2(Npy2r) | 1,35E-05 |
| **Nr1d1** | nuclear receptor subfamily 1, group D, member 1(Nr1d1) | 3,69E-02 |
| **Nr1h4** | nuclear receptor subfamily 1, group H, member 4(Nr1h4) | 1,21E-02 |

#### Additional File 5: Table S1

| | | |
|---|---|---|
| **Nr3c2** | nuclear receptor subfamily 3, group C, member 2(Nr3c2) | 3,68E-02 |
| **Nrip1** | nuclear receptor interacting protein 1(Nrip1) | 3,50E-03 |
| **Nrxn1** | neurexin I(Nrxn1) | 9,65E-03 |
| **Nt5c1a** | 5'-nucleotidase, cytosolic IA(Nt5c1a) | 3,10E-04 |
| **Ntrk3** | neurotrophic tyrosine kinase, receptor, type 3(Ntrk3) | 7,33E-03 |
| **Nudcd1** | NudC domain containing 1(Nudcd1) | 2,64E-02 |
| **Nudt15** | nudix (nucleoside diphosphate linked moiety X)-type motif 15(Nudt15) | 1,41E-02 |
| **Nudt6** | nudix (nucleoside diphosphate linked moiety X)-type motif 6(Nudt6) | 2,96E-02 |
| **Nup93** | nucleoporin 93(Nup93) | 1,87E-03 |
| **Nvl** | nuclear VCP-like(Nvl) | 1,78E-07 |
| **Nyap2** | neuronal tyrosine-phophorylated phosphoinositide 3-kinase adaptor 2(Nyap2) | 3,32E-03 |
| **Ogg1** | 8-oxoguanine DNA-glycosylase 1(Ogg1) | 1,25E-02 |
| **Olfm4** | olfactomedin 4(Olfm4) | 2,13E-03 |
| **Olfml3** | olfactomedin-like 3(Olfml3) | 2,08E-04 |
| **Olfr316** | olfactory receptor 316(Olfr316) | 1,63E-05 |
| **Olfr317** | olfactory receptor 317(Olfr317) | 1,47E-02 |
| **Osgep** | O-sialoglycoprotein endopeptidase(Osgep) | 3,50E-03 |
| **Palmd** | palmdelphin(Palmd) | 2,54E-04 |
| **Paqr8** | progestin and adipoQ receptor family member VIII(Paqr8) | 1,41E-05 |
| **Parp1** | poly (ADP-ribose) polymerase family, member 1(Parp1) | 1,67E-04 |
| **Pcdh17** | protocadherin 17(Pcdh17) | 1,69E-08 |
| **Pcdh9** | protocadherin 9(Pcdh9) | 1,87E-04 |
| **Pcdhb12** | protocadherin beta 12(Pcdhb12) | 6,27E-06 |
| **Pcdhb15** | protocadherin beta 15(Pcdhb15) | 2,18E-05 |
| **Pcdhb16** | protocadherin beta 16(Pcdhb16) | 4,60E-04 |
| **Pcdhb17** | protocadherin beta 17(Pcdhb17) | 7,96E-04 |
| **Pcdhb18** | protocadherin beta 18(Pcdhb18) | 2,87E-03 |
| **Pcdhb19** | protocadherin beta 19(Pcdhb19) | 4,22E-03 |
| **Pcdhb9** | protocadherin beta 9(Pcdhb9) | 4,74E-08 |
| **Pcdhga10** | protocadherin gamma subfamily A, 10(Pcdhga10) | 1,06E-03 |
| **Pcdhga11** | protocadherin gamma subfamily A, 11(Pcdhga11) | 7,85E-07 |
| **Pcdhga3** | protocadherin gamma subfamily A, 3(Pcdhga3) | 3,18E-03 |
| **Pcdhga8** | protocadherin gamma subfamily A, 8(Pcdhga8) | 9,67E-04 |
| **Pcdhga9** | protocadherin gamma subfamily A, 9(Pcdhga9) | 9,89E-07 |
| **Pcdhgb1** | protocadherin gamma subfamily B, 1(Pcdhgb1) | 2,27E-02 |
| **Pctp** | phosphatidylcholine transfer protein(Pctp) | 1,09E-03 |
| **Pde6h** | phosphodiesterase 6H, cGMP-specific, cone, gamma(Pde6h) | 4,80E-04 |
| **Pdf** | peptide deformylase (mitochondrial)(Pdf) | 3,95E-02 |
| **Pdp2** | pyruvate dehyrogenase phosphatase catalytic subunit 2(Pdp2) | 9,48E-03 |
| **Pex5l** | peroxisomal biogenesis factor 5-like(Pex5l) | 7,00E-04 |
| **Pi15** | peptidase inhibitor 15(Pi15) | 1,14E-02 |
| **Pigw** | phosphatidylinositol glycan anchor biosynthesis, class W(Pigw) | 4,89E-02 |
| **Pkd2** | polycystic kidney disease 2(Pkd2) | 9,37E-05 |
| **Pkd2l2** | polycystic kidney disease 2-like 2(Pkd2l2) | 4,27E-03 |
| **Pkdrej** | polycystin (PKD) family receptor for egg jelly(Pkdrej) | 1,49E-02 |
| **Pm20d2** | peptidase M20 domain containing 2(Pm20d2) | 1,97E-03 |
| **Pop1** | processing of precursor 1, ribonuclease P/MRP family, (S. cerevisiae)(Pop1) | 7,94E-04 |
| **Ppm1e** | protein phosphatase 1E (PP2C domain containing)(Ppm1e) | 1,63E-03 |
| **Ppp1r3e** | protein phosphatase 1, regulatory (inhibitor) subunit 3E(Ppp1r3e) | 8,07E-08 |
| **Prdm16** | PR domain containing 16(Prdm16) | 2,04E-03 |
| **Prickle2** | prickle planar cell polarity protein 2(Prickle2) | 4,01E-04 |
| **Prnp** | prion protein(Prnp) | 8,64E-07 |
| **Prox1** | prospero homeobox 1(Prox1) | 5,80E-03 |
| **Psmb5** | proteasome (prosome, macropain) subunit, beta type 5(Psmb5) | 4,98E-04 |
| **Ptchd1** | patched domain containing 1(Ptchd1) | 3,27E-02 |
| **Ptchd4** | patched domain containing 4(Ptchd4) | 1,58E-04 |
| **Ptgs2** | prostaglandin-endoperoxide synthase 2(Ptgs2) | 2,91E-09 |
| **Ptprj** | protein tyrosine phosphatase, receptor type, J(Ptprj) | 4,33E-05 |
| **Pttg1** | pituitary tumor-transforming gene 1(Pttg1) | 1,45E-02 |
| **Pura** | purine rich element binding protein A(Pura) | 8,60E-03 |
| **Rad1** | RAD1 checkpoint DNA exonuclease(Rad1) | 9,77E-03 |
| **Rad51ap2** | RAD51 associated protein 2(Rad51ap2) | 1,07E-02 |
| **Rasal2** | RAS protein activator like 2(Rasal2) | 1,76E-06 |
| **Rbm8a** | RNA binding motif protein 8a(Rbm8a) | 1,95E-02 |
| **Rc3h1** | RING CCCH (C3H) domains 1(Rc3h1) | 2,45E-02 |
| **Rc3h2** | ring finger and CCCH-type zinc finger domains 2(Rc3h2) | 1,08E-02 |
| **Retnlg** | resistin like gamma(Retnlg) | 2,84E-02 |
| **Rev3l** | REV3 like, DNA directed polymerase zeta catalytic subunit(Rev3l) | 6,26E-05 |
| **Riiad1** | regulatory subunit of type II PKA R-subunit (RIIa) domain containing 1(Riiad1) | 2,05E-03 |
| **Rimbp3** | RIMS binding protein 3(Rimbp3) | 3,89E-03 |
| **Ror1** | receptor tyrosine kinase-like orphan receptor 1(Ror1) | 1,72E-03 |
| **Rorc** | RAR-related orphan receptor gamma(Rorc) | 1,36E-05 |
| **Rpl26** | ribosomal protein L26(Rpl26) | 7,63E-50 |
| **Rpl34** | ribosomal protein L34(Rpl34) | 6,91E-71 |
| **Rpl5** | ribosomal protein L5(Rpl5) | 4,18E-08 |
| **Rps19** | ribosomal protein S19(Rps19) | 3,20E-02 |
| **Rsg1** | REM2 and RAB-like small GTPase 1(Rsg1) | 2,41E-02 |
| **Rsl1** | regulator of sex limited protein 1(Rsl1) | 7,44E-03 |
| **S1pr5** | sphingosine-1-phosphate receptor 5(S1pr5) | 4,54E-02 |

#### Additional File 5: Table S1

| | | |
|---|---|---|
| **Samd3** | sterile alpha motif domain containing 3(Samd3) | 1,06E-02 |
| **Samd5** | sterile alpha motif domain containing 5(Samd5) | 4,90E-02 |
| **Scai** | suppressor of cancer cell invasion(Scai) | 4,90E-02 |
| **Scn3a** | sodium channel, voltage-gated, type III, alpha(Scn3a) | 2,32E-05 |
| **Scnm1** | sodium channel modifier 1(Scnm1) | 3,18E-06 |
| **Scoc** | short coiled-coil protein(Scoc) | 4,06E-02 |
| **Sec24d** | Sec24 related gene family, member D (S. cerevisiae)(Sec24d) | 5,47E-13 |
| **Sema3e** | sema domain, immunoglobulin domain (Ig), short basic domain, secreted, (semaphorin) 3E(Sema3e) | 4,90E-03 |
| **Sema5a** | sema domain, seven thrombospondin repeats (type 1 and type 1-like), semaphorin 5A | 2,84E-02 |
| **Shroom4** | shroom family member 4(Shroom4) | 9,40E-03 |
| **Siglece** | sialic acid binding Ig-like lectin E(Siglece) | 5,18E-03 |
| **Sil1** | endoplasmic reticulum chaperone SIL1 homolog (S. cerevisiae)(Sil1) | 7,81E-03 |
| **Skil** | SKI-like(Skil) | 2,79E-04 |
| **Slc1a5** | solute carrier family 1 (neutral amino acid transporter), member 5(Slc1a5) | 3,63E-02 |
| **Slc22a15** | solute carrier family 22 (organic anion/cation transporter), member 15(Slc22a15) | 1,94E-04 |
| **Slc2a13** | solute carrier family 2 (facilitated glucose transporter), member 13(Slc2a13) | 3,46E-03 |
| **Slc35a5** | solute carrier family 35, member A5(Slc35a5) | 1,85E-06 |
| **Slc6a19** | solute carrier family 6 (neurotransmitter transporter), member 19(Slc6a19) | 3,64E-02 |
| **Slc7a1** | solute carrier family 7 (cationic amino acid transporter, y+ system), member 1(Slc7a1) | 5,99E-05 |
| **Slc7a5** | solute carrier family 7 (cationic amino acid transporter, y+ system), member 5(Slc7a5) | 1,36E-05 |
| **Slc8a1** | solute carrier family 8 (sodium/calcium exchanger), member 1(Slc8a1) | 1,01E-06 |
| **Slitrk4** | SLIT and NTRK-like family, member 4(Slitrk4) | 2,62E-03 |
| **Snrpd1** | small nuclear ribonucleoprotein D1(Snrpd1) | 1,67E-06 |
| **Sntn** | sentan, cilia apical structure protein(Sntn) | 4,04E-02 |
| **Sorcs3** | sortilin-related VPS10 domain containing receptor 3(Sorcs3) | 3,05E-03 |
| **Sorl1** | sortilin-related receptor, LDLR class A repeats-containing(Sorl1) | 6,13E-04 |
| **Sox9** | SRY (sex determining region Y)-box 9(Sox9) | 3,94E-03 |
| **Sp8** | trans-acting transcription factor 8(Sp8) | 4,40E-02 |
| **Srl** | sarcalumenin(Srl) | 9,74E-03 |
| **Srp9** | signal recognition particle 9(Srp9) | 1,05E-19 |
| **St8sia1** | ST8 alpha-N-acetyl-neuraminide alpha-2,8-sialyltransferase 1(St8sia1) | 3,77E-04 |
| **Stab1** | stabilin 1(Stab1) | 3,01E-03 |
| **Stac** | src homology three (SH3) and cysteine rich domain(Stac) | 2,30E-02 |
| **Stat5a** | signal transducer and activator of transcription 5A(Stat5a) | 5,71E-03 |
| **Stk26** | serine/threonine kinase 26(Stk26) | 4,37E-02 |
| **Stpg1** | sperm tail PG rich repeat containing 1(Stpg1) | 1,06E-02 |
| **Stxbp5l** | syntaxin binding protein 5-like(Stxbp5l) | 3,17E-04 |
| **Sult1c1** | sulfotransferase family, cytosolic, 1C, member 1(Sult1c1) | 5,34E-03 |
| **Syn3** | synapsin III(Syn3) | 5,59E-06 |
| **Syncrip** | synaptotagmin binding, cytoplasmic RNA interacting protein(Syncrip) | 4,41E-02 |
| **Syne1** | spectrin repeat containing, nuclear envelope 1(Syne1) | 2,05E-02 |
| **Syne2** | spectrin repeat containing, nuclear envelope 2(Syne2) | 1,34E-02 |
| **Tanc1** | tetratricopeptide repeat, ankyrin repeat and coiled-coil containing 1(Tanc1) | 3,12E-03 |
| **Tbc1d8b** | TBC1 domain family, member 8B(Tbc1d8b) | 3,63E-02 |
| **Tchh** | trichohyalin(Tchh) | 4,34E-02 |
| **Tdg** | thymine DNA glycosylase(Tdg) | 1,15E-04 |
| **Tecta** | tectorin alpha(Tecta) | 2,23E-02 |
| **Tenm1** | teneurin transmembrane protein 1(Tenm1) | 1,93E-02 |
| **Tenm3** | teneurin transmembrane protein 3(Tenm3) | 1,63E-02 |
| **Them4** | thioesterase superfamily member 4(Them4) | 2,02E-03 |
| **Thoc2** | THO complex 2(Thoc2) | 4,85E-03 |
| **Thrsp** | thyroid hormone responsive(Thrsp) | 1,08E-02 |
| **Tll1** | tolloid-like(Tll1) | 2,37E-03 |
| **Tmem182** | transmembrane protein 182(Tmem182) | 2,34E-02 |
| **Tmem245** | transmembrane protein 245(Tmem245) | 3,63E-02 |
| **Tmem260** | transmembrane protein 260(Tmem260) | 5,06E-04 |
| **Trib3** | tribbles pseudokinase 3(Trib3) | 2,15E-02 |
| **Trim17** | tripartite motif-containing 17(Trim17) | 1,07E-03 |
| **Trim2** | tripartite motif-containing 2(Trim2) | 2,79E-05 |
| **Trpc4** | transient receptor potential cation channel, subfamily C, member 4(Trpc4) | 1,71E-02 |
| **Trpc5** | transient receptor potential cation channel, subfamily C, member 5(Trpc5) | 6,11E-04 |
| **Trrap** | transformation/transcription domain-associated protein(Trrap) | 8,34E-03 |
| **Ttc30b** | tetratricopeptide repeat domain 30B(Ttc30b) | 1,18E-02 |
| **Ttf2** | transcription termination factor, RNA polymerase II(Ttf2) | 9,94E-12 |
| **Ttn** | titin(Ttn) | 7,74E-03 |
| **Uba1y** | ubiquitin-activating enzyme, Chr Y(Uba1y) | 2,21E-02 |
| **Ubxn10** | UBX domain protein 10(Ubxn10) | 2,84E-02 |
| **Uhrf1bp1l** | UHRF1 (ICBP90) binding protein 1-like(Uhrf1bp1l) | 3,80E-05 |
| **Usp1** | ubiquitin specific peptidase 1(Usp1) | 1,72E-04 |
| **Usp2** | ubiquitin specific peptidase 2(Usp2) | 5,70E-03 |
| **Usp29** | ubiquitin specific peptidase 29(Usp29) | 1,53E-02 |
| **Usp34** | ubiquitin specific peptidase 34(Usp34) | 7,00E-06 |
| **Usp53** | ubiquitin specific peptidase 53(Usp53) | 1,82E-04 |
| **Vps13c** | vacuolar protein sorting 13C(Vps13c) | 1,52E-02 |
| **Vps50** | VPS50 EARP/GARPII complex subunit(Vps50) | 2,06E-06 |
| **Vwa3b** | von Willebrand factor A domain containing 3B(Vwa3b) | 7,10E-03 |
| **Wdhd1** | WD repeat and HMG-box DNA binding protein 1(Wdhd1) | 2,51E-07 |
| **Wipf3** | WAS/WASL interacting protein family, member 3(Wipf3) | 3,59E-02 |
| **Xrn1** | 5'-3' exoribonuclease 1(Xrn1) | 6,22E-05 |

#### Additional File 6: Table S2

### Upregulated genes in *Prnp* $^{ZH3/ZH3}$ compared to *Prnp* $^{+/+}$

| Gene | Name | padj_Ko vs Wt |
|---|---|---|
| **Abhd8** | abhydrolase domain containing 8(Abhd8) | 0,000108971 |
| **Acad12** | acyl-Coenzyme A dehydrogenase family, member 12(Acad12) | 0,00370349 |
| **Adamts2** | a disintegrin-like and metallopeptidase (reprolysin type) with thrombospondin type 1 motif, 2(Adamts2) | 0,013525708 |
| **Adamtsl5** | ADAMTS-like 5(Adamtsl5) | 2,82101E-05 |
| **Adarb2** | adenosine deaminase, RNA-specific, B2(Adarb2) | 0,007849456 |
| **Adhfe1** | alcohol dehydrogenase, iron containing, 1(Adhfe1) | 0,002190853 |
| **Adora2b** | adenosine A2b receptor(Adora2b) | 4,24189E-18 |
| **Adrb2** | adrenergic receptor, beta 2(Adrb2) | 0,000439986 |
| **AI987944** | expressed sequence AI987944(AI987944) | 1,92487E-14 |
| **Ajuba** | ajuba LIM protein(Ajuba) | 0,005071766 |
| **Aldh1a3** | aldehyde dehydrogenase family 1, subfamily A3(Aldh1a3) | 0,000443768 |
| **Aldh1l1** | aldehyde dehydrogenase 1 family, member L1(Aldh1l1) | 0,001718877 |
| **Anapc13** | anaphase promoting complex subunit 13(Anapc13) | 1,99277E-06 |
| **Angptl7** | angiopoietin-like 7(Angptl7) | 0,024025842 |
| **Apoc3** | apolipoprotein C-III(Apoc3) | 0,019261329 |
| **Arhgap26** | Rho GTPase activating protein 26(Arhgap26) | 1,09315E-08 |
| **Arhgap33** | Rho GTPase activating protein 33(Arhgap33) | 0,000123622 |
| **Arhgdig** | Rho GDP dissociation inhibitor (GDI) gamma(Arhgdig) | 3,15408E-07 |
| **Arl5c** | ADP-ribosylation factor-like 5C(Arl5c) | 0,026903754 |
| **Aspdh** | aspartate dehydrogenase domain containing(Aspdh) | 0,017047962 |
| **Aspg** | asparaginase(Aspg) | 0,006771728 |
| **Atp5d** | ATP synthase, H+ transporting, mitochondrial F1 complex, delta subunit(Atp5d) | 0,00039469 |
| **Atp5e** | ATP synthase, H+ transporting, mitochondrial F1 complex, epsilon subunit(Atp5e) | 2,71606E-05 |
| **Atp5j2** | ATP synthase, H+ transporting, mitochondrial F0 complex, subunit F2(Atp5j2) | 0,000951904 |
| **B3gnt7** | UDP-GlcNAc:betaGal beta-1,3-N-acetylglucosaminyltransferase 7(B3gnt7) | 0,026262529 |
| **B9d1** | B9 protein domain 1(B9d1) | 0,00013909 |
| **Bace2** | beta-site APP-cleaving enzyme 2(Bace2) | 0,04258515 |
| **Baiap2l1** | BAI1-associated protein 2-like 1(Baiap2l1) | 3,46451E-05 |
| **Blnk** | B cell linker(Blnk) | 0,01909235 |
| **Blvrb** | biliverdin reductase B (flavin reductase (NADPH))(Blvrb) | 0,003374943 |
| **Bola2** | bolA-like 2 (E. coli)(Bola2) | 0,001621668 |
| **C1qa** | complement component 1, q subcomponent, alpha polypeptide(C1qa) | 0,001061274 |
| **C2cd4c** | C2 calcium-dependent domain containing 4C(C2cd4c) | 0,003273898 |
| **C77080** | expressed sequence C77080(C77080) | 0,001282304 |
| **Car8** | carbonic anhydrase 8(Car8) | 0,001173458 |
| **Cartpt** | CART prepropeptide(Cartpt) | 0,037712862 |
| **Casq2** | calsequestrin 2(Casq2) | 0,00014535 |
| **Catsperg2** | catsper channel auxiliary subunit gamma 2(Catsperg2) | 1,90518E-10 |
| **Cav3** | caveolin 3(Cav3) | 0,041711219 |
| **Ccnb1ip1** | cyclin B1 interacting protein 1(Ccnb1ip1) | 0,003848558 |
| **Cd14** | CD14 antigen(Cd14) | 0,000374514 |
| **Cd209b** | CD209b antigen(Cd209b) | 0,042559733 |
| **Cd209g** | CD209g antigen(Cd209g) | 0,042785929 |
| **Cd244** | CD244 natural killer cell receptor 2B4(Cd244) | 0,013998529 |
| **Cdh23** | cadherin 23 (otocadherin)(Cdh23) | 0,044266118 |
| **Cdh24** | cadherin-like 24(Cdh24) | 2,29496E-33 |
| **Ceacam15** | carcinoembryonic antigen-related cell adhesion molecule 15(Ceacam15) | 0,022321421 |
| **Cebpa** | CCAAT/enhancer binding protein (C/EBP), alpha(Cebpa) | 0,018975288 |
| **Cgref1** | cell growth regulator with EF hand domain 1(Cgref1) | 0,026222006 |
| **Cib2** | calcium and integrin binding family member 2(Cib2) | 2,7864E-05 |
| **Clcnka** | chloride channel, voltage-sensitive Ka(Clcnka) | 0,0009127 |
| **Cldn10** | claudin 10(Cldn10) | 3,44589E-10 |
| **Clec11a** | C-type lectin domain family 11, member a(Clec11a) | 0,025691417 |
| **Cmbl** | carboxymethylenebutenolidase-like (Pseudomonas)(Cmbl) | 0,04339448 |
| **Col11a1** | collagen, type XI, alpha 1(Col11a1) | 0,020643276 |
| **Col16a1** | collagen, type XVI, alpha 1(Col16a1) | 0,023948264 |
| **Col9a2** | collagen, type IX, alpha 2(Col9a2) | 0,009065738 |
| **Colgalt2** | collagen beta(1-O)galactosyltransferase 2(Colgalt2) | 1,89405E-12 |
| **Coq8b** | coenzyme Q8B(Coq8b) | 4,28282E-05 |
| **Cox14** | cytochrome c oxidase assembly protein 14(Cox14) | 0,000252078 |
| **Cox16** | cytochrome c oxidase assembly protein 16(Cox16) | 0,02205668 |
| **Cox18** | cytochrome c oxidase assembly protein 18(Cox18) | 0,010310937 |
| **Cox8a** | cytochrome c oxidase subunit VIIIa(Cox8a) | 0,000273249 |
| **Cplx1** | complexin 1(Cplx1) | 7,7592E-12 |
| **Crhr1** | corticotropin releasing hormone receptor 1(Crhr1) | 0,0212643 |
| **Csf3r** | colony stimulating factor 3 receptor (granulocyte)(Csf3r) | 0,014773473 |
| **Cst6** | cystatin E/M(Cst6) | 0,045690563 |
| **Ctu1** | cytosolic thiouridylase subunit 1(Ctu1) | 4,94178E-05 |
| **Cyp2j13** | cytochrome P450, family 2, subfamily j, polypeptide 13(Cyp2j13) | 9,64977E-05 |
| **Cyp2j9** | cytochrome P450, family 2, subfamily j, polypeptide 9(Cyp2j9) | 3,6628E-07 |
| **Cyp4f14** | cytochrome P450, family 4, subfamily f, polypeptide 14(Cyp4f14) | 3,50098E-05 |
| **Cyr61** | cysteine rich protein 61(Cyr61) | 0,003444989 |
| **D8Ertd738e** | DNA segment, Chr 8, ERATO Doi 738, expressed(D8Ertd738e) | 2,54235E-06 |
| **Dapl1** | death associated protein-like 1(Dapl1) | 0,034745775 |

#### Additional File 6: Table S2

| | | |
|---|---|---|
| **Derl3** | Der1-like domain family, member 3(Derl3) | 0,028116083 |
| **Dgcr6** | DiGeorge syndrome critical region gene 6(Dgcr6) | 0,000398309 |
| **Dhrs3** | dehydrogenase/reductase (SDR family) member 3(Dhrs3) | 1,24633E-06 |
| **Diaph3** | diaphanous related formin 3(Diaph3) | 0,011686966 |
| **Dmpk** | dystrophia myotonica-protein kinase(Dmpk) | 0,004617504 |
| **Dnah14** | dynein, axonemal, heavy chain 14(Dnah14) | 5,02964E-15 |
| **Draxin** | dorsal inhibitory axon guidance protein(Draxin) | 0,011359434 |
| **Emc9** | ER membrane protein complex subunit 9(Emc9) | 0,006178541 |
| **Eno1b** | enolase 1B, retrotransposed(Eno1b) | 1,66114E-17 |
| **Epb41l4a** | erythrocyte membrane protein band 4.1 like 4a(Epb41l4a) | 0,005523429 |
| **Ephx2** | epoxide hydrolase 2, cytoplasmic(Ephx2) | 4,05297E-05 |
| **Ethe1** | ethylmalonic encephalopathy 1(Ethe1) | 0,005413669 |
| **Fahd2a** | fumarylacetoacetate hydrolase domain containing 2A(Fahd2a) | 0,001626607 |
| **Fars2** | phenylalanine-tRNA synthetase 2 (mitochondrial)(Fars2) | 0,000286633 |
| **Fau** | Finkel-Biskis-Reilly murine sarcoma virus (FBR-MuSV) ubiquitously expressed (fox derived)(Fau) | 7,54284E-05 |
| **Fbxo2** | F-box protein 2(Fbxo2) | 5,79723E-06 |
| **Fbxo6** | F-box protein 6(Fbxo6) | 3,13747E-06 |
| **Fcgr1** | Fc receptor, IgG, high affinity I(Fcgr1) | 0,001805163 |
| **Fchsd1** | FCH and double SH3 domains 1(Fchsd1) | 0,028000124 |
| **Fgd2** | FYVE, RhoGEF and PH domain containing 2(Fgd2) | 0,007298807 |
| **Filip1l** | filamin A interacting protein 1-like(Filip1l) | 0,001506306 |
| **Foxo6** | forkhead box O6(Foxo6) | 0,040219376 |
| **Frmd3** | FERM domain containing 3(Frmd3) | 0,02462044 |
| **Fxyd1** | FXYD domain-containing ion transport regulator 1(Fxyd1) | 0,048938601 |
| **Fxyd6** | FXYD domain-containing ion transport regulator 6(Fxyd6) | 0,04378888 |
| **Fzd1** | frizzled class receptor 1(Fzd1) | 0,048910076 |
| **Gabra2** | gamma-aminobutyric acid (GABA) A receptor, subunit alpha 2(Gabra2) | 0,002552168 |
| **Gadd45b** | growth arrest and DNA-damage-inducible 45 beta(Gadd45b) | 0,020103162 |
| **Gal** | galanin(Gal) | 0,045213367 |
| **Gbp2b** | guanylate binding protein 2b(Gbp2b) | 1,87894E-07 |
| **Ghsr** | growth hormone secretagogue receptor(Ghsr) | 0,000278679 |
| **Gimap6** | GTPase, IMAP family member 6(Gimap6) | 0,029150268 |
| **Gipc1** | GIPC PDZ domain containing family, member 1(Gipc1) | 2,38641E-05 |
| **Gng13** | guanine nucleotide binding protein (G protein), gamma 13(Gng13) | 5,02712E-09 |
| **Gon4l** | gon-4-like (C.elegans)(Gon4l) | 8,53386E-13 |
| **Gpat2** | glycerol-3-phosphate acyltransferase 2, mitochondrial(Gpat2) | 0,005660112 |
| **Gpatch11** | G patch domain containing 11(Gpatch11) | 0,000160568 |
| **Gpld1** | glycosylphosphatidylinositol specific phospholipase D1(Gpld1) | 0,000730313 |
| **Gpx3** | glutathione peroxidase 3(Gpx3) | 0,009403913 |
| **Grap** | GRB2-related adaptor protein(Grap) | 0,000110831 |
| **Grem2** | gremlin 2, DAN family BMP antagonist(Grem2) | 1,27291E-05 |
| **Grk4** | G protein-coupled receptor kinase 4(Grk4) | 0,018519885 |
| **Gsta4** | glutathione S-transferase, alpha 4(Gsta4) | 7,05251E-07 |
| **Gstp1** | glutathione S-transferase, pi 1(Gstp1) | 0,004849194 |
| **Gucy2c** | guanylate cyclase 2c(Gucy2c) | 0,008990065 |
| **H2-Q2** | histocompatibility 2, Q region locus 2(H2-Q2) | 0,047318323 |
| **H2-T23** | histocompatibility 2, T region locus 23(H2-T23) | 0,044740374 |
| **H2-T24** | histocompatibility 2, T region locus 24(H2-T24) | 0,036180825 |
| **Hacd1** | 3-hydroxyacyl-CoA dehydratase 1(Hacd1) | 0,003737597 |
| **Hap1** | huntingtin-associated protein 1(Hap1) | 0,009139401 |
| **Haus4** | HAUS augmin-like complex, subunit 4(Haus4) | 6,24157E-08 |
| **Hddc3** | HD domain containing 3(Hddc3) | 0,002322164 |
| **Hebp2** | heme binding protein 2(Hebp2) | 0,014896141 |
| **Hist1h2ba** | histone cluster 1, H2ba(Hist1h2ba) | 0,00024879 |
| **Hist2h2bb** | histone cluster 2, H2bb(Hist2h2bb) | 1,17483E-12 |
| **Hmgcs2** | 3-hydroxy-3-methylglutaryl-Coenzyme A synthase 2(Hmgcs2) | 0,002194486 |
| **Hn1** | hematological and neurological expressed sequence 1(Hn1) | 3,54789E-07 |
| **Hnf1b** | HNF1 homeobox B(Hnf1b) | 0,000123622 |
| **Hpn** | hepsin(Hpn) | 0,016575122 |
| **Hrh2** | histamine receptor H2(Hrh2) | 0,01214204 |
| **Hspb7** | heat shock protein family, member 7 (cardiovascular)(Hspb7) | 0,039219681 |
| **Htra3** | HtrA serine peptidase 3(Htra3) | 0,047713851 |
| **Ifi202b** | interferon activated gene 202B(Ifi202b) | 5,78911E-09 |
| **Igfbp3** | insulin-like growth factor binding protein 3(Igfbp3) | 0,010686063 |
| **Igfbp6** | insulin-like growth factor binding protein 6(Igfbp6) | 0,006690813 |
| **Igsf21** | immunoglobulin superfamily, member 21(Igsf21) | 0,001969598 |
| **Il10ra** | interleukin 10 receptor, alpha(Il10ra) | 0,000138811 |
| **Inpp4a** | inositol polyphosphate-4-phosphatase, type I(Inpp4a) | 4,28282E-05 |
| **Invs** | inversin(Invs) | 0,003682237 |
| **Irf5** | interferon regulatory factor 5(Irf5) | 0,006063789 |
| **Itih3** | inter-alpha trypsin inhibitor, heavy chain 3(Itih3) | 8,19695E-05 |
| **Kcne1l** | potassium voltage-gated channel, Isk-related family, member 1-like, pseudogene(Kcne1l) | 0,035393066 |
| **Kcnh6** | potassium voltage-gated channel, subfamily H (eag-related), member 6(Kcnh6) | 0,003501711 |
| **Kcnj16** | potassium inwardly-rectifying channel, subfamily J, member 16(Kcnj16) | 0,000245359 |
| **Klhl33** | kelch-like 33(Klhl33) | 0,023876645 |
| **Klrg1** | killer cell lectin-like receptor subfamily G, member 1(Klrg1) | 0,000290116 |
| **Kptn** | kaptin(Kptn) | 4,48332E-11 |
| **Lcat** | lecithin cholesterol acyltransferase(Lcat) | 0,00115684 |
| **Ldhd** | lactate dehydrogenase D(Ldhd) | 8,09733E-09 |

#### Additional File 6: Table S2

| | | |
|---|---|---|
| **Lgi4** | leucine-rich repeat LGI family, member 4(Lgi4) | 9,93698E-12 |
| **Limk1** | LIM-domain containing, protein kinase(Limk1) | 0,000431861 |
| **Lims2** | LIM and senescent cell antigen like domains 2(Lims2) | 0,029705811 |
| **Lpl** | lipoprotein lipase(Lpl) | 0,024385469 |
| **Lrrc27** | leucine rich repeat containing 27(Lrrc27) | 0,000137807 |
| **Lsm3** | LSM3 homolog, U6 small nuclear RNA and mRNA degradation associated(Lsm3) | 0,001412329 |
| **Lsp1** | lymphocyte specific 1(Lsp1) | 0,000236691 |
| **Lst1** | leukocyte specific transcript 1(Lst1) | 0,003984899 |
| **Ly6h** | lymphocyte antigen 6 complex, locus H(Ly6h) | 0,033625344 |
| **Ly86** | lymphocyte antigen 86(Ly86) | 0,016535425 |
| **Lyplal1** | lysophospholipase-like 1(Lyplal1) | 1,0244E-14 |
| **Mad2l2** | MAD2 mitotic arrest deficient-like 2(Mad2l2) | 7,67117E-06 |
| **Mael** | maelstrom spermatogenic transposon silencer(Mael) | 0,035363452 |
| **March3** | membrane-associated ring finger (C3HC4) 3(March3) | 0,045087706 |
| **Mdk** | midkine(Mdk) | 0,014512013 |
| **Mien1** | migration and invasion enhancer 1(Mien1) | 3,30779E-10 |
| **Mill2** | MHC I like leukocyte 2(Mill2) | 0,000506448 |
| **Mme** | membrane metallo endopeptidase(Mme) | 0,001023525 |
| **Mrap2** | melanocortin 2 receptor accessory protein 2(Mrap2) | 0,005085478 |
| **Mroh5** | maestro heat-like repeat family member 5(Mroh5) | 0,034391061 |
| **Mroh7** | maestro heat-like repeat family member 7(Mroh7) | 0,010174898 |
| **Mrpl34** | mitochondrial ribosomal protein L34(Mrpl34) | 0,029340219 |
| **Mt3** | metallothionein 3(Mt3) | 7,87719E-06 |
| **Mttp** | microsomal triglyceride transfer protein(Mttp) | 1,09827E-08 |
| **Mup5** | major urinary protein 5(Mup5) | 0,020290342 |
| **Mxra7** | matrix-remodelling associated 7(Mxra7) | 0,009825154 |
| **Myh6** | myosin, heavy polypeptide 6, cardiac muscle, alpha(Myh6) | 2,20426E-13 |
| **Myh8** | myosin, heavy polypeptide 8, skeletal muscle, perinatal(Myh8) | 2,90074E-09 |
| **Myl1** | myosin, light polypeptide 1(Myl1) | 0,002327962 |
| **Nat8f5** | N-acetyltransferase 8 (GCN5-related) family member 5(Nat8f5) | 0,023403006 |
| **Ndn** | necdin(Ndn) | 8,76444E-08 |
| **Ndrg2** | N-myc downstream regulated gene 2(Ndrg2) | 1,20517E-15 |
| **Nedd8** | neural precursor cell expressed, developmentally down-regulated gene 8(Nedd8) | 1,03678E-05 |
| **Ngdn** | neuroguidin, EIF4E binding protein(Ngdn) | 4,42626E-06 |
| **Nkain4** | Na+/K+ transporting ATPase interacting 4(Nkain4) | 2,32098E-05 |
| **Nnat** | neuronatin(Nnat) | 0,032092937 |
| **Npas1** | neuronal PAS domain protein 1(Npas1) | 0,003227579 |
| **Nr2f6** | nuclear receptor subfamily 2, group F, member 6(Nr2f6) | 0,02166182 |
| **Nrk** | Nik related kinase(Nrk) | 0,047417636 |
| **Nrsn1** | neurensin 1(Nrsn1) | 5,15932E-08 |
| **Ntn4** | netrin 4(Ntn4) | 8,98546E-05 |
| **Nudt2** | nudix (nucleoside diphosphate linked moiety X)-type motif 2(Nudt2) | 0,000620555 |
| **Ost4** | oligosaccharyltransferase complex subunit 4 (non-catalytic)(Ost4) | 9,38489E-05 |
| **Ostf1** | osteoclast stimulating factor 1(Ostf1) | 0,00035024 |
| **Oxtr** | oxytocin receptor(Oxtr) | 0,032103073 |
| **P4ha3** | procollagen-proline, 2-oxoglutarate 4-dioxygenase (proline 4-hydroxylase), alpha polypeptide III(P4ha3) | 0,015854643 |
| **Padi2** | peptidyl arginine deiminase, type II(Padi2) | 0,007437629 |
| **Palm3** | paralemmin 3(Palm3) | 0,006189613 |
| **Papss2** | 3'-phosphoadenosine 5'-phosphosulfate synthase 2(Papss2) | 0,005787804 |
| **Pcbd2** | pterin 4 alpha carbinolamine dehydratase/dimerization cofactor of hepatocyte nuclear factor 1 alpha (TCF1) 2(Pcbd2) | 0,007332987 |
| **Pcdhb3** | protocadherin beta 3(Pcdhb3) | 8,19695E-05 |
| **Pcdhb6** | protocadherin beta 6(Pcdhb6) | 0,001327989 |
| **Pcdhb7** | protocadherin beta 7(Pcdhb7) | 0,000432829 |
| **Pcdhb8** | protocadherin beta 8(Pcdhb8) | 6,62307E-06 |
| **Pcsk1n** | proprotein convertase subtilisin/kexin type 1 inhibitor(Pcsk1n) | 0,008338887 |
| **Pdzk1** | PDZ domain containing 1(Pdzk1) | 0,007782851 |
| **Pgbd1** | piggyBac transposable element derived 1(Pgbd1) | 0,016680571 |
| **Pigf** | phosphatidylinositol glycan anchor biosynthesis, class F(Pigf) | 0,005869219 |
| **Pin1** | protein (peptidyl-prolyl cis/trans isomerase) NIMA-interacting 1(Pin1) | 0,005918898 |
| **Pip5kl1** | phosphatidylinositol-4-phosphate 5-kinase-like 1(Pip5kl1) | 0,014731124 |
| **Pkp1** | plakophilin 1(Pkp1) | 0,019978324 |
| **Plscr2** | phospholipid scramblase 2(Plscr2) | 0,014896141 |
| **Plvap** | plasmalemma vesicle associated protein(Plvap) | 0,008338887 |
| **Podxl2** | podocalyxin-like 2(Podxl2) | 0,020259441 |
| **Ppp1r3g** | protein phosphatase 1, regulatory (inhibitor) subunit 3G(Ppp1r3g) | 0,014655538 |
| **Proca1** | protein interacting with cyclin A1(Proca1) | 5,37244E-08 |
| **Proser3** | proline and serine rich 3(Proser3) | 0,024183861 |
| **Prtn3** | proteinase 3(Prtn3) | 0,045690563 |
| **Psmb10** | proteasome (prosome, macropain) subunit, beta type 10(Psmb10) | 4,14628E-05 |
| **Psmb11** | proteasome (prosome, macropain) subunit, beta type, 11(Psmb11) | 9,03225E-16 |
| **Psmg4** | proteasome (prosome, macropain) assembly chaperone 4(Psmg4) | 0,000308357 |
| **Pth1r** | parathyroid hormone 1 receptor(Pth1r) | 0,012803732 |
| **Pvalb** | parvalbumin(Pvalb) | 0,009247559 |
| **Pxmp2** | peroxisomal membrane protein 2(Pxmp2) | 0,000309658 |
| **Pyroxd2** | pyridine nucleotide-disulphide oxidoreductase domain 2(Pyroxd2) | 0,000209316 |
| **Qsox1** | quiescin Q6 sulfhydryl oxidase 1(Qsox1) | 3,15507E-07 |
| **Rabac1** | Rab acceptor 1 (prenylated)(Rabac1) | 2,13046E-06 |
| **Ramp1** | receptor (calcitonin) activity modifying protein 1(Ramp1) | 0,000326559 |
| **Rdh13** | retinol dehydrogenase 13 (all-trans and 9-cis)(Rdh13) | 2,45414E-05 |

### Additional File 6: Table S2

| | | |
|---|---|---|
| **Rdm1** | RAD52 motif 1(Rdm1) | 0,00021784 |
| **Reg3b** | regenerating islet-derived 3 beta(Reg3b) | 0,000705532 |
| **Resp18** | regulated endocrine-specific protein 18(Resp18) | 0,038218746 |
| **Rgcc** | regulator of cell cycle(Rgcc) | 0,027545958 |
| **Rgsl1** | regulator of G-protein signaling like 1(Rgsl1) | 0,012803732 |
| **Rnase1** | ribonuclease, RNase A family, 1 (pancreatic)(Rnase1) | 9,11616E-16 |
| **Rnaseh2c** | ribonuclease H2, subunit C(Rnaseh2c) | 0,003719927 |
| **Rnd2** | Rho family GTPase 2(Rnd2) | 0,000598745 |
| **Rnf223** | ring finger 223(Rnf223) | 0,012886903 |
| **Rnls** | renalase, FAD-dependent amine oxidase(Rnls) | 0,030187967 |
| **Romo1** | reactive oxygen species modulator 1(Romo1) | 0,002521045 |
| **Rpa2** | replication protein A2(Rpa2) | 2,32898E-05 |
| **Rplp1** | ribosomal protein, large, P1(Rplp1) | 0,001762362 |
| **Rxfp3** | relaxin family peptide receptor 3(Rxfp3) | 0,025262545 |
| **S100b** | S100 protein, beta polypeptide, neural(S100b) | 6,62307E-06 |
| **Sag** | S-antigen, retina and pineal gland (arrestin)(Sag) | 0,005760837 |
| **Samd11** | sterile alpha motif domain containing 11(Samd11) | 0,024128788 |
| **Scrg1** | scrapie responsive gene 1(Scrg1) | 0,029641461 |
| **Sdhc** | succinate dehydrogenase complex, subunit C, integral membrane protein(Sdhc) | 3,23692E-11 |
| **Selenbp2** | selenium binding protein 2(Selenbp2) | 7,60827E-10 |
| **Sema4g** | sema domain, immunoglobulin domain (Ig),(semaphorin) 4G(Sema4g) | 0,001488735 |
| **Serf1** | small EDRK-rich factor 1(Serf1) | 2,01762E-06 |
| **Serpina3f** | serine (or cysteine) peptidase inhibitor, clade A, member 3F(Serpina3f) | 0,006759948 |
| **Serpind1** | serine (or cysteine) peptidase inhibitor, clade D, member 1(Serpind1) | 0,002291179 |
| **Sgpp2** | sphingosine-1-phosphate phosphotase 2(Sgpp2) | 0,047318323 |
| **Sgsm2** | small G protein signaling modulator 2(Sgsm2) | 1,99277E-06 |
| **Sh2b2** | SH2B adaptor protein 2(Sh2b2) | 0,005441559 |
| **Shh** | sonic hedgehog(Shh) | 0,044428741 |
| **Slamf8** | SLAM family member 8(Slamf8) | 5,06196E-07 |
| **Slc13a5** | solute carrier family 13 (sodium-dependent citrate transporter), member 5(Slc13a5) | 0,000112413 |
| **Slc15a3** | solute carrier family 15, member 3(Slc15a3) | 0,007430117 |
| **Slc16a1** | solute carrier family 16 (monocarboxylic acid transporters), member 1(Slc16a1) | 0,020817532 |
| **Slc1a6** | solute carrier family 1 (high affinity aspartate/glutamate transporter), member 6(Slc1a6) | 0,030729958 |
| **Slc25a31** | solute carrier family 25 (mitochondrial carrier; adenine nucleotide translocator), member 31(Slc25a31) | 0,018115939 |
| **Slc2a5** | solute carrier family 2 (facilitated glucose transporter), member 5(Slc2a5) | 0,000676652 |
| **Slc30a2** | solute carrier family 30 (zinc transporter), member 2(Slc30a2) | 0,024438813 |
| **Slc36a2** | solute carrier family 36 (proton/amino acid symporter), member 2(Slc36a2) | 0,044726931 |
| **Slc7a7** | solute carrier family 7 (cationic amino acid transporter, y+ system), member 7(Slc7a7) | 0,04830052 |
| **Smdt1** | single-pass membrane protein with aspartate rich tail 1(Smdt1) | 1,26407E-05 |
| **Smim1** | small integral membrane protein 1(Smim1) | 0,002788164 |
| **Smyd1** | SET and MYND domain containing 1(Smyd1) | 0,001871246 |
| **Sncb** | synuclein, beta(Sncb) | 0,004661403 |
| **Snrnp25** | small nuclear ribonucleoprotein 25 (U11/U12)(Snrnp25) | 2,60939E-05 |
| **Sp100** | nuclear antigen Sp100(Sp100) | 0,033739435 |
| **Sparc** | secreted acidic cysteine rich glycoprotein(Sparc) | 0,003889208 |
| **Spink10** | serine peptidase inhibitor, Kazal type 10(Spink10) | 0,035313138 |
| **Stk10** | serine/threonine kinase 10(Stk10) | 0,019988929 |
| **Strc** | stereocilin(Strc) | 0,007385701 |
| **Sumf2** | sulfatase modifying factor 2(Sumf2) | 0,003052813 |
| **Susd2** | sushi domain containing 2(Susd2) | 0,003052813 |
| **Susd5** | sushi domain containing 5(Susd5) | 0,006459583 |
| **Swt1** | SWT1 RNA endoribonuclease homolog (S. cerevisiae)(Swt1) | 0,002117681 |
| **Sytl1** | synaptotagmin-like 1(Sytl1) | 0,003088137 |
| **Tbc1d9b** | TBC1 domain family, member 9B(Tbc1d9b) | 4,33403E-20 |
| **Tcap** | titin-cap(Tcap) | 0,000401008 |
| **Tgfb1** | transforming growth factor, beta 1(Tgfb1) | 0,007728873 |
| **Timp4** | tissue inhibitor of metalloproteinase 4(Timp4) | 9,3705E-06 |
| **Tlcd2** | TLC domain containing 2(Tlcd2) | 0,022415558 |
| **Tlr1** | toll-like receptor 1(Tlr1) | 0,0128052 |
| **Tlr6** | toll-like receptor 6(Tlr6) | 0,022460838 |
| **Tmem132e** | transmembrane protein 132E(Tmem132e) | 0,03038241 |
| **Tmem173** | transmembrane protein 173(Tmem173) | 0,000334003 |
| **Tmem176a** | transmembrane protein 176A(Tmem176a) | 0,038218746 |
| **Tmem30c** | transmembrane protein 30C(Tmem30c) | 0,014993972 |
| **Tmem74b** | transmembrane protein 74B(Tmem74b) | 0,012898105 |
| **Tmod3** | tropomodulin 3(Tmod3) | 0,00275412 |
| **Tmod4** | tropomodulin 4(Tmod4) | 0,000122148 |
| **Tmsb10** | thymosin, beta 10(Tmsb10) | 0,00167863 |
| **Tnfaip8** | tumor necrosis factor, alpha-induced protein 8(Tnfaip8) | 0,000483409 |
| **Tpm2** | tropomyosin 2, beta(Tpm2) | 0,007924549 |
| **Trabd2b** | TraB domain containing 2B(Trabd2b) | 0,019580787 |
| **Traf3ip3** | TRAF3 interacting protein 3(Traf3ip3) | 0,015136822 |
| **Trappc6a** | trafficking protein particle complex 6A(Trappc6a) | 0,000178642 |
| **Tsks** | testis-specific serine kinase substrate(Tsks) | 0,007782851 |
| **Tspan17** | tetraspanin 17(Tspan17) | 8,61665E-05 |
| **Ttc9b** | tetratricopeptide repeat domain 9B(Ttc9b) | 0,017345165 |
| **Ttk** | Ttk protein kinase(Ttk) | 0,015098252 |
| **Tuba1c** | tubulin, alpha 1C(Tuba1c) | 0,0254744 |
| **Tusc1** | tumor suppressor candidate 1(Tusc1) | 0,040833696 |

#### Additional File 6: Table S2

| | | |
|---|---|---|
| **Tyrobp** | TYRO protein tyrosine kinase binding protein(Tyrobp) | 0,000181857 |
| **Uba5** | ubiquitin-like modifier activating enzyme 5(Uba5) | 6,89074E-24 |
| **Unc119** | unc-119 lipid binding chaperone(Unc119) | 0,000251962 |
| **Uqcc2** | ubiquinol-cytochrome c reductase complex assembly factor 2(Uqcc2) | 0,00010448 |
| **Uqcc3** | ubiquinol-cytochrome c reductase complex assembly factor 3(Uqcc3) | 1,79836E-05 |
| **Uqcr11** | ubiquinol-cytochrome c reductase, complex III subunit XI(Uqcr11) | 0,000374312 |
| **Uqcrq** | ubiquinol-cytochrome c reductase, complex III subunit VII(Uqcrq) | 0,001631318 |
| **Uty** | ubiquitously transcribed tetratricopeptide repeat gene, Y chromosome(Uty) | 0,000835085 |
| **Wbscr27** | Williams Beuren syndrome chromosome region 27 (human)(Wbscr27) | 0,01110656 |
| **Wdfy1** | WD repeat and FYVE domain containing 1(Wdfy1) | 0,000122974 |
| **Wnt4** | wingless-type MMTV integration site family, member 4(Wnt4) | 0,009155246 |
| **Wnt7a** | wingless-type MMTV integration site family, member 7A(Wnt7a) | 0,01214204 |

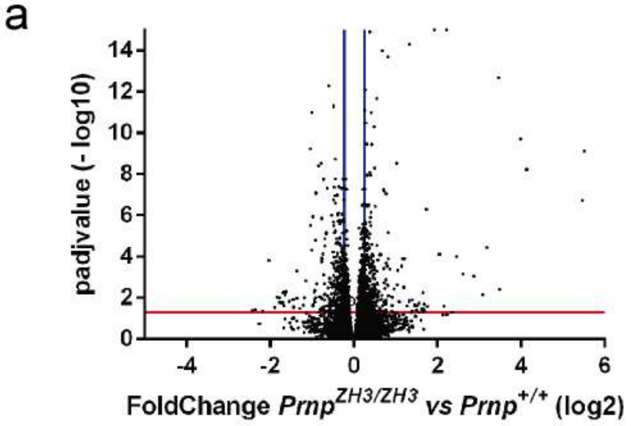


b

**Downregulated genes**

| Pathway name | Gene Ratio | p-value |
|---|---|---|
| MECP2 regulates neuronal receptors and channels | 10/32 | 1,06E-07 |
| Nuclear Receptor transcription pathway | 10/86 | 4,79E-04 |
| Potassium Channels | 11/107 | 6,99E-04 |
| Protein-protein interactions at synapses | 10/93 | 8,65E-04 |
| Neuronal System | 28/487 | 0,002 |

**Upregulated genes**

| Pathway name | Gene Ratio | p-value |
|---|---|---|
| Respiratory electron transport, ATP synthesis by chemiosmotic coupling, and heat production by uncoupling proteins | 10/153 | 0,006 |
| The citric acid (TCA) cycle and respiratory electron transport | 13/235 | 0,007 |
| Cellular response to chemical stress | 11/208 | 0,017 |

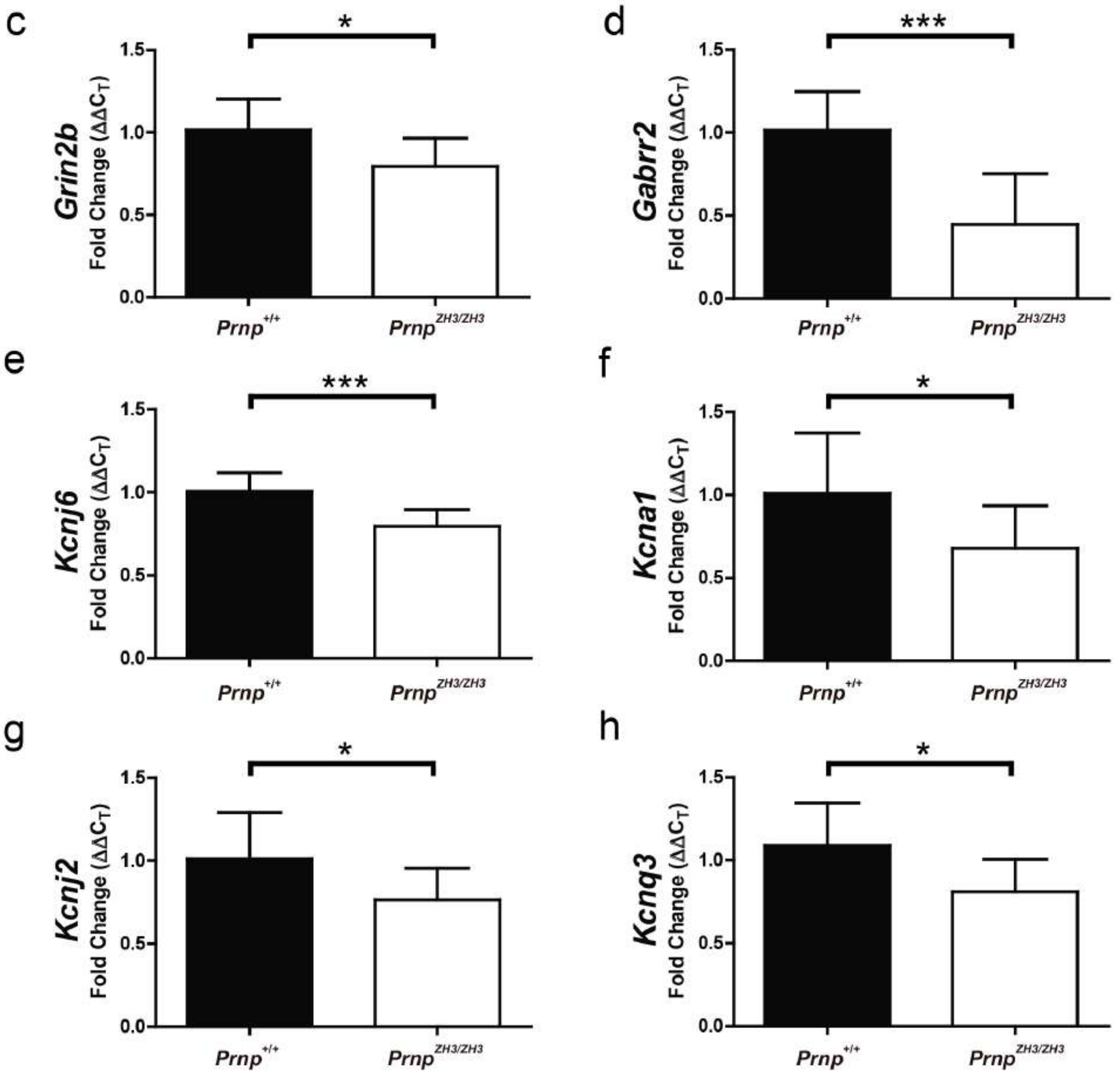


Additional File 7: Figure S5
